# Sepsis-associated Pathways Segregate Cancer Groups

**DOI:** 10.1101/635243

**Authors:** Himanshu Tripathi, Samanwoy Mukhopadhyay, Saroj Kant Mohapatra

## Abstract

**Background:** Sepsis and cancer are both leading causes of death, and occurrence of any one, increases the likelihood of the other. While cancer patients are susceptible to sepsis, survivors of sepsis are also susceptible to develop certain cancers. This mutual dependence for susceptibility suggests shared biology between the two disease categories. Earlier analysis had revealed cancer-related pathway to be up-regulated in Septic Shock (SS), an advanced stage of sepsis. This has motivated a more comprehensive comparison of the transcriptomes of SS and cancer.

**Methods:** Gene Set Enrichment Analysis was performed to detect the pathways enriched in SS and cancer. Thereafter, hierarchical clustering was applied to identify relative segregation of 17 cancer types in to two groups *vis-a-vis* SS. Biological significance of the selected pathways was explored by network analysis. Clinical significance of the pathways was tested by survival analysis. A robust classifier of cancer groups was developed based on machine learning.

**Results:** A total of 66 pathways were observed to be enriched in both SS and cancer. However, clustering segregated cancer types into two categories based on the direction of transcriptomic change. In general, there was up-regulation in SS and one group of cancer (termed Sepsis-Like Cancers, or SLC), but not in other cancers. SLC group mainly consisted of malignancies of the gastrointestinal tract (head and neck, oesophagus, stomach, liver and biliary system) often associated with infection. Machine learning classifier successfully segregated the two cancer groups with high accuracy (> 98%). Additionally, pathway up-regulation was observed to be associated with survival in the SLC group of cancers.

**Conclusion:** Transcriptome-based systems biology approach segregates cancer into two groups (SLC and CA) based on similarity with SS. Host response to infection plays a key role in pathogenesis of SS and SLC. However, we hypothesize that some component of the host response is protective in both SS and SLC.

## Background

Sepsis is a potentially life-threatening complication caused by dysregulated host response to infection, often leading to organ failure and death. Estimated global burden of sepsis is more than 48.9 million people in 2017 with 11 million deaths [1]. Septic shock is the advanced stage of sepsis with metabolic dysregulation and uncontrolled hypotension. Several epidemiological studies have linked sepsis and cancer [2, 3]. Liu *et al.* [2] conducted an association study between sepsis and ensuing risk of cancer in elderly adult population of the United States, and observed that sepsis is significantly associated with increased risk for many cancers including chronic myeloid leukemia, myelodysplastic syndrome, acute myeloid leukemia, cancers of the cervix, liver, lung, rectum, colon. Another association study revealed 2.5 fold increased risk of sepsis in survivors of cancer in community-dwelling adults (the risk is increased up to 10 times in hospitalized cancer patients) [3]. Co-occurrence of cancer with sepsis is associated with higher mortality than sepsis alone without cancer [4]. On the other hand, sepsis is a common cause of death in critically ill patients with cancer, with high ICU and hospital mortality [5–7]. Interestingly, mortality due to sepsis varies widely from 42 to 82% across cancer tissue types [8], suggesting varying likelihood of survival of patients suffering from different cancers.

This study is motivated by multiple shared features of septic shock (SS) and cancer. There is co-occurrence of the two entities, with synergistic effect on mortality. Both are associated with inflammation at some stage of the disease. Inflammation is well understood to promote malignant growth with participation of diverse immune cells and molecules, such as, cytokines [9]. Similarly, sepsis is understood as a non-resolving inflammatory response to infection that leads to organ dysfunction [10]. Both the diseases are associated with anaerobic metabolism with lactic acidosis being a hallmark of septic shock [9, 11]. In our earlier work, we have observed a cancer associated pathway to be significantly up-regulated in septic shock [12]. Additionally, there are previous reports on shared molecular changes in sepsis and cancer. Bergenfelz *et al.* [13] reported that Wnt5a induces immunosuppressive phenotype of macrophages in sepsis and breast cancer patients. HMGB1, a key late inflammatory mediator of systemic inflammatory response syndrome associated with bacterial sepsis, is also implicated in tumorigenesis and disease progression [14]. Muscle wasting - observed in patients with cancer, severe injury and sepsis - is associated with increased expression of several genes, particularly transcription factors and nuclear cofactors, regulating different proteolytic pathways [15]. Methodologically, all of these studies employed gene-level analysis, i.e., considering gene as the functional unit. On the other hand, a gene set or pathway represents coordinated molecular activity and represents a higher-order functional unit in a tissue or cell. Pathway-level analysis allows detection of a cumulative signal that is not accessible at the gene-level. To our knowledge, there is no report in literature on pathway-level comparison between sepsis and cancer. In the present study, we have performed unbiased analysis of SS and cancer datasets to discover shared patterns of pathway-level transcriptional alteration underlying the two illnesses.

## Methods

### Gene expression data

#### TCGA data

Gene expression data as FPKM (Fragments Per Kilobase of transcript per Million mapped reads) values were retrieved from The Cancer Genome Atlas (TCGA) database (https://portal.gdc.cancer.gov/) on July 5, 2018 for 17 different human cancers i.e., bladder urothelial carcinoma (BLCA), breast invasive carcinoma (BRCA), cholangiocarcinoma (CHOL), colon adenocarcinoma (COAD), esophageal carcinoma (ESCA), head and neck squamous cell carcinoma (HNSC), kidney chromophobe (KICH), kidney renal clear cell carcinoma (KIRC), kidney renal papillary cell carcinoma (KIRP), liver hepatocellular carcinoma (LIHC), lung adenocarcinoma (LUAD), lung squamous cell carcinoma (LUSC), prostate adenocarcinoma (PRAD), rectum adenocarcinoma (READ), stomach adenocarcinoma (STAD), thyroid carcinoma (THCA) and uterine corpus endometrial carcinoma (UCEC). For each cancer type, TCGA project code was provided in the search field and RNA-seq data for paired samples (each pair consisting of tissue from tumour zone and tissue from adjacent unaffected zone in the same individual) were downloaded. In all, gene expression data were derived from 687 patients with cancer. Data were transformed to logarithmic scale (base 2).

#### GEO data

Six studies of septic shock (SS) were selected following the procedure described earlier [12]. Normalized gene expression data from the series matrix files were retrieved from NCBI Gene Expression Omnibus (GEO) database (https://www.ncbi.nlm.nih.gov/geo/) on April 10, 2019. Data were transformed to logarithmic scale (base 2). Expression intensity for each Entrez gene ID was calculated after removing duplicated probe sets. Genes common to all SS studies were included in the analysis. In all, gene expression data were derived from 445 patients with SS and 116 control subjects. Redundant samples (other than control and SS) were excluded from analysis.

Characteristics of 23 distinct data sets used in the current study (6 of SS and 17 of cancer) are listed in Table 1. Each data set consisted of transcriptome-wide expression data from a number of patients suffering from a single disease (SS or cancer). For SS, each data set consisted of blood transcriptome data. For cancer, each data set consisted of transcriptome data from a tissue. Control was defined as adjacent normal tissue for cancer (TCGA), and a healthy subject for SS (GEO).

**Table 1:** Characteristics (such as tissue of origin, source database, disease type, sample size of each study and details about the platform technology used to generate the data) of the 23 data sets (17 cancer + 6 septic shock) included in the analysis. Study_code refers to the code assigned by the source database (either TCGA or GEO) to the data set. TCGA stands for The Cancer Genome Atlas. GEO stands for NCBI Gene Expression Omnibus. Paired control refers to adjacent normal tissue in the same cancer patient. Technology for transcriptome assay is either sequencing (RNA-seq) or hybridization-based microarray (Affymetrix).

| Study_Code | Data source | Disease | Tissue | Number of samples | Paired control | Technology |
| --- | --- | --- | --- | --- | --- | --- |
| BLCA | TCGA | Cancer | Bladder | 38 | Yes | RNA-seq (Illumina) |
| BRCA | TCGA | Cancer | Breast | 220 | Yes | RNA-seq (Illumina) |
| CHOL | TCGA | Cancer | Gallbladder, liver and parts of biliary tract | 18 | Yes | RNA-seq (Illumina) |
| COAD | TCGA | Cancer | Colon and rectosigmoid junction | 82 | Yes | RNA-seq (Illumina) |
| ESCA | TCGA | Cancer | Esophagus | 16 | Yes | RNA-seq (Illumina) |
| HNSC | TCGA | Cancer | Base of tongue, floor of mouth, gum, hypo- and oro-pharynx, larynx, etc. | 86 | Yes | RNA-seq (Illumina) |
| KICH | TCGA | Cancer | Kidney | 48 | Yes | RNA-seq (Illumina) |
| KIRC | TCGA | Cancer | Kidney | 144 | Yes | RNA-seq (Illumina) |
| KIRP | TCGA | Cancer | Kidney | 64 | Yes | RNA-seq (Illumina) |
| LIHC | TCGA | Cancer | Liver and intrahepatic bile ducts | 100 | Yes | RNA-seq (Illumina) |
| LUAD | TCGA | Cancer | Bronchus and lung | 114 | Yes | RNA-seq (Illumina) |
| LUSC | TCGA | Cancer | Bronchus and lung | 98 | Yes | RNA-seq (Illumina) |
| PRAD | TCGA | Cancer | Prostate gland | 104 | Yes | RNA-seq (Illumina) |
| READ | TCGA | Cancer | Rectum rectosigmoid junction | 20 | Yes | RNA-seq (Illumina) |
| STAD | TCGA | Cancer | Stomach | 62 | Yes | RNA-seq (Illumina) |
| THCA | TCGA | Cancer | Thyroid gland | 116 | Yes | RNA-seq (Illumina) |
| UCEC | TCGA | Cancer | Corpus uteri | 46 | Yes | RNA-seq (Illumina) |
| GSE4607 | GEO | Septic Shock | Whole Blood | 84 | No | Microarray (Affymetrix HGU 133 Plus 2.0) |
| GSE8121 | GEO | Septic Shock | Whole Blood | 75 | No | Microarray (Affymetrix HGU 133 Plus 2.0) |
| GSE9692 | GEO | Septic Shock | Whole Blood | 45 | No | Microarray (Affymetrix HGU 133 Plus 2.0) |
| GSE13904 | GEO | Septic Shock | Whole Blood | 124 | No | Microarray (Affymetrix HGU 133 Plus 2.0) |
| GSE26378 | GEO | Septic Shock | Whole Blood | 103 | No | Microarray (Affymetrix HGU 133 Plus 2.0) |
| GSE26440 | GEO | Septic Shock | Whole Blood | 130 | No | Microarray (Affymetrix HGU 133 Plus 2.0) |

### Pathway Enrichment Analysis

Pathway (gene set) enrichment analysis was performed using the algorithm previously described [12, 16]. Gene sets were defined based on pathways annotated at KEGG [17]. Any pathway with small number (10 or less) of genes was discarded from analysis. For each gene, t-statistic was computed to denote change in gene expression in case group compared to the control group. For each pathway, a score was calculated by weighted averaging (i.e., sum of the gene-level t-statistics divided by the square root of the number of genes in the pathway) of all gene-level t-statistic for the pathway. Significance of the observed pathway score was calculated by permutation testing performed in the following manner. In each permutation, the samples were randomly re-labelled as case and control, with calculation of a simulated pathway score. This was done 10000 times generating 10000 simulated values representing the null distribution of the pathway score. Deviation of the observed pathway score from the null distribution was quantified by the fraction of times that the simulated score was more extreme than the observed score. This result was assigned as permutation p-value of the observed pathway score. Pathway enrichment analysis was performed using code modified from the R function *gseattperm()* of the package **Category** [18].

### Cluster Analysis

Pathway scores of all 23 studies (6 SS + 17 cancer) were subjected to hierarchical clustering in the following manner. First, the Euclidean distance matrix was computed to capture the pair-wise dissimilarity among the studies. Thereafter, the distance matrix was subjected to agglomerative hierarchical cluster analysis, with “complete” linkage method. Distance matrix computation and cluster analysis were performed using the R functions *dist()* and *hclust()* respectively. Output of the cluster analysis was plotted as a dendrogram.

### Visualization of Pathway-level and Gene-level Expression Scores

A pathway was selected if it was significantly enriched (permutation p < 0.01) in 80% or more of the studies in one or both of the cancer groups (SLC, CA). Pathway score matrix was generated for all selected pathways across all 23 data sets. Heat map was generated with the R function *heatmap.2()* of the package **gplots** [19]. For each pathway, mean pathway score across the disease group was calculated separately. Boxplot of the pathway scores for the three disease groups (SS, SLC and CA) was drawn using R function *boxplot()*.

For generating gene-level heat map of individual pathway, the following steps were followed. First, average log-expression level was calculated for case group and control group separately. Log2(fold-change) or LFC was calculated by subtracting the average control value from the average case value. This was done for each gene. For each disease type (SS, SLC or CA), median LFC was calculated across the studies in that disease group. Combined heat map was generated based on the pathway genes and the three disease groups. Only those genes with similar LFC directionality in SLC and SS were included in visualization.

### Network analysis

Firstly, a background network was constructed by including all the pathways (KEGG) [17] with overlapping gene memberships, i.e., for a pathway to be included, it must share at least 5% of the total number of genes with another pathway. Pathways connected to less than 3 other pathways each were dropped from network analysis. In this network, each pathway was considered a node, and the edge between two nodes represented overlap between the two pathways. In this way, a network was constructed with 244 nodes and 5304 edges. Degree distribution of the network was calculated using the function *degree_distribution()* of the package **igraph** [20]. Plot of the degree distribution of the network was drawn. For each node in the network, degree and betweenness centrality were calculated with the R functions *degree()* and *betweenness()* respectively. In the network diagram, selected pathways were shown as nodes coloured in red. Box plot was drawn to show the difference of degrees between selected set and other pathways.

### Sample-level pathway score

Individual pathway scores for the cancer patients were computed in the following manner. For each subject, two genome-wide expression vectors were retrieved from TCGA: one for the tumour tissue (T) and another for the adjacent normal tissue (N). Normalized gene expression vector (E) was then calculated by subtracting the normal expression values from the tumour expression values (E = T - N). For each of the selected 66 pathways, a score (Z) was calculated as the weighted average of the expression of the genes in the pathway (i.e., sum of the individual gene expression scores divided by the square root of the number of genes in the pathway). This resulted in 66 pathway scores for each patient. These individual pathway scores were used for survival analysis.

### Machine learning

For the machine learning-based prediction, we retrieved additional transcriptome data for SLC and CA cancers from NCBI GEO. Six datasets containing 542 cancer subjects (of both SLC and CA) and 180 healthy control subjects were included (Additional File 7: Table S4.). The additional validation data sets were required because the TCGA data sets were used for feature selection (i.e., 66 pathways) and segregation of cancer groups (SLC and CA). For each cancer sample, pathway scores were generated by weighted averaging of the gene-level t-statistic between the case group and control group, (i.e., sum of the individual t-statistic divided by the square root of the number of genes in the pathway). The scores of the sixty-six pathways were used as input and the cancer group label (SLC or CA) as output in machine learning-based classification of cancer patients. Support vector machine (SVM) and neural net (NN) classifiers were implemented using R package **MLInterfaces** [21]. SVM was used through function call *Mlearn()* with learning function svmI. Neural Network was used through function call *MLearn()* with learning function nnetI and three nodes in the hidden layer, with weight decay set to 0.01. Confusion matrix from the classifier output of five-fold cross-validation was collected for calculation of misclassification rate (fraction of samples wrongly predicted by the algorithm) and accuracy.

### Survival Analysis

Based on the number of cases and available survival information, three SLC cancer types (HNSC, KIRC and LIHC) were selected for survival analysis. The following clinical metadata were collected as provided by TCGA: days (to death or last follow-up) and outcome (survivor or non-survivor). Patient-level (sample-level) pathway score was used for analysis. For each cancer type, the patients were divided into two groups (high-score and low-score) by applying the median pathway score as threshold. Survival plots were generated using the functions in the R package **survival** [22] as described here. First, a survival object was created applying the function *Surv()*, with day and outcome. The output, i.e., the fitted model, was passed to the function *survfit()* that created two survival curves, based on the level of pathway score (high or low). Survival plots were displayed using the function *ggsurvplot()* from the package **survminer** [23]. Significance of the difference between the survival plots of the two curves was calculated by the function *surv_pvalue()* from the same package.

### Code and data availability

All programming was done in the R programming language [24]. Data and R code are available at the following link: https://figshare.com/ (https://doi.org/10.6084/m9.figshare.8118413.v3, https://doi.org/10.6084/m9.figshare.8118647.v5). The whole data can be downloaded as a single zip file (tcnibmgdoc.zip). Upon uncompressing the zip, instructions for running the code and generating the figures of this manuscript are available in the file howto.pdf.

Selection of the transcriptome data sets and analysis workflow leading to the final list of 66 significant pathways are described in Fig. 1. The study characteristics of the data sets are listed in Table 1.

**Fig. 1:**
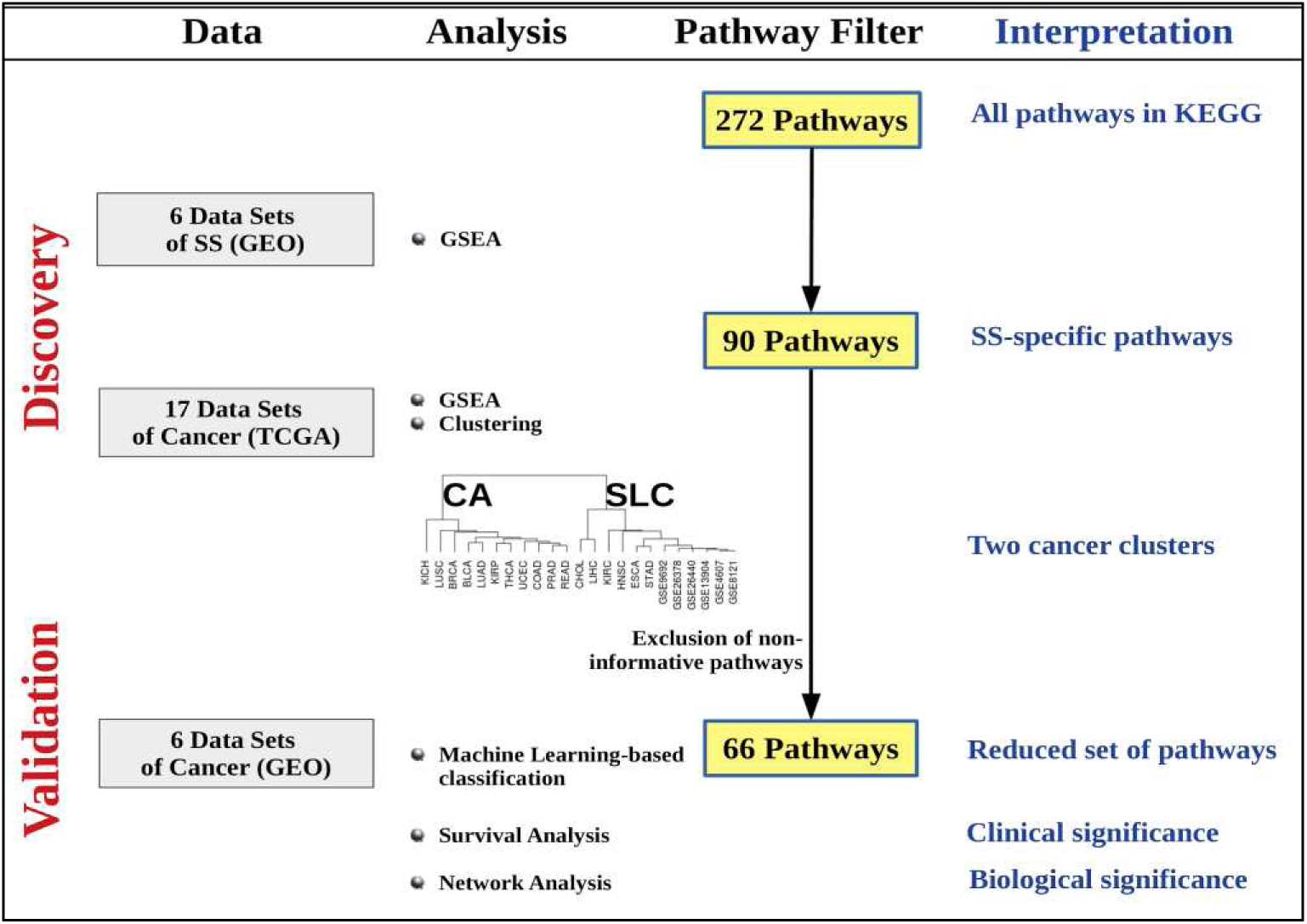
Selection of data and analysis workflow. Left-most column shows the source of data from GEO for SS and TCGA for cancer. Data for SS were applied first followed by data from TCGA. Next column describes the processing of the data that form a progressive filter for narrowing down the number of pathways of relevance. KEGG contains 272 pathways of which 90 were assigned significant pathway score by Gene Set Enrichment Analysis (details in the main text). The right-most column lists the interpretations made at each horizontal step.

## Results

The transcriptomic data for 23 studies (17 cancer and 6 SS) consisted of 687 patients with cancer and 445 with SS. The data were subjected to progressive analysis and pathway filtering (Fig. 1).

### Hierarchical clustering revealed two groups of cancers

At the outset, independent of our data, there were a total of 272 pathways in the KEGG database. Application of permutation-based testing of each pathway, revealed that 90 of these were significantly (p < 0.01) enriched in each of the 6 SS data sets, thus constituting a robust set of SS-specific pathways. Hierarchical clustering on combined cancer and sepsis data sets for these 90 pathways revealed two groups of cancers, with one group segregating with SS (Additional File 1: Figure S1). The 6 cancers (HNSC, ESCA, STAD, LIHC, CHOL, KIRC) belonging to the SS clade were termed Sepsis-Like Cancer (SLC). The other 11 cancers (BLCA, BRCA, COAD, KICH, KIRP, LUAD, LUSC, PRAD, READ, THCA and UCEC) formed the other clade and were termed the CA (Cancer Alone) group. In order to exclude those pathways that may be irrelevant to cancer biology, the 90 pathways were tested for enrichment in any or both of the cancer groups (i.e., significantly enriched in at least 80% of a group of cancer). 24 out of the 90 SS-specific pathways were observed not to be associated with any of the two cancer groups. Exclusion of these 24 non-significant pathways resulted in retention of 66 pathways significantly associated with both SS and cancer. Pathway score matrix of these 66 pathways across all diseases (including SS, SLC, CA) was visualized as a heat map (Fig. 2A). The sample dendrogram of this heat map recapitulated the segregation of the cancer groups as observed earlier with 90 pathways (Additional File 1: Figure S1). In general, the heat map demonstrated up-regulation of the pathways in SS and SLC. Further, for each pathway, mean score was calculated across all data sets in one disease group (SS, SLC or CA). Box plot of these scores clearly demonstrated the shared directionality in pathway dysregulation in SS and SLC, i.e., in both these conditions there was up-regulation of the pathway (Fig. 2B). Additionally, the average number of pathways up-regulated in a disease group was observed to be 63, 58 and 11 for SS, SLC and CA respectively (Additional File 4: Table S1), thus establishing general agreement in the direction of pathway transcriptional change between SS and SLC. The overall trend in pathway dysregulation was also reflected in the gene-level heat map for individual pathways (Additional File 2: Figure S2).

**Fig. 2:**
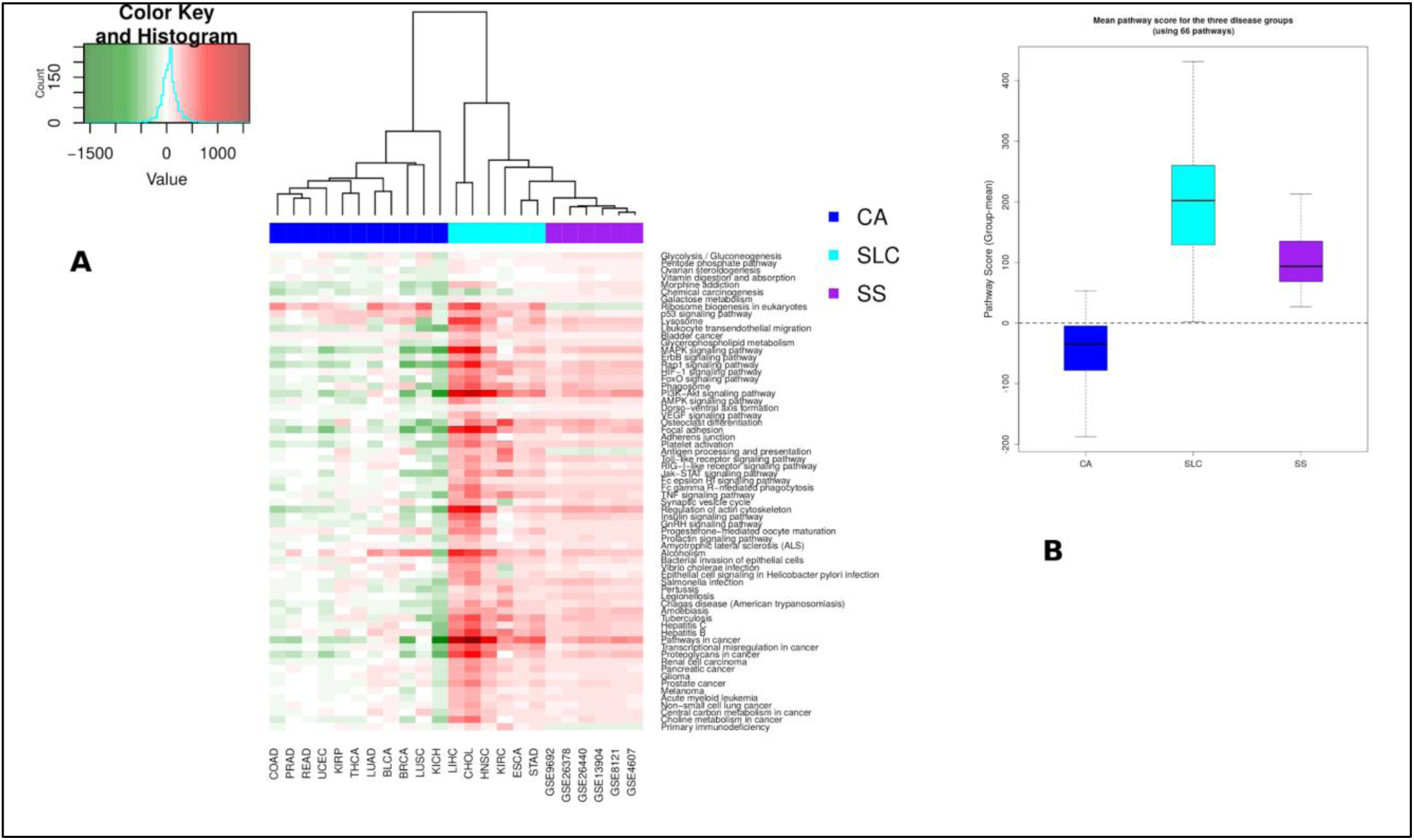
(A) Pathway heat map for 66 selected KEGG pathways where each pathway is significantly perturbed in at least 80% of the studies in one or both of the two groups of cancers. Colour of the cell is explained by the colour key histogram provided on the top left. Green indicated down-regulation of pathway, red indicates up-regulation. In the column-sidebar, purple indicates SS group, cyan indicates SLC group and blue indicates CA group. Positive pathway score is presented in red, negative pathway score in green. (B) Box plot shows group-mean pathway scores for the three groups of diseases. In general, there is pathway up-regulation in SLC group, even higher than that in SS, while down-regulation in CA group.

### Network analysis

Genome-level biological significance of the selected pathways was revealed by network analysis. Here, the network consisted of all KEGG pathways as nodes and substantial pairwise gene-overlap among pathways as edges. The centrality of the selected pathways was further quantified in terms of degree and betweenness centrality. Degree captures short-range connectivity of a node (i.e., how many nodes are connected to this node?), while betweenness centrality captures long-range centrality (i.e., how often this node falls on the shortest path between any two other nodes?) Degree captures the number of immediate neighbours of a node, while betweenness captures the essentiality of a node to the structure of the network. The degree and betweenness values for the selected 66 pathways were recorded (Additional File 5: Table S2).

The degree distribution of the network showed scale-free property of a biological network, i.e., many nodes with few edges and a few nodes with large number of edges (Additional File 3: Figure S3 - panel A). Visual exploration revealed the selected pathways at the core of the network (Fig. 3), and this was confirmed by comparing the degree of these 66 pathways with other pathways in the human genome. As shown in Additional File 3: Figure S3 (panel B), selected pathways had higher degree than the other pathways.

**Fig. 3:**
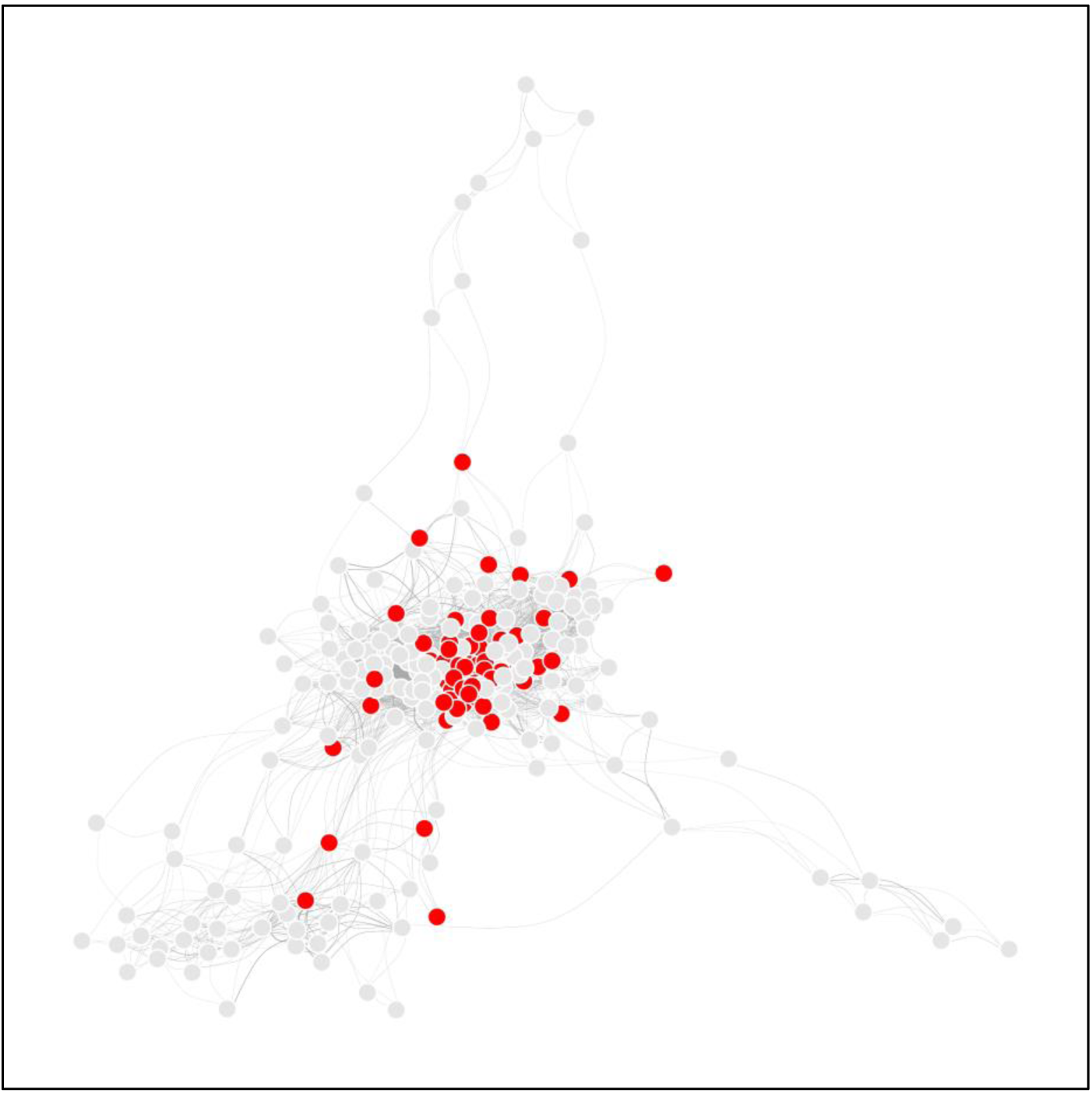
KEGG overlap network is constructed with each KEGG pathway as a node and overlapping gene membership between two pathways as an edge. The selected 66 pathways are shown in red, while the other pathways are shown in grey. Red nodes (selected pathways) have more connecting edges than the grey nodes, thus representing hub-like nature of selected pathways. This suggests that key biological processes are perturbed in both SS and SLC.

### Machine learning-based prediction of cancer group (SLC/CA) from 66 pathway scores

In order to assess the clinical relevance of the selected pathways, we asked if it is possible to predict (by a machine learning algorithm) the cancer group based on the selected pathway scores of an individual patient of cancer. For each patient, 66 pathway scores were input for the classifier, with the expected output being SLC or CA. Five-fold cross-validation was employed with balanced partitioning in each fold. Both algorithms - Support Vector Machine (SVM) and Neural Net (NN) – performed with high accuracy in assigning the samples to their respective cancer groups. SVM correctly assigned 537 (99.08%) out of 542 patients, while NN correctly assigned 536 (98.89%) out of 542 patients, with a misclassification rate of 0.9% and 1.1% respectively (Table 2). This independent validation showed the clinical relevance of the selected 66 pathways for prediction of cancer group from individual patient-level data.

**Table 2:** Two machine learning algorithms - Support Vector Machine (SVM) and Neural Network (NN) - were implemented through the function call MLearn (of R package MLInterfaces). SVM was applied with default parameters. Neural Network applied with three nodes in the hidden layer, with weight decay set to 0.01. Five-fold cross-validation was performed by generating a partition function (for cross-validation, where the partitions are approximately balanced with respect to the distribution of cancer tissue types (SLC or CA) in training and test sets in each fold. Confusion matrix from the classifier output was collected for calculation of misclassification rates.

| **Support Vector Machine** |  | **Predicted** |  |
| --- | --- | --- | --- |
|  |  | **CA** | **SLC** |
| **Given** | **CA** | **349** | **1** |
|  | **SLC** | **4** | **188** |
| **Misclassification rate** |  |  | **0.9** |
| **Neural Network** |  | **Predicted** |  |
|  |  | **CA** | **SLC** |
| **Given** | **CA** | **347** | **3** |
|  | **SLC** | **3** | **189** |
| **Misclassification rate** |  |  | **1.1** |

### Survival analysis

Another measure of clinical relevance of pathway signature was provided by its association with survival. For a given pathway, the patients were divided into two groups of subjects (high-scoring and low-scoring) based on the level of pathway score. Survival probability for these two groups was assessed both graphically and statistically. For many of the pathways, there was a higher pathway score in survivors compared to non-survivors in SLC. This directionality was also maintained in SS, i.e., higher pathway score in survivors compared to non-survivors (Additional File 6: Table S3). The number of pathways associated (p < 0.1) with survival varied: 9 (14%) in HNSC, 28 (42%) in LIHC and 34 (52%) in KIRC (Fig. 4A). Representative plots were drawn to show survival difference between high-score and low-score patients for the pathway “Fc gamma R-mediated phagocytosis” (Fig. 4B-D).

**Fig. 4:**
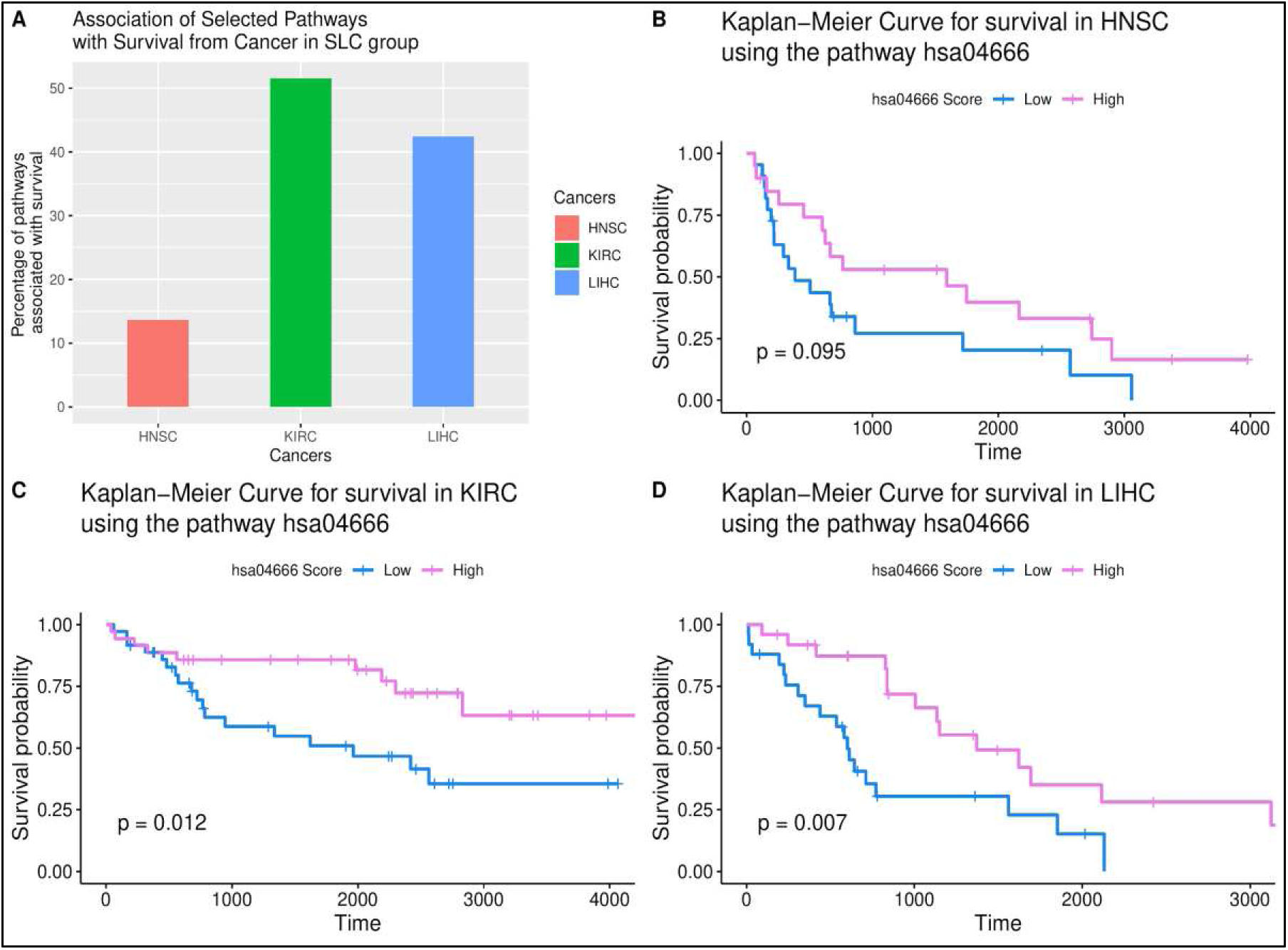
(A) Three SLC studies were selected for survival analysis. For each cancer, association with survival was accepted at threshold of p < 0.1. For each cancer type, proportion of pathways associated with survival was calculated and converted to a percentage. The percentage values are presented in this bar plot. As shown, up to 50% of the pathways are associated with survival in the SLC group. (B-D) Kaplan-Meier plot showing significant

### Functional classification of the pathways

The selected 66 pathways were divided into categories based on their functional relevance. The categories included: immune system, infection, cancer, catabolism, signal transduction, ribosome biogenesis and carbohydrate metabolism (Additional File 8: Text S1).

### Viral integration in TCGA samples

Out of the 687 cancer samples from TCGA analysed by us, there was overlap of 425 (61.9%) samples with the samples analysed for viral integration by Tang *et al*. [25]; of the 425 samples, 120 belonged to SLC group, and 305 belonged to CA group. Viral integration was observed in 9 (7.5%) SLC samples and 2 (0.66%) CA samples, showing significantly (p = 0.0003) higher viral integration in SLC. In another study by Cao *et al.* [26], there was an overlap of 533 (77.6%) of 687 samples analysed by us. Of the 533, 200 (37.5%) were SLC while 333 (62.5%) were CA cancers. There was significantly (p = 0.001) higher viral integration in SLC (30%) compared to CA (17.4%). The report by Kazemian *et al.* [27] was not associated with supplementary sample-level data. However, re-analysis of summary data provided (Additional File 8: Table S5) led to the finding of significantly (p = 4.10E-13) higher viral integration in SLC (18.7%) compared to CA (5.6%). Over all, the findings were consistent with SLC cancers to have greater likelihood of prediction of viral integration.

## Discussion

Comprehensive system-level analysis reveals shared transcriptomic response between septic shock (SS) and a subset of cancers (SLC). The SLC group predominantly consists of cancer of the upper GI tract, including head and neck, oesophagus, stomach, liver and biliary tract. The striking segregation of the SLC group with SS suggests shared elements of pathophysiology operational in these disorders. Sixty-six pathways differentially expressed in both SS and SLC represent critical biological processes, such as metabolism, immune response and protection against infection. Network analysis reveals the selected pathways as a core functional module (Figure 3) of the genome-scale network dysregulated in both SS and SLC. Many of these processes, such as metabolism and inflammatory response are known to contribute to the pathogenesis of both sepsis and cancer [9].

The segregation of cancer into two groups (SLC and CA) prompted us to investigate the prospect of the selected pathways as a classifier. Machine learning-based validation of an independent data set proved that the pathways indeed consist of a robust sample-level signature of the cancer groups. It is thus possible to predict, with high accuracy, whether a patient belongs to a particular cancer group (SLC or not). With plummeting costs and increasing acceptability of clinical transcriptomics, this signature may be useful in future for patient stratification.

Similarity in the direction of change in pathway activity immediately evokes an explanation in terms of shared immunological response in either case. SLC are often associated with infection (i.e., human papilloma virus in “head and neck”, *H. pylori* in stomach, hepatitis viruses in liver). In fact, TCGA data analysed for the presence of viral sequence reads in RNA-seq datasets of several cancers have revealed infection status of the samples. Interestingly, the majority of the infected samples belong to the GI associated cancers including head and neck (HNSC), oesophagus (ESCA), stomach (STAD) and liver (LIHC) [25–29]. Cao et al. (2016) [26] estimated high percentage of infected samples in TCGA datasets for liver (21.2%), head and neck (16.6%), stomach (14.8%) and oesophagus (12%). It is known that sepsis starts out as an infection, but it is the uncontrolled host response that drives the phase of shock. Similarly, in cancers with an infectious origin, host response (e.g., inflammation) plays a role in malignant transformation [9]. Pathways related to immune response are consistently differentially expressed in both SS and SLC. Although the time scale of pathogenesis is much shorter in SS compared to cancer, our finding calls for greater attention into the shared cellular processes underlying these two distinct clinical phenotypes.

The finding of pathway up-regulation associated with survival brings new insight into the relationship between the shared transcriptomic patterns and disease outcome. Sepsis is a highly lethal dysregulated host response to infection, and SLC, by extension, appears to share a part of that response. Since the pathways are selected on the basis of up-regulation in both SLC and SS (compared to control), plausible explanation is that the pathway up-regulation is protective rather than detrimental to the human host. It is worth mentioning that viral integration or bacterial infection in a setting of cancer has been reported to favour survival in SLC cancers of oropharyngeal [30], liver [31] and kidney [32].

Alteration of intestinal permeability is known in both sepsis and cancer [33]. It may be speculated that anatomical proximity increases the likelihood of the liver and upper gastrointestinal tissue to be exposed to the translocated microbial products through a permeable gut, eliciting a host response in both SS and SLC. Significant association between survival and up-regulation of phagocytosis-associated pathway (i.e., Fc gamma R-mediated phagocytosis) lends support to this view. In general, sepsis is less lethal in patients with cancer of the digestive system than cancer of other organs [8]. The role of gut in survival from sepsis and cancer is an active area of research with translational potential.

## Conclusions

Firstly, transcriptome-based systems biology approach segregates cancer into two groups (SLC and CA) based on similarity with SS. Secondly, the similarity is based on a set of pathways associated with pathogenesis of sepsis and cancer. These pathways form a robust signature of a novel cancer grouping. Lastly, up-regulation of the pathways is protective rather than detrimental to the patient. A mechanism of the shared protective host response is hypothesized in terms of immunocompetence induced by microbial products from the permeable gut. This work is the first step towards a systems biology-based patient stratification. It is hoped that future work in this direction shall generate actionable knowledge for clinical management of both cancer and septic shock.

## List of abbreviations

BLCA: Bladder Urothelial Carcinoma
BRCA: Breast invasive carcinoma
CA: Cancer Alone
CHOL: Cholangiocarcinoma
COAD: Colon adenocarcinoma
ESCA: Esophageal carcinoma
FPKM: Fragments Per Kilobase of transcript per Million mapped reads
GEO: Gene Expression Omnibus
HNSC: Head and Neck squamous cell carcinoma
ICU: Intensive Care Unit
KEGG: Kyoto Encyclopaedia of Genes and Genomes
KICH: Kidney Chromophobe
KIRC: Kidney renal clear cell carcinoma
KIRP: Kidney renal papillary cell carcinoma
LFC: log2(fold change)
LIHC: Liver hepatocellular carcinoma
LUAD: Lung adenocarcinoma
LUSC: Lung squamous cell carcinoma
NCBI: National Center for Biotechnology Information
NN: Neural Net
PRAD: Prostate adenocarcinoma
READ: Rectum adenocarcinoma
SLC: Sepsis-Like Cancers
SS: Septic shock
STAD: Stomach adenocarcinoma
SVM: Support Vector Machine
TCGA: The Cancer Genome Atlas
THCA: Thyroid carcinoma
UCEC: Uterine Corpus Endometrial Carcinoma

## Declarations

### Ethics approval and consent to participate

In this study, patient data from public databases (GEO and TCGA) have been analysed. We have not used any patient data that are not already available in the public domain.

### Consent to publish

Not applicable.

### Availability of data and materials

All programming was done in the R programming language [20]. Data and R code are available at the following link: https://figshare.com/ (https://doi.org/10.6084/m9.figshare.8118413.v3, https://doi.org/10.6084/m9.figshare.8118647.v5). The whole data can be downloaded as a single zip file (tcnibmgV3doc.zip). Upon uncompressing the zip, instructions for running the code and generating the figures of this manuscript are available in the file howto.pdf.

### Authors’ contributions

SKM conceptualized the study and contributed to Funding Acquisition, Project Administration, Resources, Supervision. HT contributed to Data Curation. SM contributed to Data Curation, Software, Validation, Visualization. All authors contributed to Formal Analysis, Investigation, Methodology, Manuscript writing and approved of the final manuscript.

## Acknowledgements

We thank the National Cancer Institute for making TCGA data available in the public domain. We thank the National Center for Biotechnology Information for making GEO data available in the public domain. This study would not have been possible without data from these two resources. We thank the Director of National Institute of Biomedical Genomics for making the computational resources accessible to us. HT and SKM thank the Department of Biotechnology, Government of India for financial support to carry out the analysis. SKM acknowledges very helpful advice received from Prof. Subrata Sinha during the revision of the manuscript.

## Authors’ information

HT and SM are co-first authors. All authors are affiliated to the National Institute of Biomedical Genomics, P.O. NSS, Kalyani, Nadia, West Bengal, India. 741251.

## Supplementary Information

**Supplementary Figure S1.**
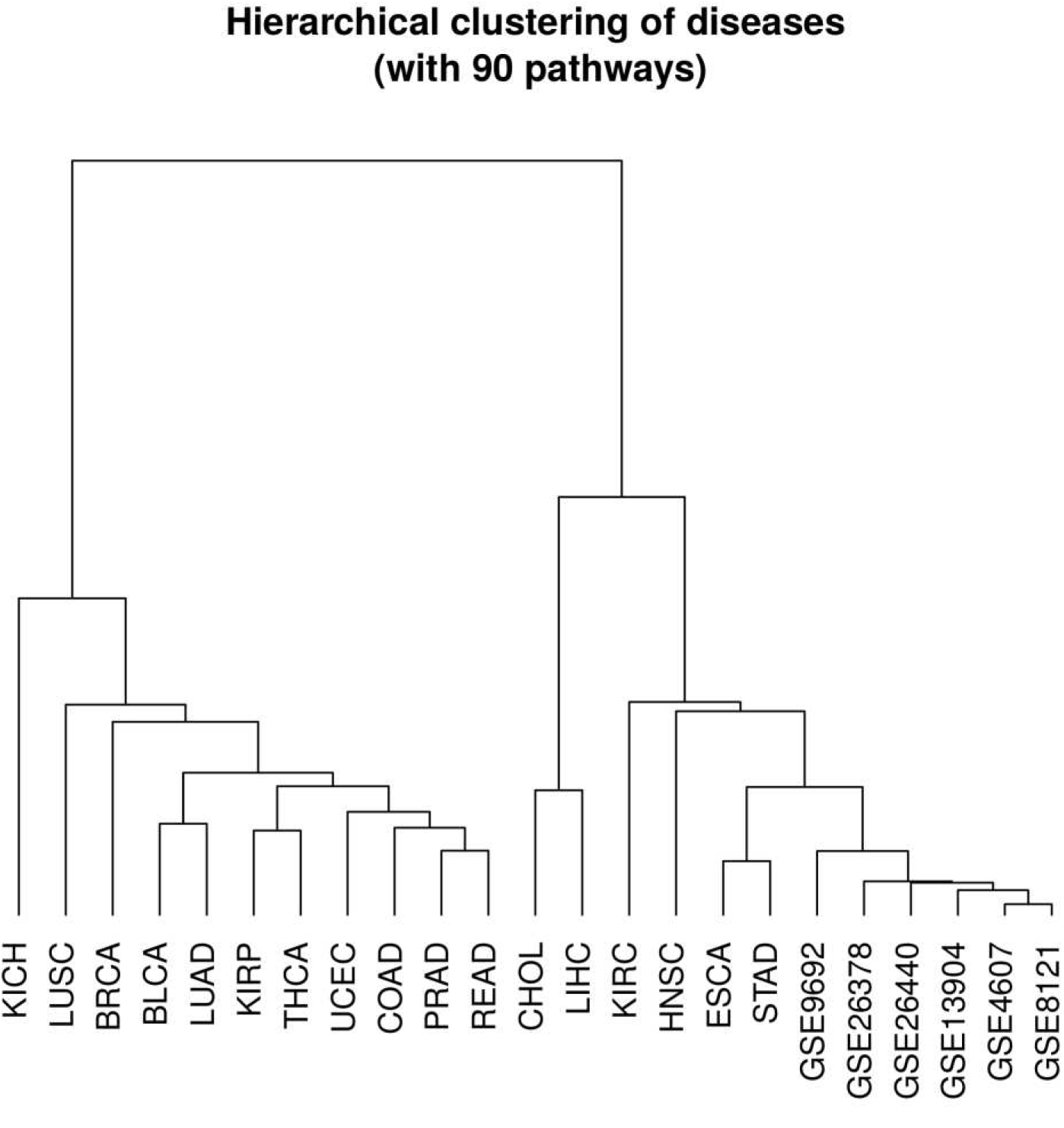
For the 90 pathways significantly associated with SS, pathway scores were computed for the cancer studies. Scores for SS (6 studies) and cancer (17 studies) were subjected to hierarchical clustering (details in main text). As shown in this dendrogram, there is clear segregation of the cancer studies into two groups: sepsis–like (SLC) and other cancer (CA).

**Supplementary Figure S2.** Gene level heat map of each of the selected 66 pathways. LFC for each gene was calculated from disease and normal samples (disease-normal) for all studies. For each disease type (CA, SLC or SS), median LFC was calculated across the studies in that disease type. Combined heat map was generated based on the pathway genes and the three disease groups. We considered only those genes which have similar LFC directionality in SLC and SS. Red colour represents up-regulation in disease compared to control, while green colour represents down-regulation. Pathway names with KEGG identifiers are shown on the top, gene symbol and names are row-labels of heat map. The major class (to which the pathway belongs) is shown at the bottom.

**Supplementary Figure S3.**
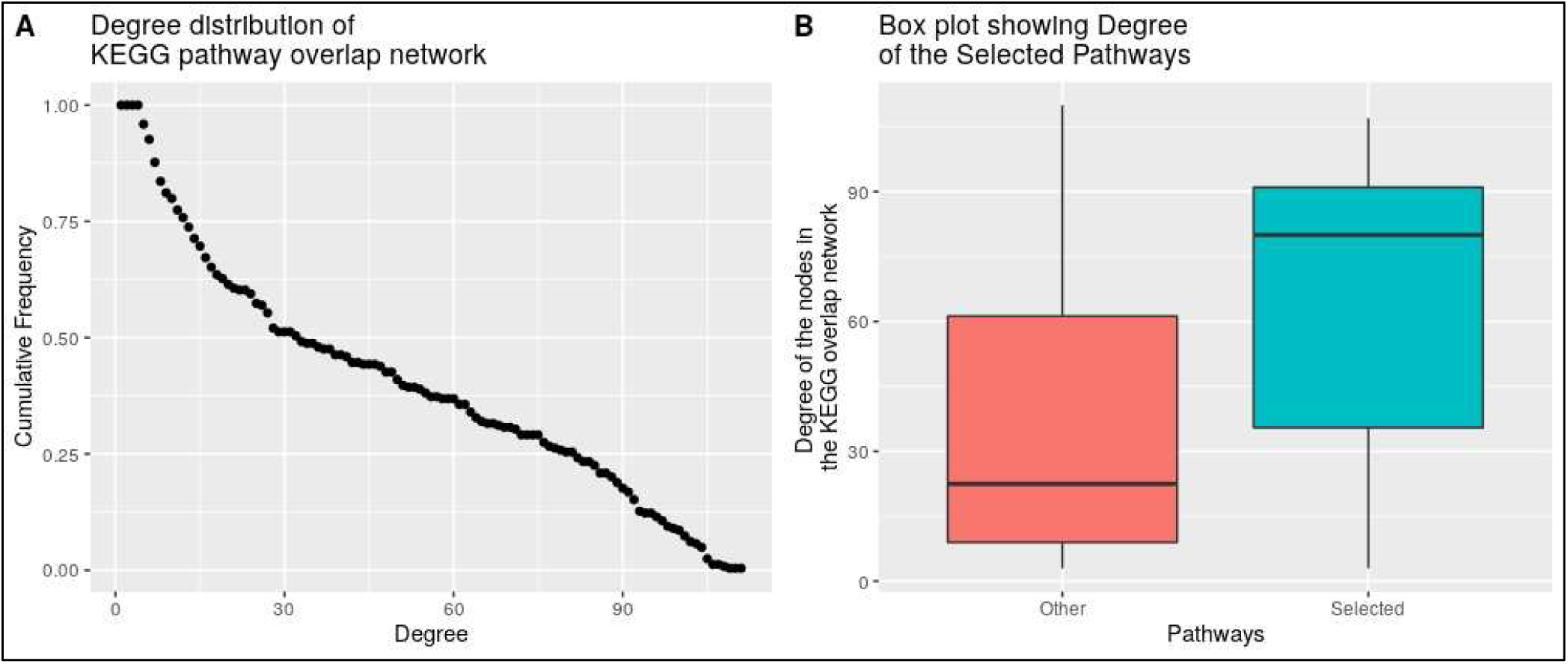
(A) This is the degree distribution of the network generated by overlapping KEGG pathways. This shows that there are many nodes with very low connectivity but a few nodes with very high connectivity. This is consistent with the scale-free property of biological networks. (B) Box plot showing comparison of degree between two groups of nodes in this network

**Supplementary Table S1.**
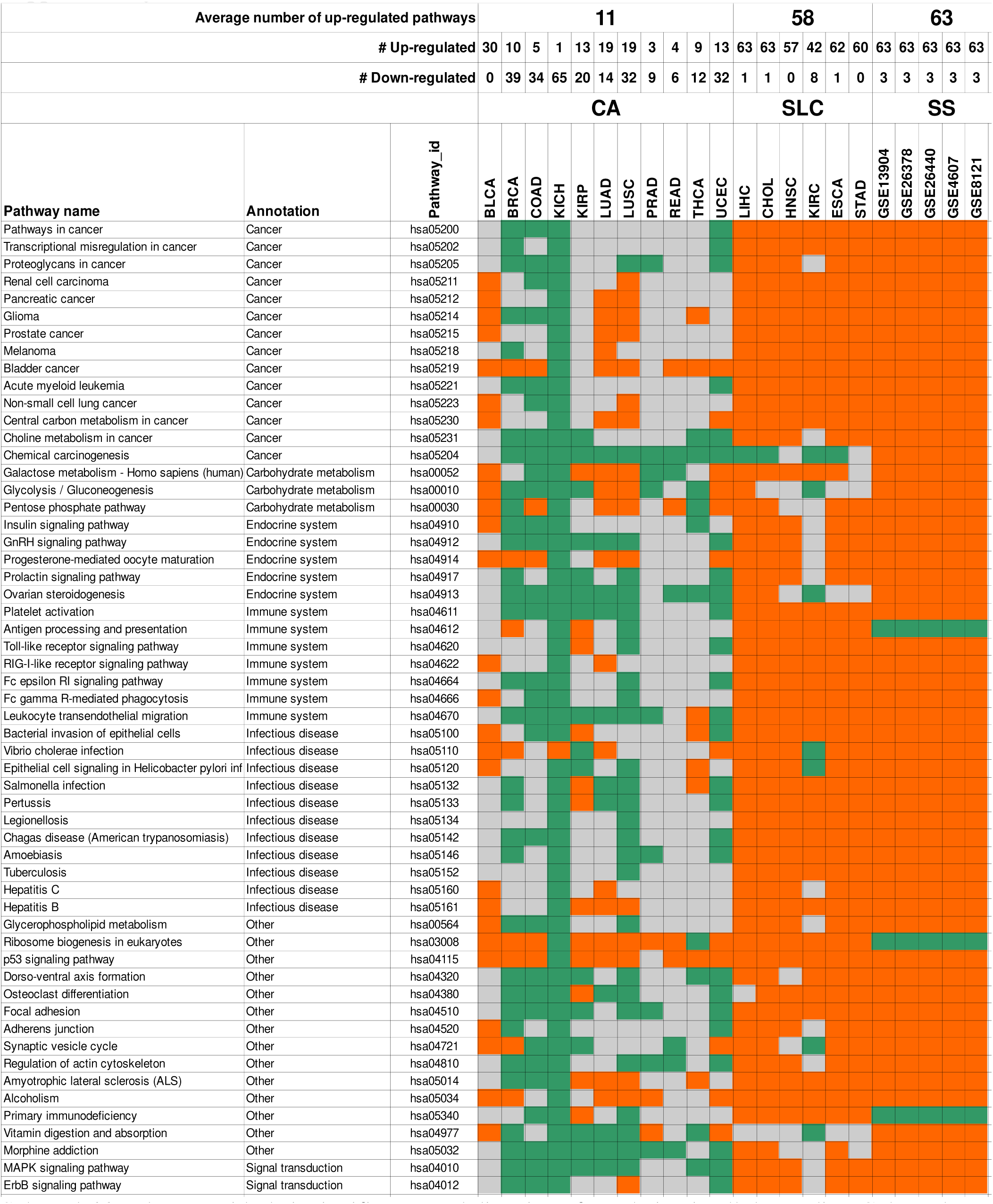
Selected 66 pathways with their significance and direction of regulation in all the studies. Coloured cells are a representation of pathway scores. Red indicates significant up-regulation, green indicates significant down-regulation and grey indicates non-significant change in pathway. Number of up- and down-regulated pathways in each study is shown at the top. Group membership (to CA, SLC or SS) is shown as a label on top of the code of the data set. First three columns of the sheet are pathway name, annotation (functional class) and KEGG ID respectively.

**Supplementary Table S2:** This table contains the property of each node of the network. Degree of a node is the number of edges connected to a node. Betweenness centrality is a measure of how much information flows through a specific node, i.e., average number of shortest paths passing through the node. Degree refers to the number of neighbours of a node while betweenness centrality refers to how important the node is for maintaining the overall structure of that network. The two nodes with highest betweenness centrality (Glycolysis/Gluconeogenesis, Lysosome) in the network are highlighted.

| KEGG Pathway Names | KEGG ID | Degree | Betweenness Centrality |
| --- | --- | --- | --- |
| Vitamin digestion and absorption | hsa04977 | NA | NA |
| Ribosome biogenesis in eukaryotes | hsa03008 | NA | NA |
| Glycolysis / Gluconeogenesis | hsa00010 | 26 | 1599.07 |
| Lysosome | hsa04142 | 7 | 680.2 |
| Insulin signaling pathway | hsa04910 | 89 | 560.44 |
| Epithelial cell signaling in Helicobacter pylori infection | hsa05120 | 49 | 520.57 |
| AMPK signaling pathway | hsa04152 | 62 | 520.28 |
| Vibrio cholerae infection | hsa05110 | 37 | 439.59 |
| Choline metabolism in cancer | hsa05231 | 90 | 433.04 |
| Tuberculosis | hsa05152 | 87 | 410.94 |
| Central carbon metabolism in cancer | hsa05230 | 81 | 403.15 |
| Ovarian steroidogenesis | hsa04913 | 48 | 387.79 |
| GnRH signaling pathway | hsa04912 | 107 | 383.52 |
| Hepatitis B | hsa05161 | 106 | 347.57 |
| Chagas disease (American trypanosomiasis) | hsa05142 | 104 | 328.8 |
| HIF-1 signaling pathway | hsa04066 | 80 | 323.29 |
| Fc epsilon RI signaling pathway | hsa04664 | 102 | 317.94 |
| Pertussis | hsa05133 | 67 | 295.91 |
| Progesterone-mediated oocyte maturation | hsa04914 | 103 | 287.93 |
| Pathways in cancer | hsa05200 | 98 | 286.47 |
| VEGF signaling pathway | hsa04370 | 103 | 265.42 |
| Amoebiasis | hsa05146 | 99 | 248.79 |
| Proteoglycans in cancer | hsa05205 | 100 | 230.53 |
| FoxO signaling pathway | hsa04068 | 90 | 206.86 |
| Phagosome | hsa04145 | 31 | 195.33 |
| PI3K-Akt signaling pathway | hsa04151 | 80 | 176.5 |
| Fc gamma R-mediated phagocytosis | hsa04666 | 83 | 170.39 |
| MAPK signaling pathway | hsa04010 | 90 | 167.12 |
| Platelet activation | hsa04611 | 100 | 166.21 |
| Toll-like receptor signaling pathway | hsa04620 | 91 | 156.38 |
| Rap1 signaling pathway | hsa04015 | 96 | 153.11 |
| Morphine addiction | hsa05032 | 46 | 152.64 |
| Prostate cancer | hsa05215 | 95 | 150.98 |
| Glioma | hsa05214 | 97 | 145.97 |
| TNF signaling pathway | hsa04668 | 91 | 136.56 |
| Hepatitis C | hsa05160 | 91 | 131.46 |
| Antigen processing and presentation | hsa04612 | 26 | 123.22 |
| Bacterial invasion of epithelial cells | hsa05100 | 56 | 116.33 |
| Pancreatic cancer | hsa05212 | 91 | 101.47 |
| Osteoclast differentiation | hsa04380 | 87 | 97.12 |
| Focal adhesion | hsa04510 | 80 | 90.01 |
| Acute myeloid leukemia | hsa05221 | 84 | 78.81 |
| Chemical carcinogenesis | hsa05204 | 14 | 75.3 |
| Leukocyte transendothelial migration | hsa04670 | 66 | 72.24 |
| Non-small cell lung cancer | hsa05223 | 86 | 64.38 |
| Salmonella infection | hsa05132 | 48 | 61.81 |
| Alcoholism | hsa05034 | 46 | 60.62 |
| Regulation of actin cytoskeleton | hsa04810 | 62 | 60.6 |
| Jak-STAT signaling pathway | hsa04630 | 61 | 58.74 |
| Prolactin signaling pathway | hsa04917 | 91 | 57.33 |
| Renal cell carcinoma | hsa05211 | 84 | 56.81 |
| ErbB signaling pathway | hsa04012 | 91 | 56.53 |
| Melanoma | hsa05218 | 81 | 45.43 |
| Synaptic vesicle cycle | hsa04721 | 10 | 42.04 |
| Galactose metabolism | hsa00052 | 8 | 41.96 |
| Amyotrophic lateral sclerosis (ALS) | hsa05014 | 30 | 39.8 |
| Glycerophospholipid metabolism | hsa00564 | 14 | 26.42 |
| Bladder cancer | hsa05219 | 53 | 18.91 |
| Adherens junction | hsa04520 | 27 | 17.54 |
| RIG-I-like receptor signaling pathway | hsa04622 | 39 | 17.21 |
| Legionellosis | hsa05134 | 27 | 9.08 |
| Dorso-ventral axis formation | hsa04320 | 26 | 2 |
| p53 signaling pathway | hsa04115 | 14 | 1.21 |
| Transcriptional misregulation in cancer | hsa05202 | 6 | 0.78 |
| Primary immunodeficiency | hsa05340 | 5 | 0.29 |
| Pentose phosphate pathway | hsa00030 | 3 | 0 |

**Supplementary Table S3:**
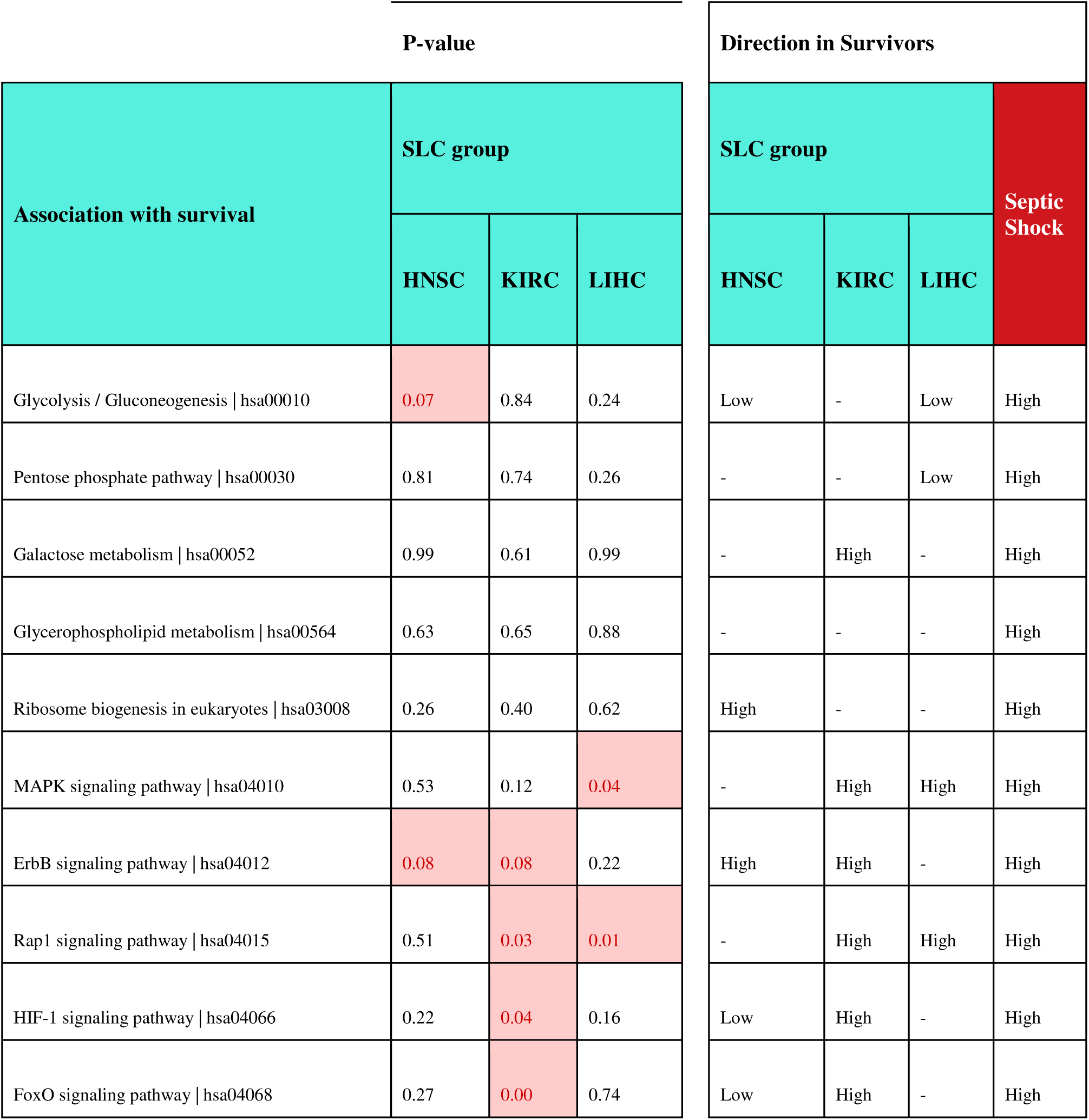
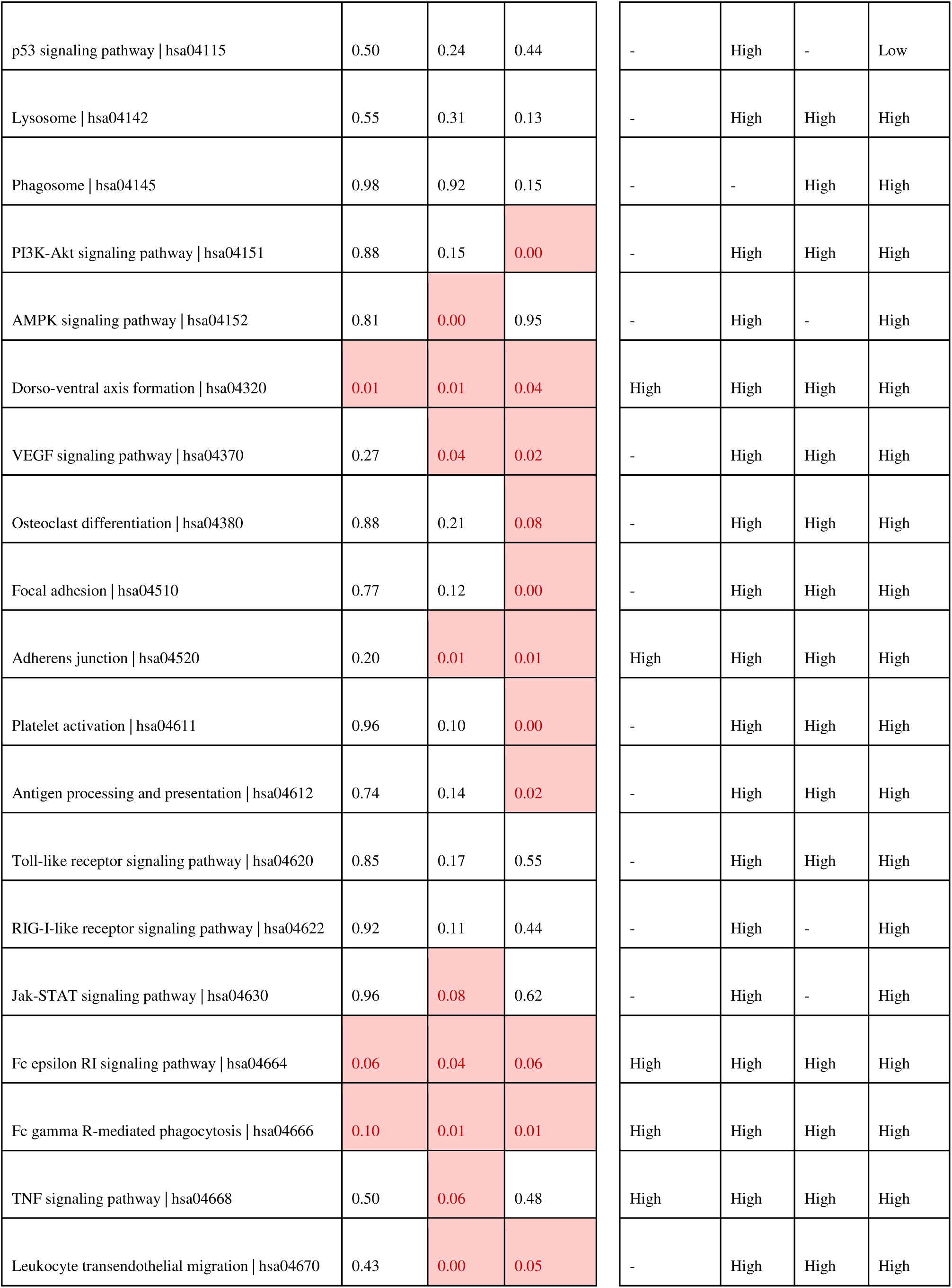
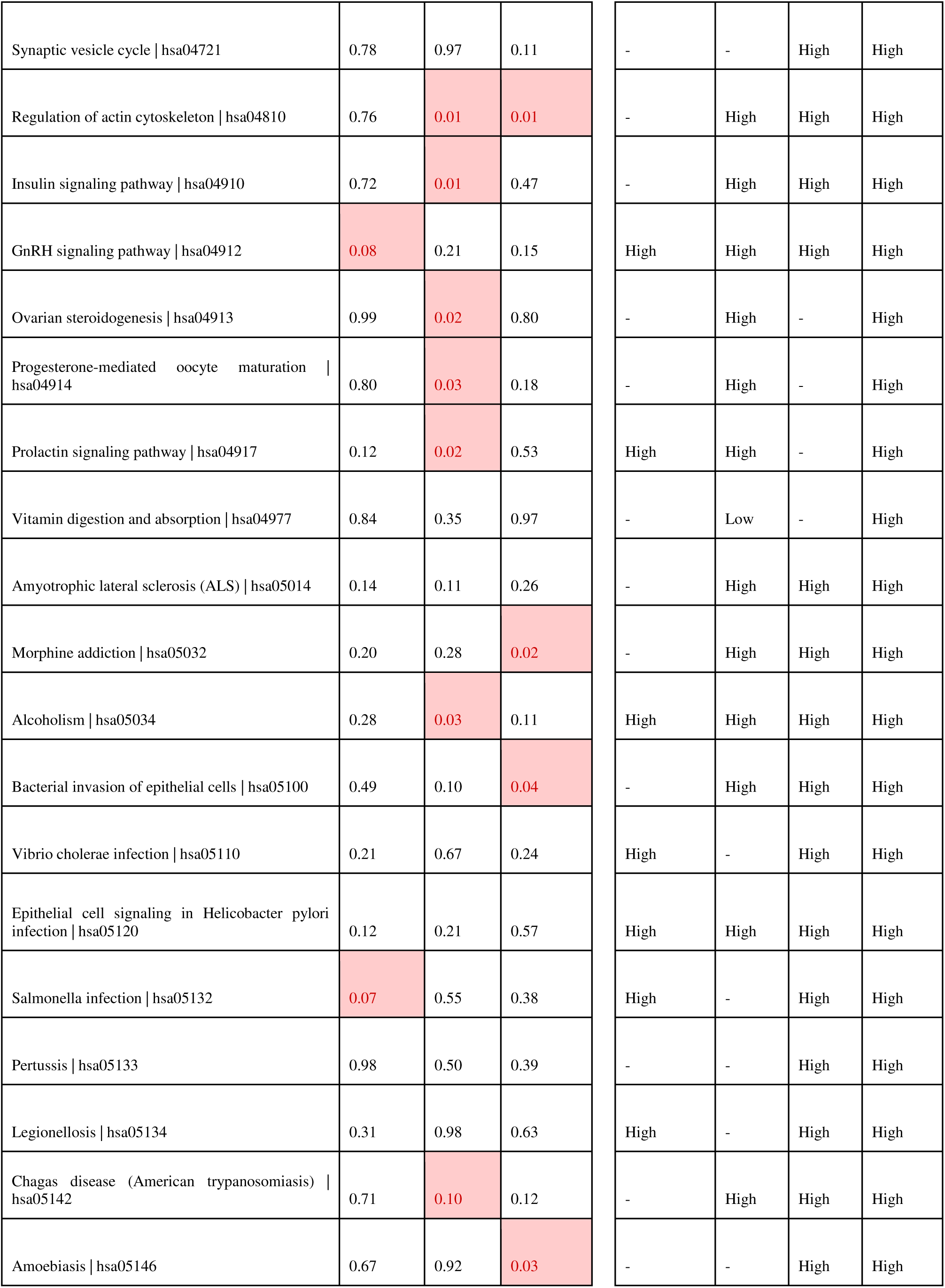
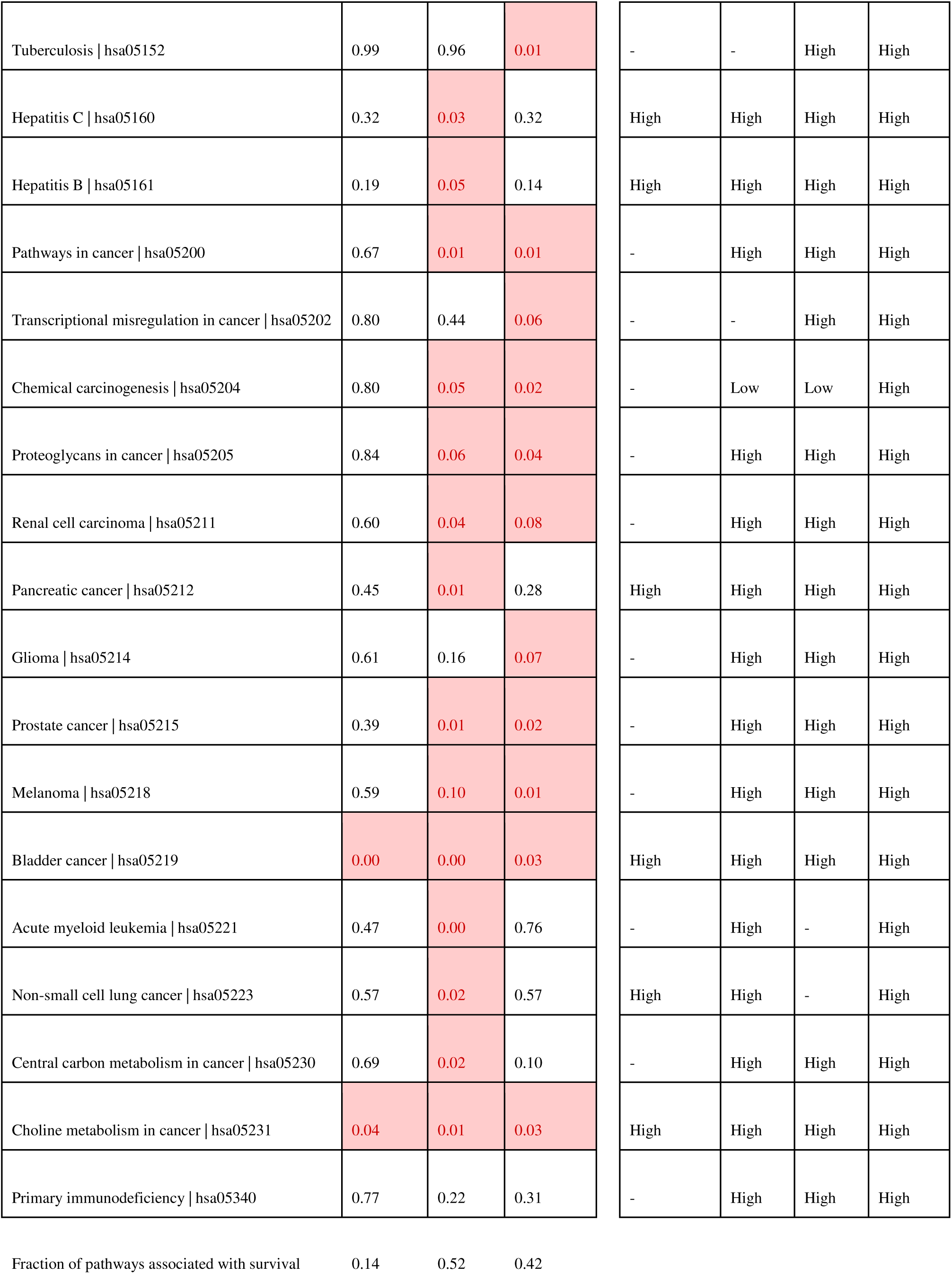
Three SLC cancers were selected for survival analysis. The value in each cell corresponds to the p-value for association of the pathway with the cancer type. At a threshold of p < 0.1, the cell has been high-lighted, noting significant association of the pathway with survival from cancer. Direction of pathway expression score was estimated from visual inspection of survival plot in the SLC cancers. For each cancer, proportion of pathways associated with survival was calculated and converted to percentage (bottom row of the table). For septic shock, pathway score was calculated in the survivor group compared to the non-survivor group, as described in the Methods. A direction of “High” was assigned, if the pathway was observed to be higher in the survivor group for four or more of the eight septic shock data sets tested. Otherwise, the direction of “Low” was assigned.

**Supplementary Table S4:** List of additional cancer data sets for machine learning-based prediction of cancer samples.

| GEO_ID | Cancer | Type of tumour | Number of samples | Number of tumour samples used in analysis | Number of control samples used in analysis | Platform | Normalization |
| --- | --- | --- | --- | --- | --- | --- | --- |
| **GSE103512** | Breast, Colorectal, Prostate, And Non-Small Cell Lung Cancer | Solid Tumor | 280 | 242 | 38 | GPL13158[HT_HG-U133_Plus_PM] Affymetrix HT HG-U133+ PM Array Plate | Quantile |
| **GSE112790** | Hepatocellular Carcinoma | Solid Tumor | 198 | 183 | 15 | GPL570 [HG-U133_Plus_2] Affymetrix Human Genome U133 Plus 2.0 Array | Quantile |
| **GSE32665** | Lung adenocarcinoma | Solid Tumor | 170 | 85 | 85 | GPL6102Illumina human-6 v2.0 expression beadchip | Quantile |
| **GSE33479** | Lung squamous carcinogenesis | Solid Tumor | 122 | 14 | 27 | GPL6480 Agilent-014850 Whole Human Genome Microarray 4x44K G4112F (Probe Name version) | Natural log of (Cy5/Cy3) |
| **GSE84984** | Colorectal Cancer | Solid Tumor | 15 | 9 | 6 | GPL17586 [HTA-2_0] Affymetrix Human Transcriptome Array 2.0 [transcript (gene) version] | RMA |
| **GSE103236** | Gastric Cancer | Solid Tumor | 19 | 9 | 9 | GPL4133Agilent-014850 Whole Human Genome Microarray 4x44K G4112F (Feature Number version) | Lowess |

**Supplemental Table S5:** Viral integration in TCGA samples. Results are derived from published data.

|  |  |  |  |  |  |  |
| --- | --- | --- | --- | --- | --- | --- |
|  | **Tang2013 [a]** |  | **Cao2016 [b]** |  | **Kazemina2015 [c] \*\*** |  |
|  | N (%) | P-value | N (%) | P-value | N (%) | P-value |
| Samples analysed in current manuscript | 687 | 3.00E-04 | 687 | 1.00E-03 | N.A. | 4.10E-13 |
| Overlap with published study | 425 (61.9%) |  | 533 (77.6%) |  | N.A. |  |
| SLC samples | 120 (28.2%) |  | 200 (37.5%) |  | 305 |  |
| CA samples | 305 (71.8%) |  | 333 (62.5%) |  | 1175 |  |
| Viral integration in SLC | 9 (7.5%) |  | 60 (30%) |  | 57 (18.7%) |  |
| Viral integration in CA | 2 (0.66 %) |  | 58 (17.4 %) |  | 66 (5.6%) |  |
|  |  |  |  |  | ** Sample-level data are not available. Number and proportion of viral integration in SLC and CA samples are calculated from summary data provided in the publication (Table 1 of [c]). |  |

- [a] Tang KW, Alaei-Mahabadi B, Samuelsson T, Lindh M, Larsson E. The landscape of viral expression and host gene fusion and adaptation in human cancer. *Nature Communications*. 2013;4:1–9. - [b] Cao S, Wendl MC, Wyczalkowski MA, Wylie K, Ye K, Jayasinghe R, et al. Divergent viral presentation among human tumors and adjacent normal tissues. *Scientific Reports*. 2016;6 June:1–12. - [c] Kazemian M, Ren M, Lin J-X, Liao W, Spolski R, Leonard WJ. Possible Human Papillomavirus 38 Contamination of Endometrial Cancer RNA Sequencing Samples in The Cancer Genome Atlas Database. *Journal of Virology*. 2015;89:8967–73.

### Additional text : Functional classification of the pathways

The selected 66 pathways are divided into categories based on their functional relevance. The categories include catabolism, immune system, infection, cancer, signal transduction, ribosome biogenesis and carbohydrate metabolism. Many of these pathways are related to immune response to infection, such as, clot formation, proinflammatory effects, antiviral effects, chemotactic effects and T cell stimulation, degranulation, cytokine gene transcription and generation of eicosanoids, phagocytosis, and migration of leukocytes to the site of injury, and are consistently up-regulated in both SS and SLC group.

#### Catabolism

Two catabolic pathways namely, Lysosome and Phagosome, are observed to be mostly up-regulated in SS and SLC (Fig. 2, Additional File 4: Table S1). Upon network analysis, it is revealed that lysosome pathway has a very high value for betweenness centrality (680.2) (Additional File 5: Table S2), thus being essential to the structure of the network. Likewise, Phagosome, another catabolic pathway linked to lysosome, is uniformly up regulated in SS and SLC groups (Additional File 2: Figure S2).

Muscle wasting is one of the most common phenomena in both SS and cancer. Lysosome mediated protein degradation has been noted on a substantial scale in such cases. Further, transcriptome based meta-analysis conducted by Ma et al., reported up-regulation of lysosome pathway in sepsis while increased biogenesis of lysosomes has been observed in cancer cells. In this study, we observe consistent up-regulation of catabolic pathways associated with lysosome and phagosome in SS and SLC, suggesting high catabolic rate in both the groups. Besides, lysosome is now considered a hub of metabolic signalling given its key role in maintaining cellular homeostasis. It also appears as a central node in our network with high betweenness centrality.

#### Immune system

Both cancer and septic shock are associated with dysregulation in immune function. It is observed that immune system pathways associated with clot formation (Platelet activation), proinflammatory effects, antiviral effects, chemotactic effects and T cell stimulation (Toll-like receptor signalling pathway), degranulation, cytokine gene transcription and generation of eicosanoids (Fc epsilon RI signalling pathway), phagocytosis (Fc gamma R-mediated phagocytosis), and migration of leukocytes to the site of injury (Leukocyte transendothelial migration) are uniformly up regulated in both SS and SLC groups (Additional File 2: Figure S2).

#### Infection

Infection is a common theme in both septic shock (which originates as sepsis) and many cancers (known to be induced by microbial infection). Many infection-associated pathways, such as, Bacterial invasion of epithelial cells, Vibrio cholerae infection, Epithelial cell signalling in Helicobacter pylori infection, Salmonella infection, Pertussis, Legionellosis, Tuberculosis, Chagas disease (American trypanosomiasis), Amoebiasis, Hepatitis C and Hepatitis B are significantly up regulated in SS and SLC group (Additional File 2: Figure S2).

#### Cancer pathways

Many cancer-associated pathways, such as Transcriptional misregulation in cancer, Proteoglycans in cancer, Central carbon metabolism in cancer, Choline metabolism in cancer, Chemical carcinogenesis, Renal cell carcinoma, Pancreatic cancer, Glioma, Prostate cancer, Melanoma, Bladder cancer, Acute myeloid leukemia, Non-small cell lung cancer are up regulated in SS and SLC group (Additional File 2: Figure S2).

#### Signal transduction

Signal transduction pathways involved in cell proliferation, differentiation, adhesion, migration, survival, cell cycle progression, anti-apoptosis (MAPK, ErbB, Rap1, VEGF, JAK-STAT and PI3k-Akt signalling pathways) and cell cycle regulation, apoptosis, autophagy, oxidative stress response, DNA repair, immune regulation, muscle atrophy, growth arrest, leukocyte recruitment and activation, synthesis of inflammatory mediators, remodelling of extracellular matrix (HIF1, AMPK, FoxO and TNF signalling pathways) are uniformly up-regulated in SS group, and in most of the SLC group (Additional File 2: Figure S2).

#### Endocrine system

Five pathways associated with endocrine system viz., Insulin signalling, GnRH signalling, Progesterone-mediated oocyte maturation, Prolactin signalling pathway and Ovarian steroidogenesis are up-regulated in all SS groups as well as in SLC group (Additional File 2: Figure S2).

#### Carbohydrate metabolism

Carbohydrate metabolism pathways including Galactose metabolism, Glycolysis/Gluconeogenesis, and Pentose phosphate pathway are up-regulated in SS group and most of the SLC group cancers. In our study, Galactose metabolism pathway is up-regulated in SLC and SS groups. Glycolysis/Gluconeogenesis (a source of energy and metabolite precursors of other pathways) plays a central role in cell as reflected by highest betweenness centrality parameter value i.e. 1599.06 (Additional File 5: Table S2). Pentose phosphate pathway, which generates precursors for synthesis of purine, pyrimidine and histidine, is up-regulated across the SS group along with the LIHC, CHOL, ESCA and STAD of SLC group.

### Gene–level heatmap for 66 pathways

#### Cancer

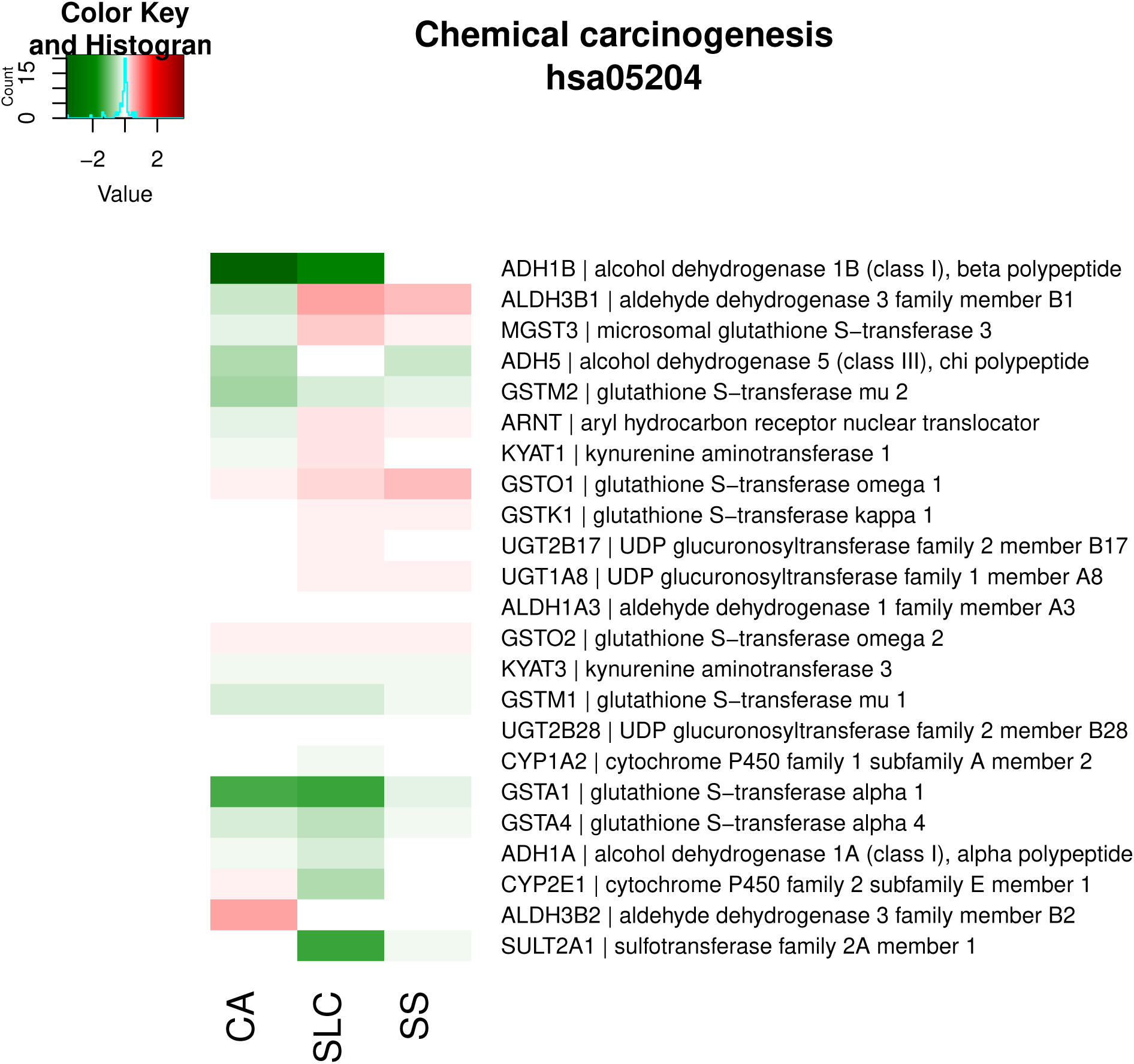

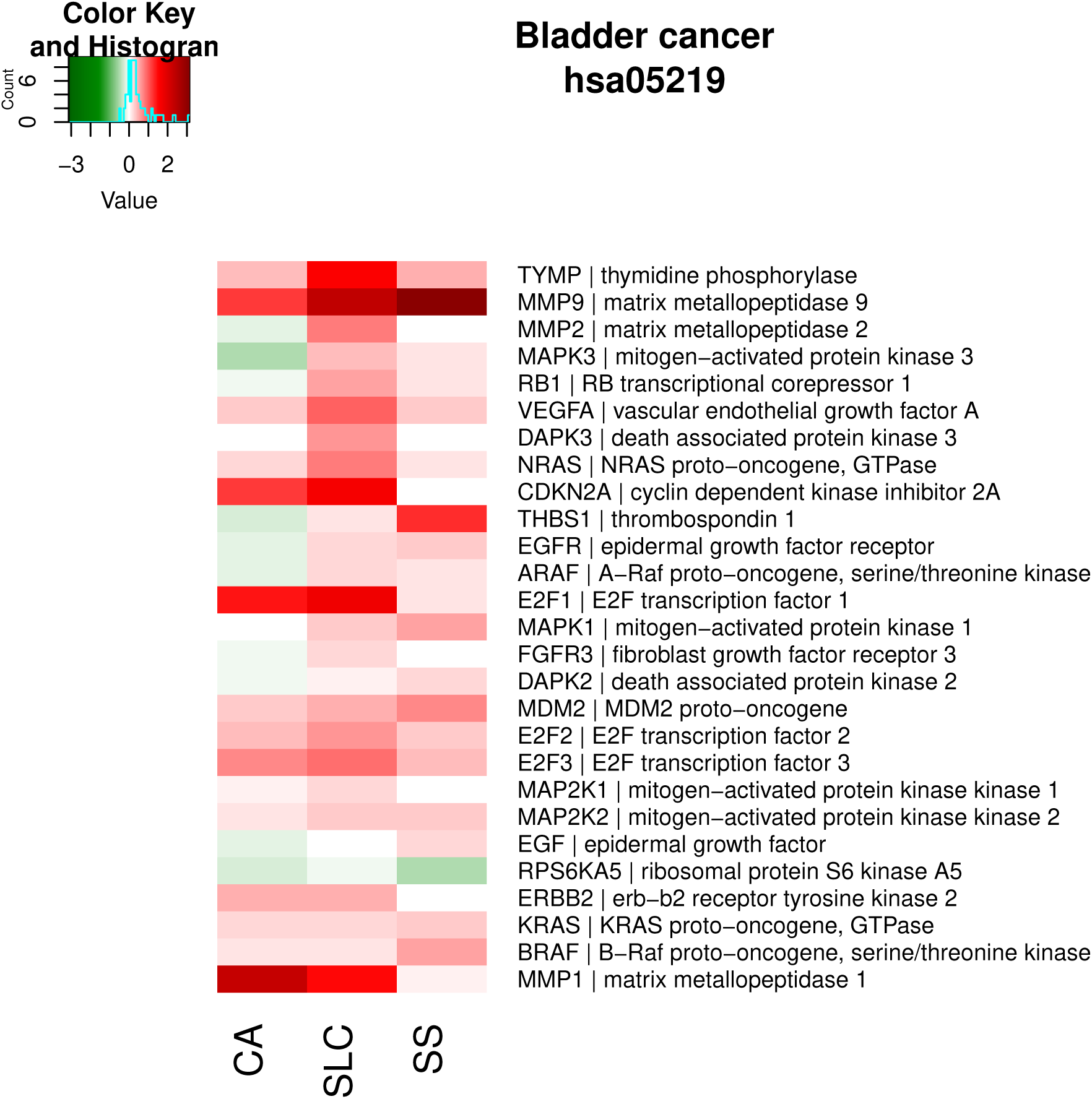

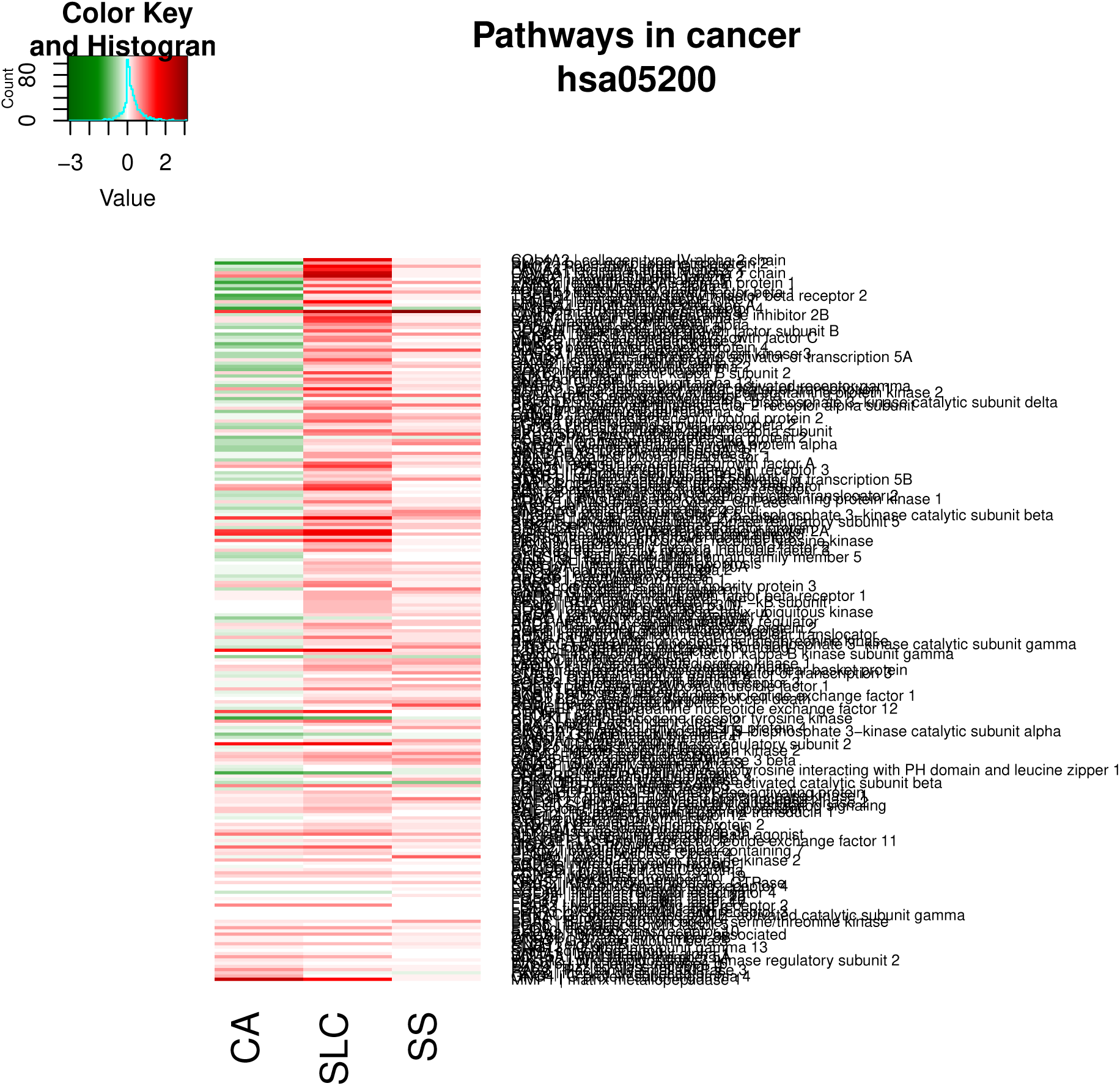

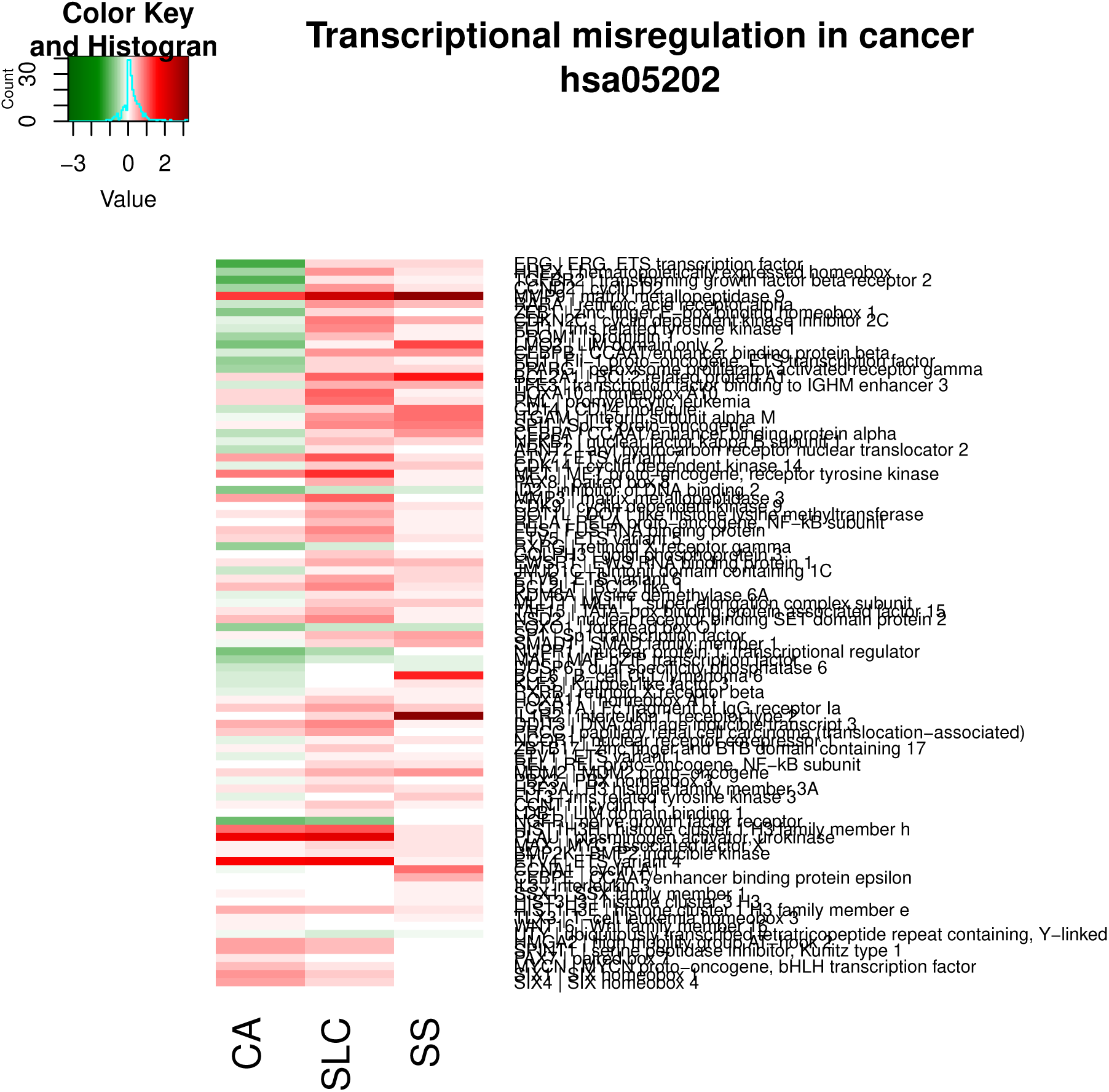

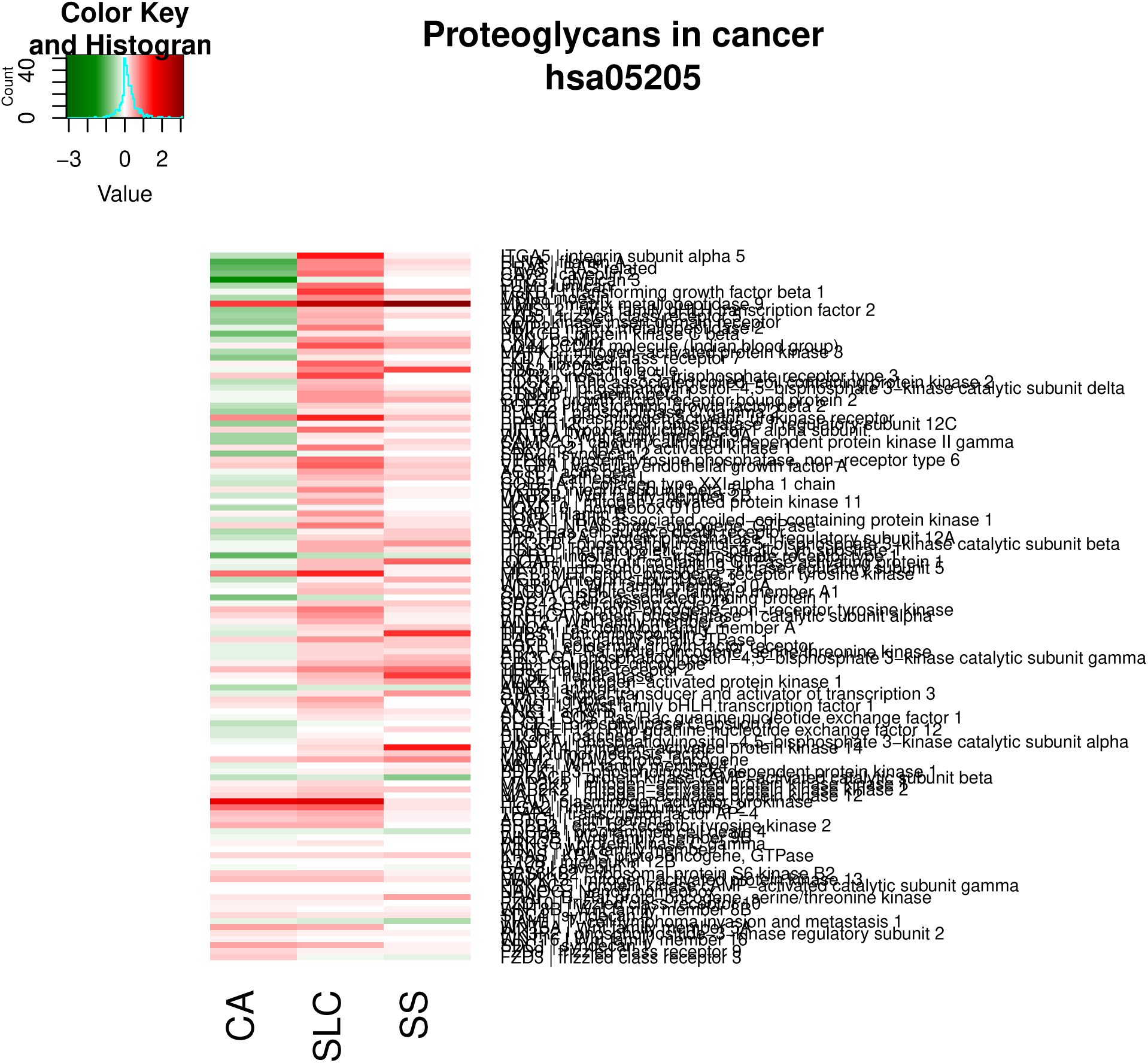

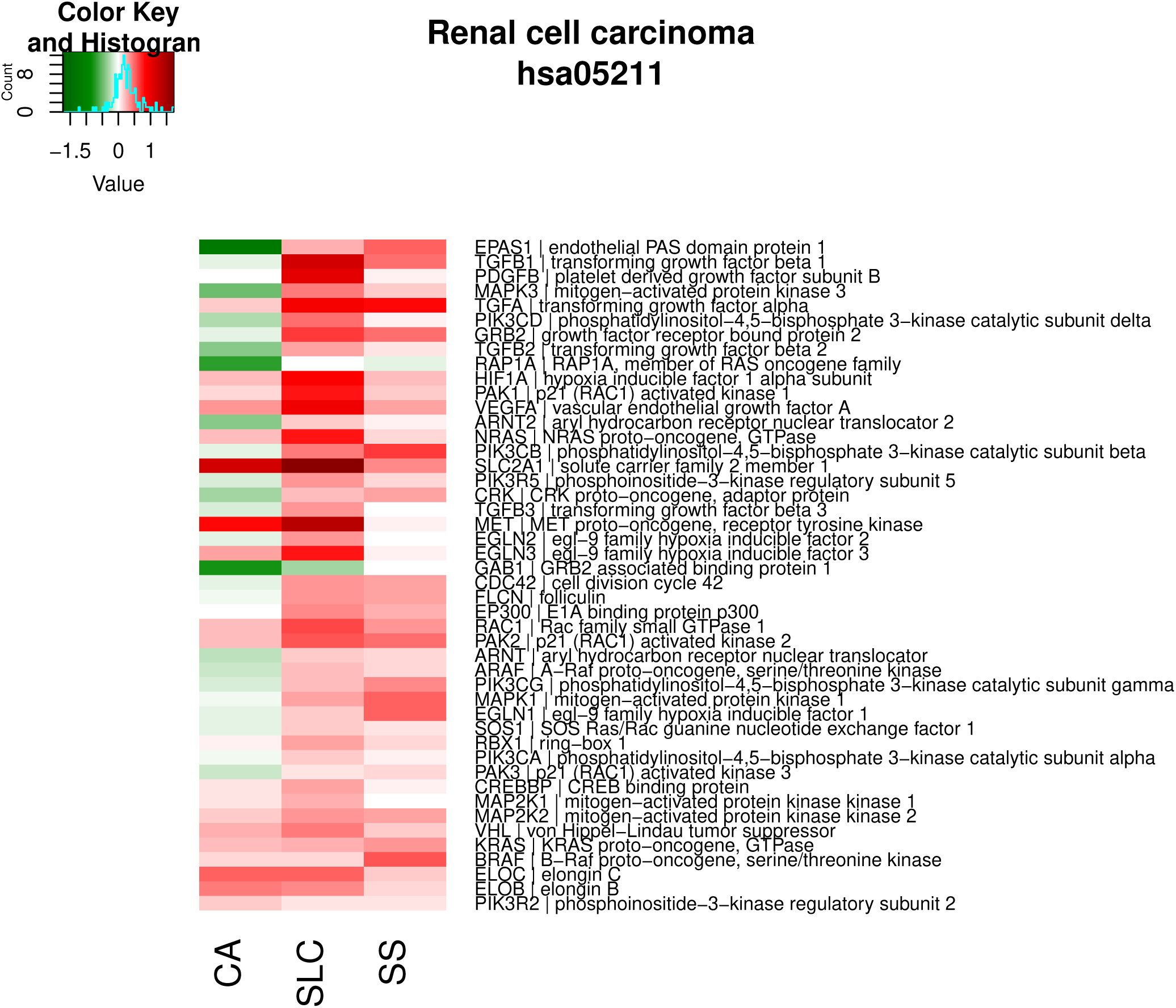

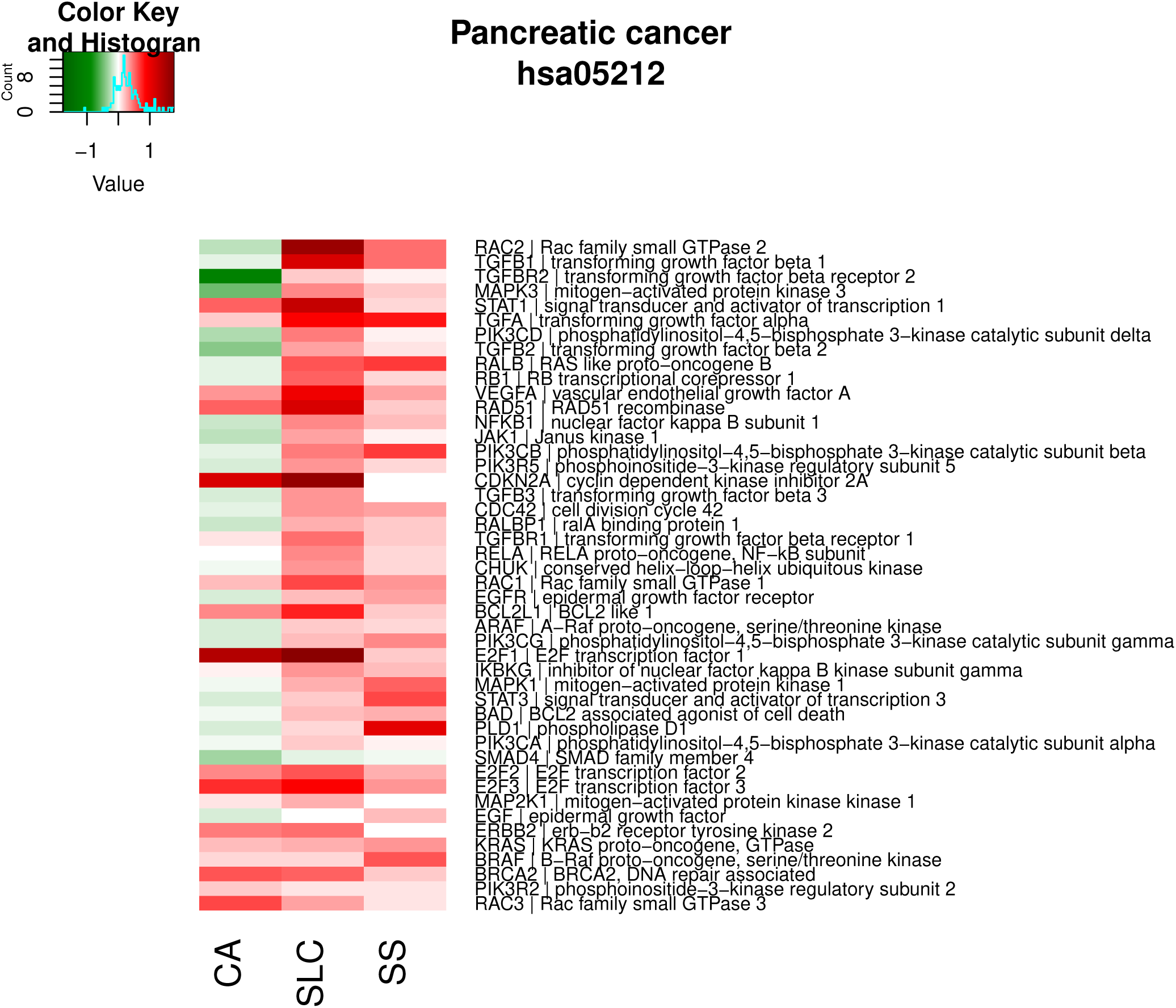

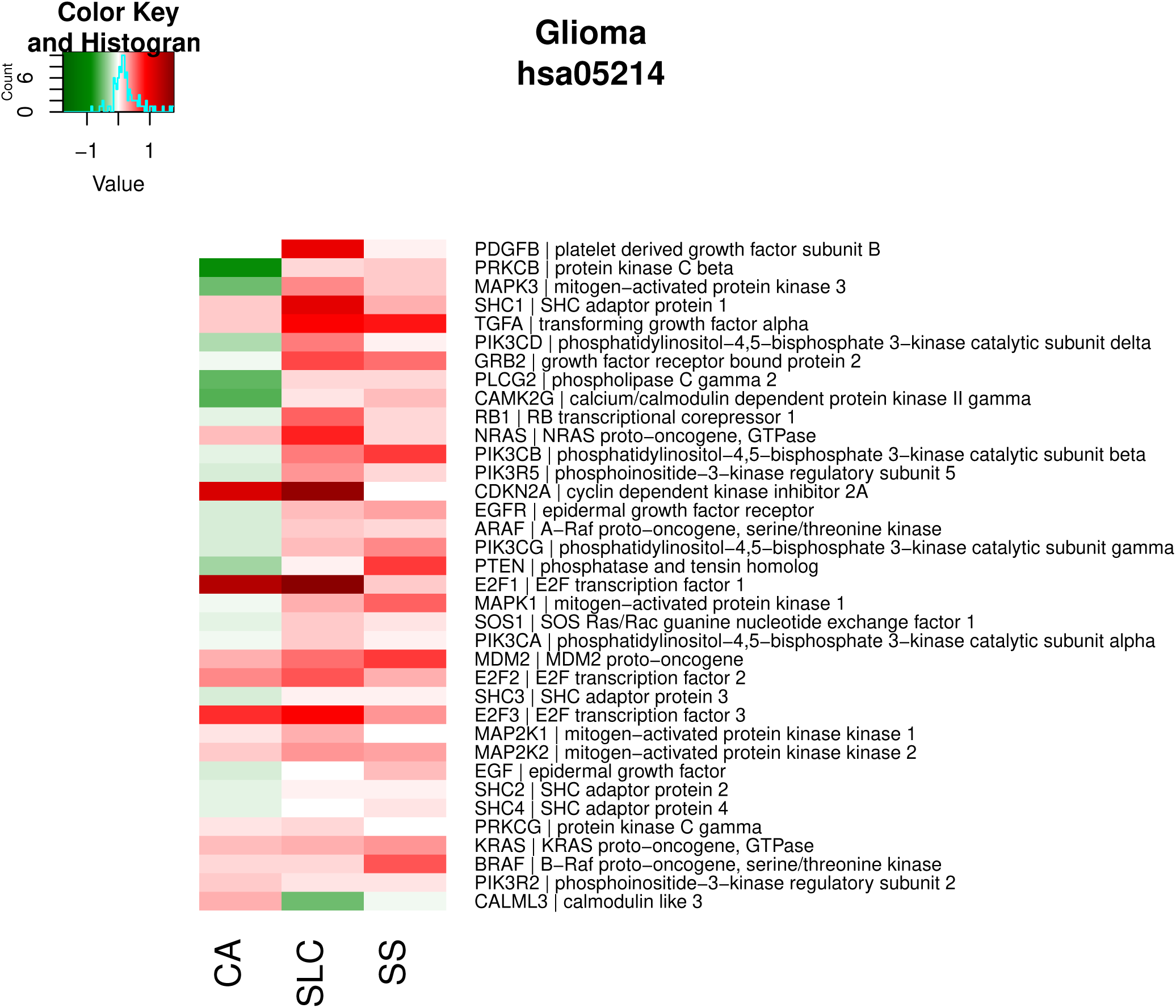

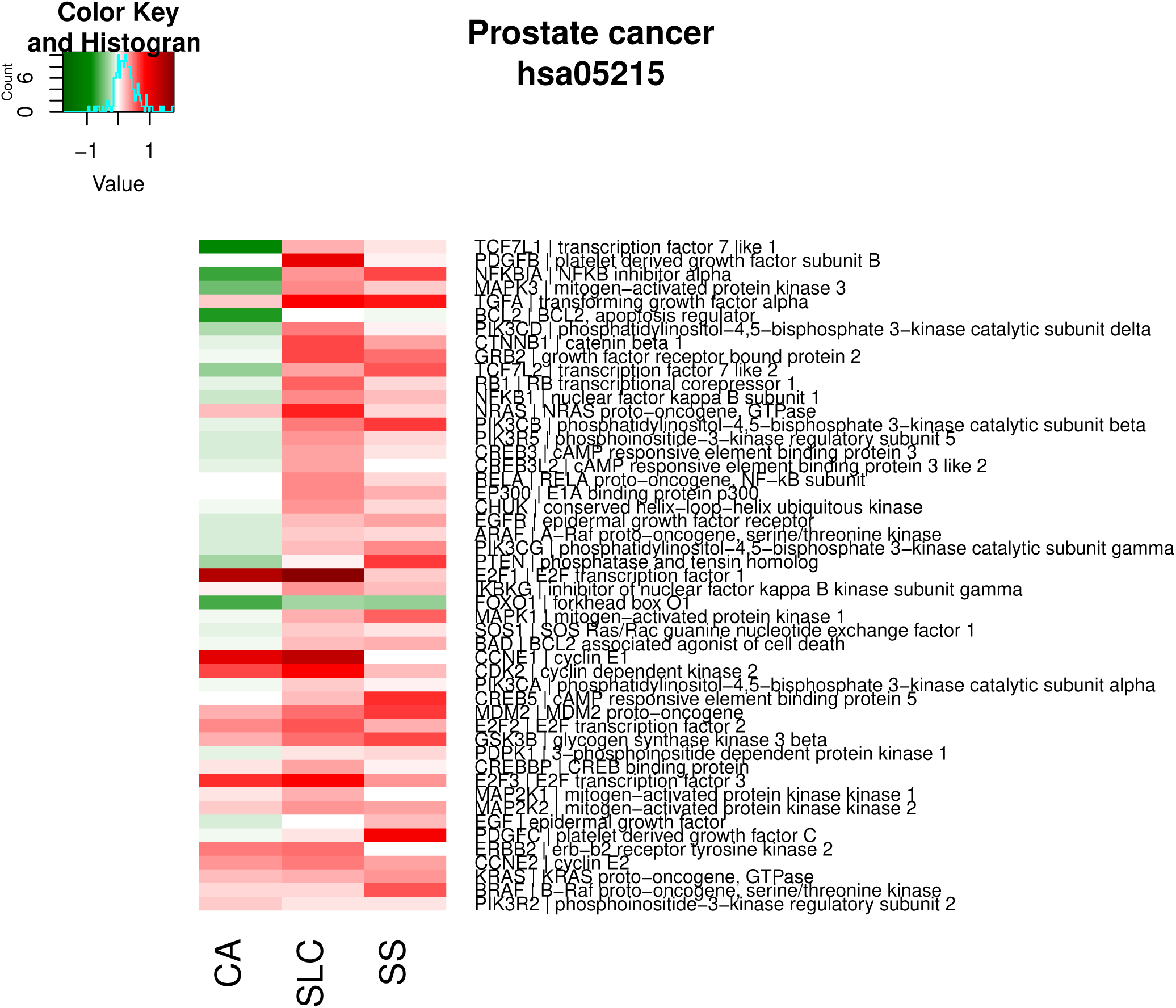

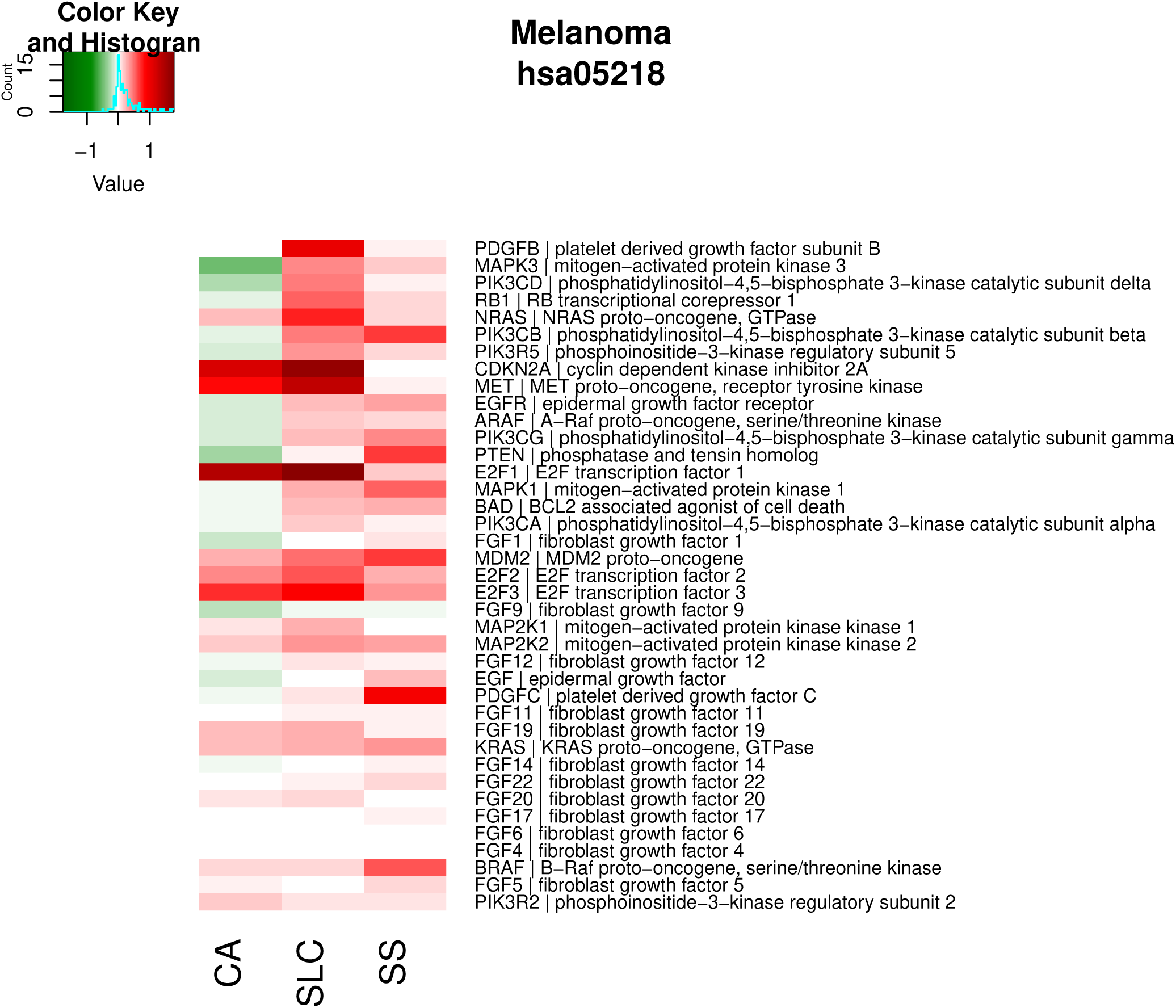

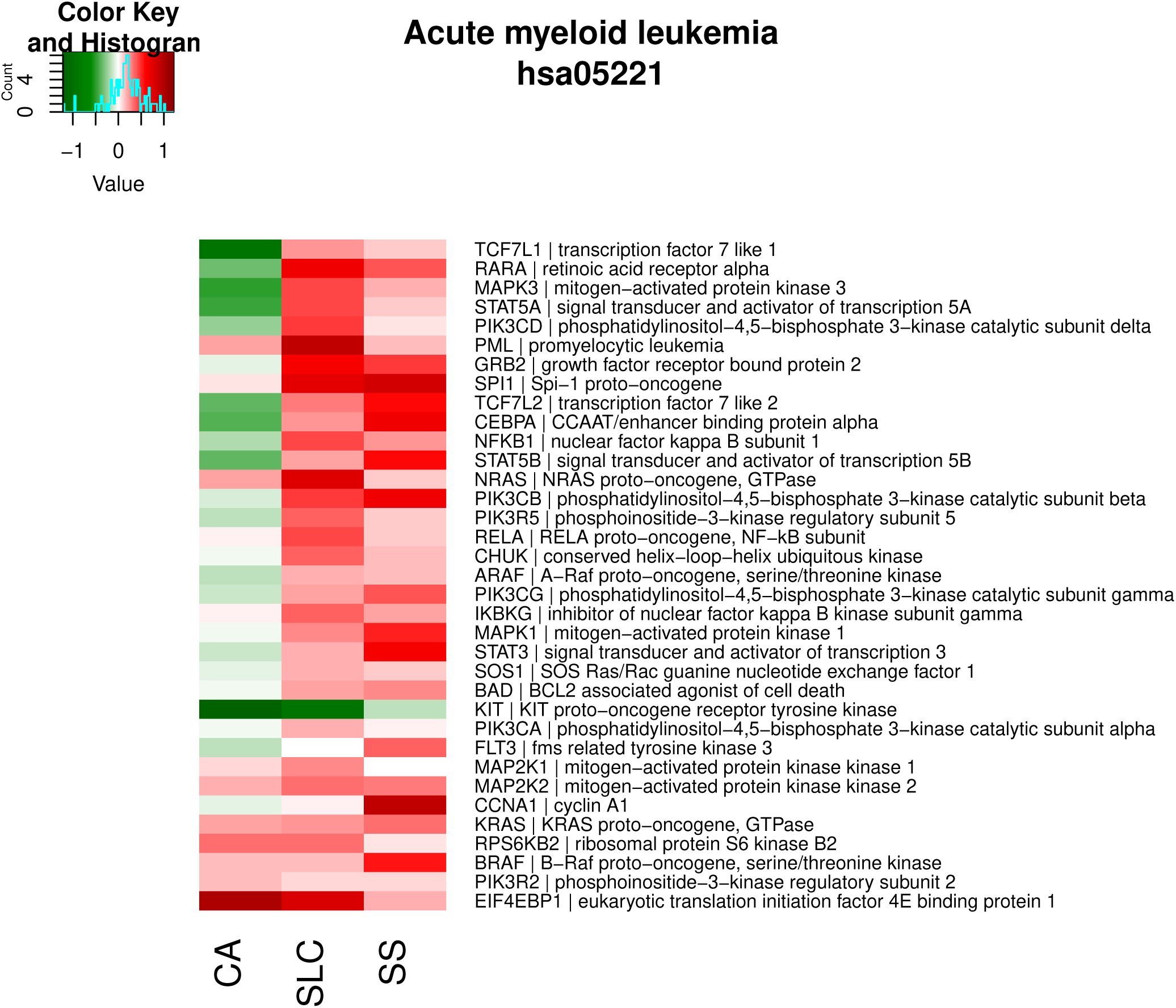

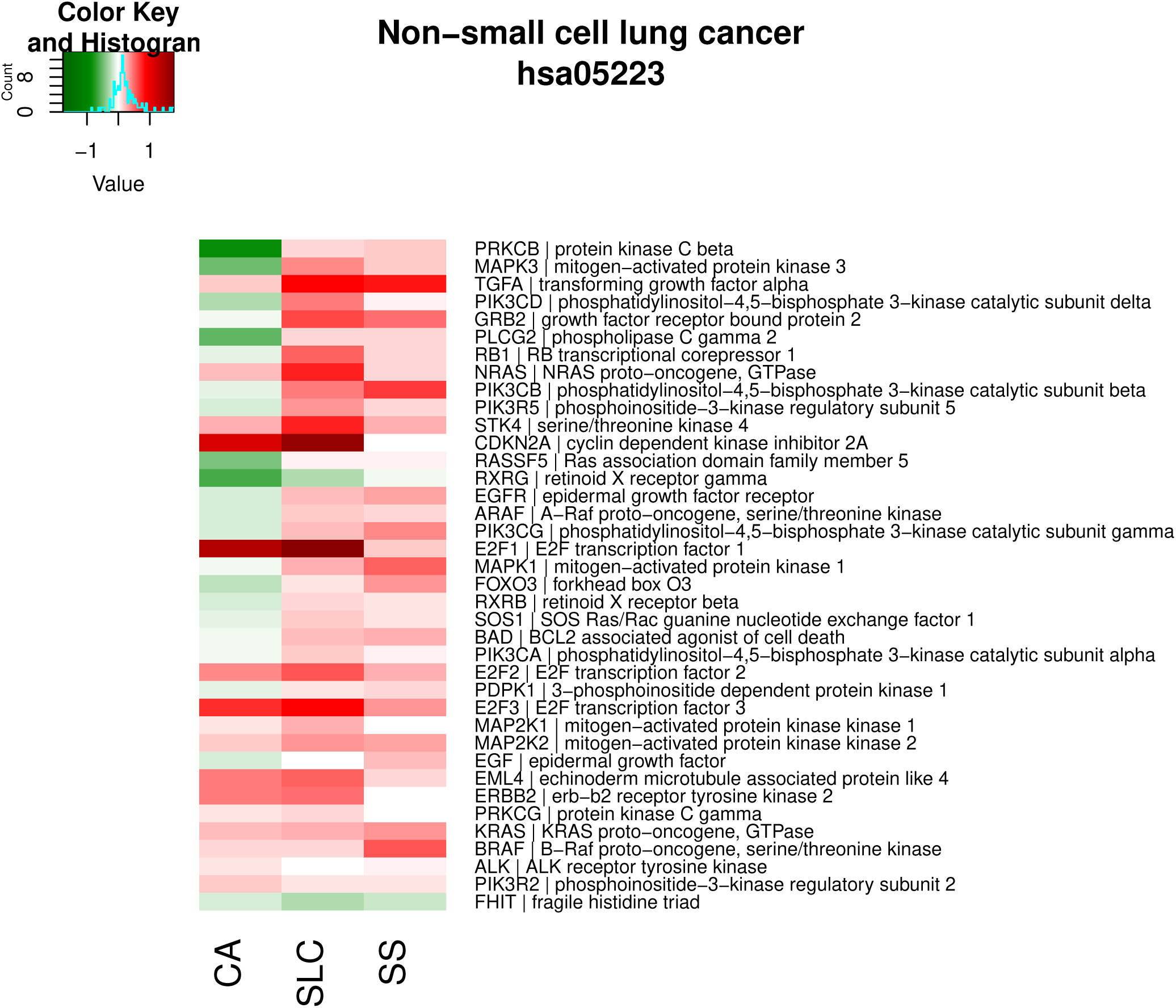

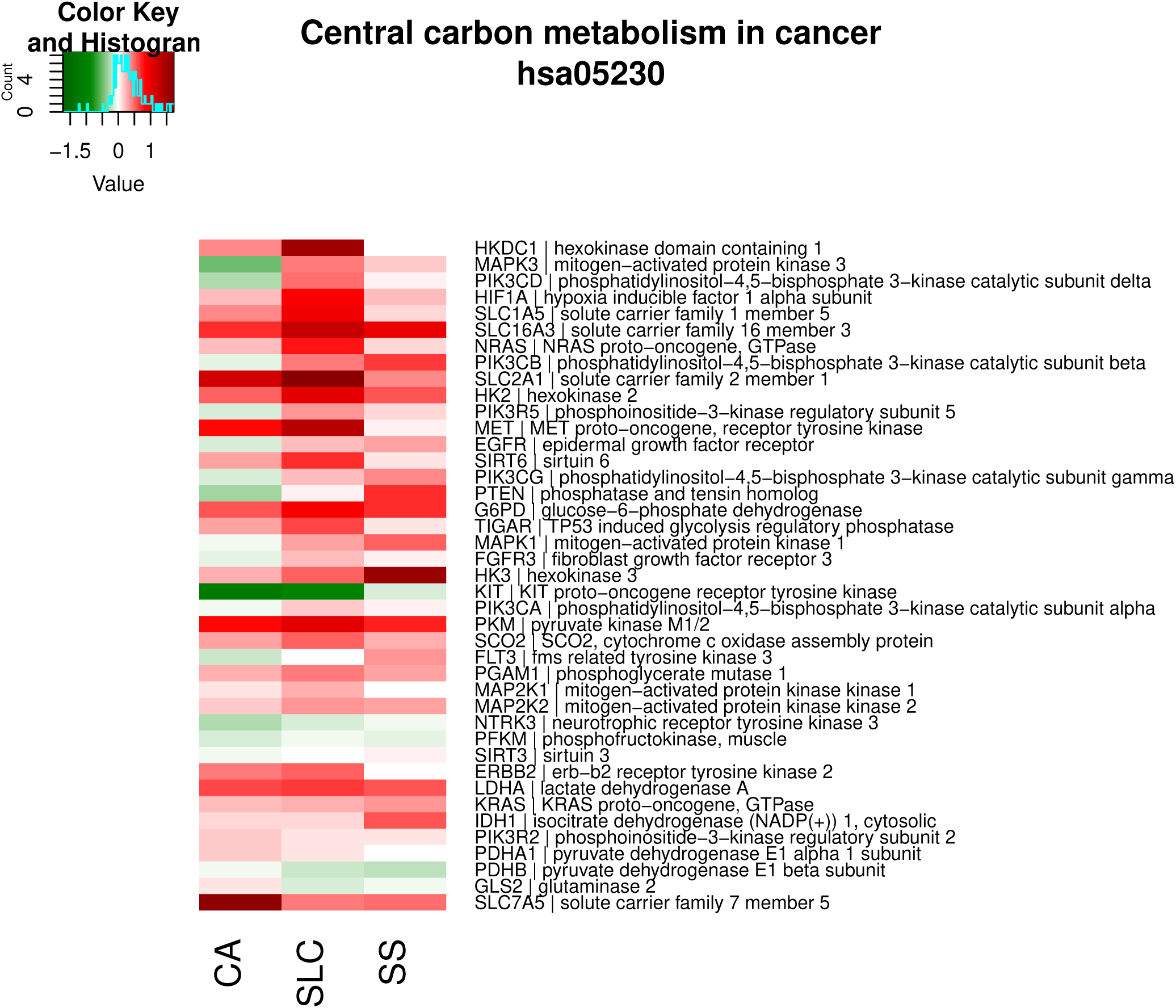

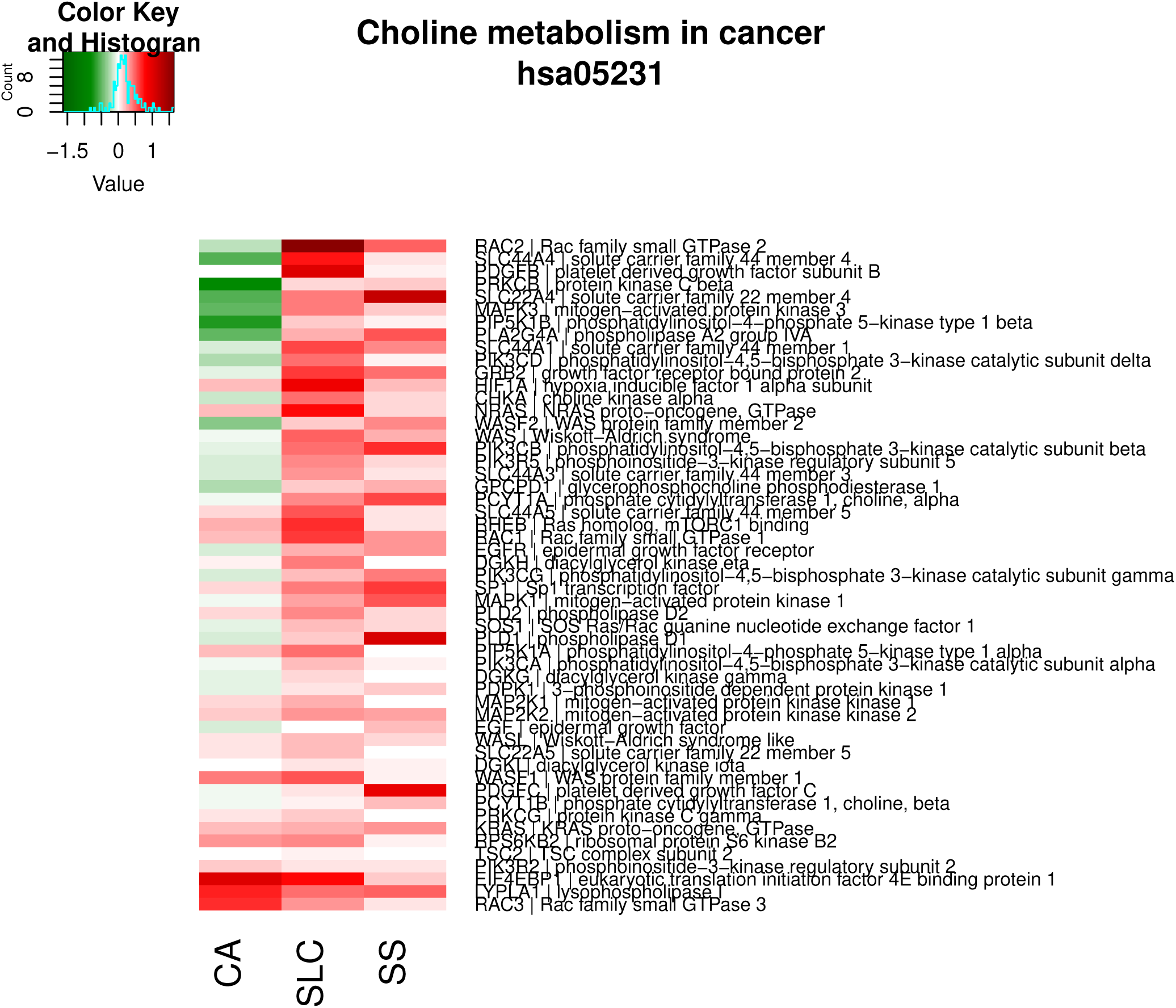

#### Carbohydrate metabolism

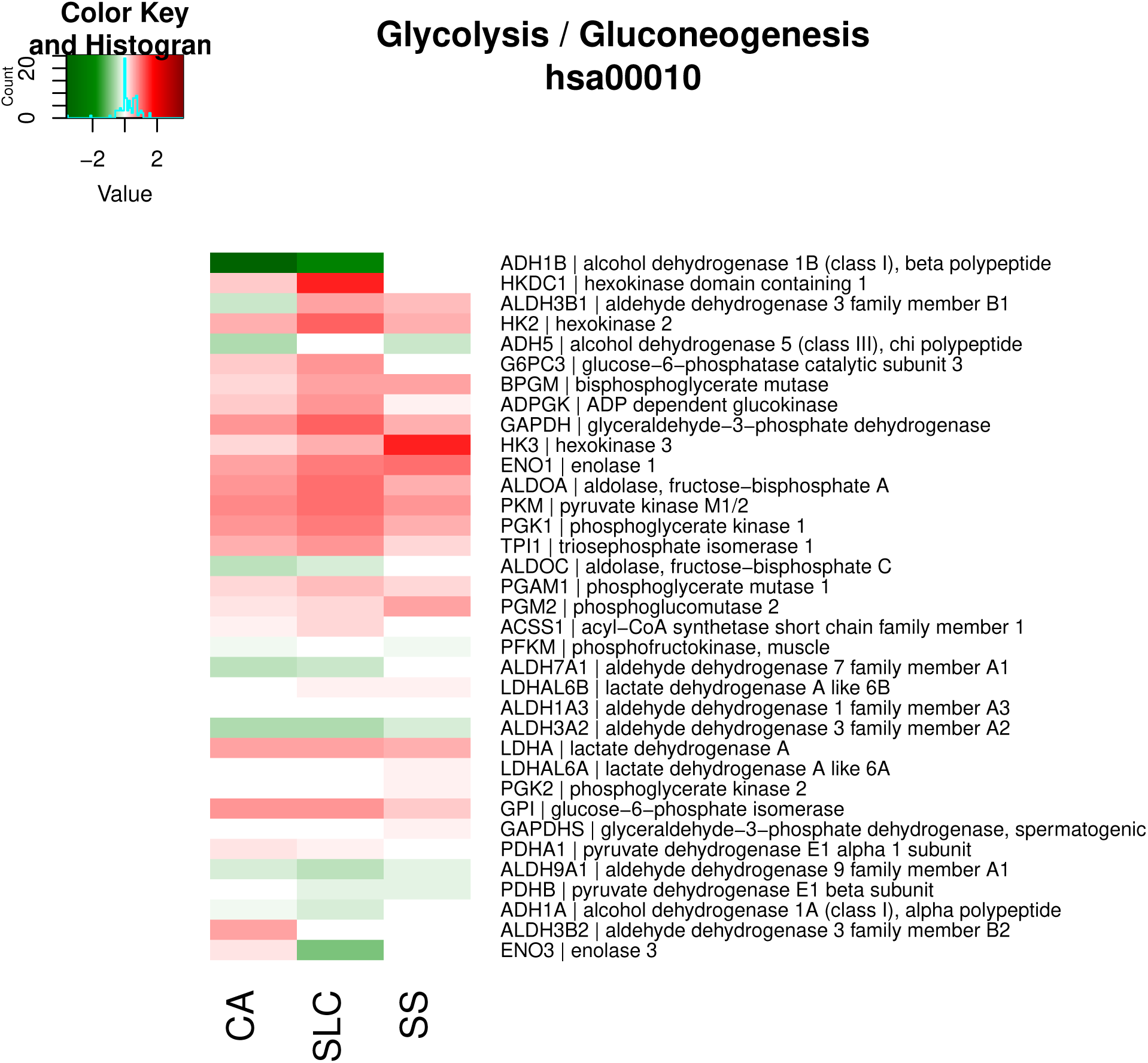

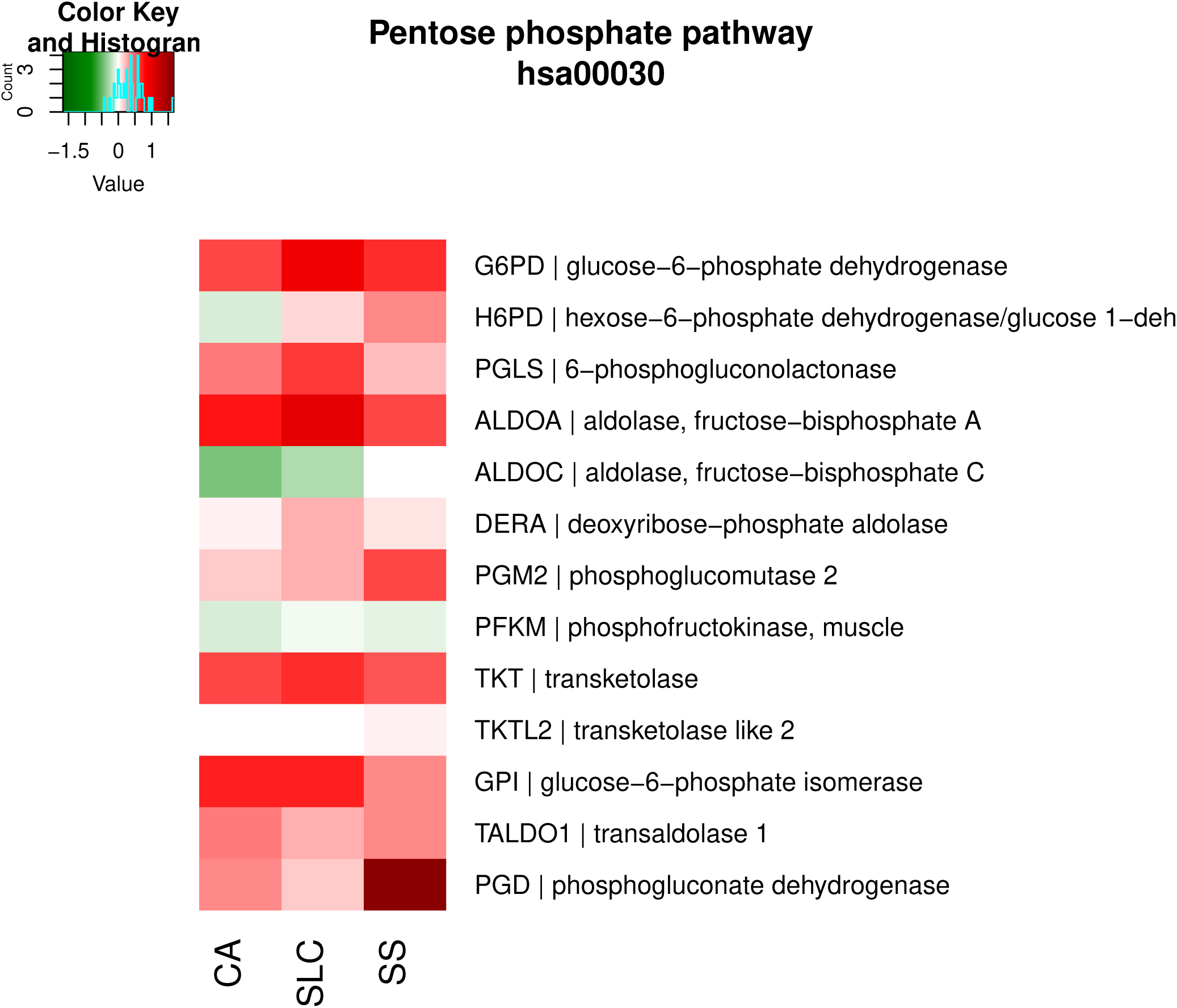

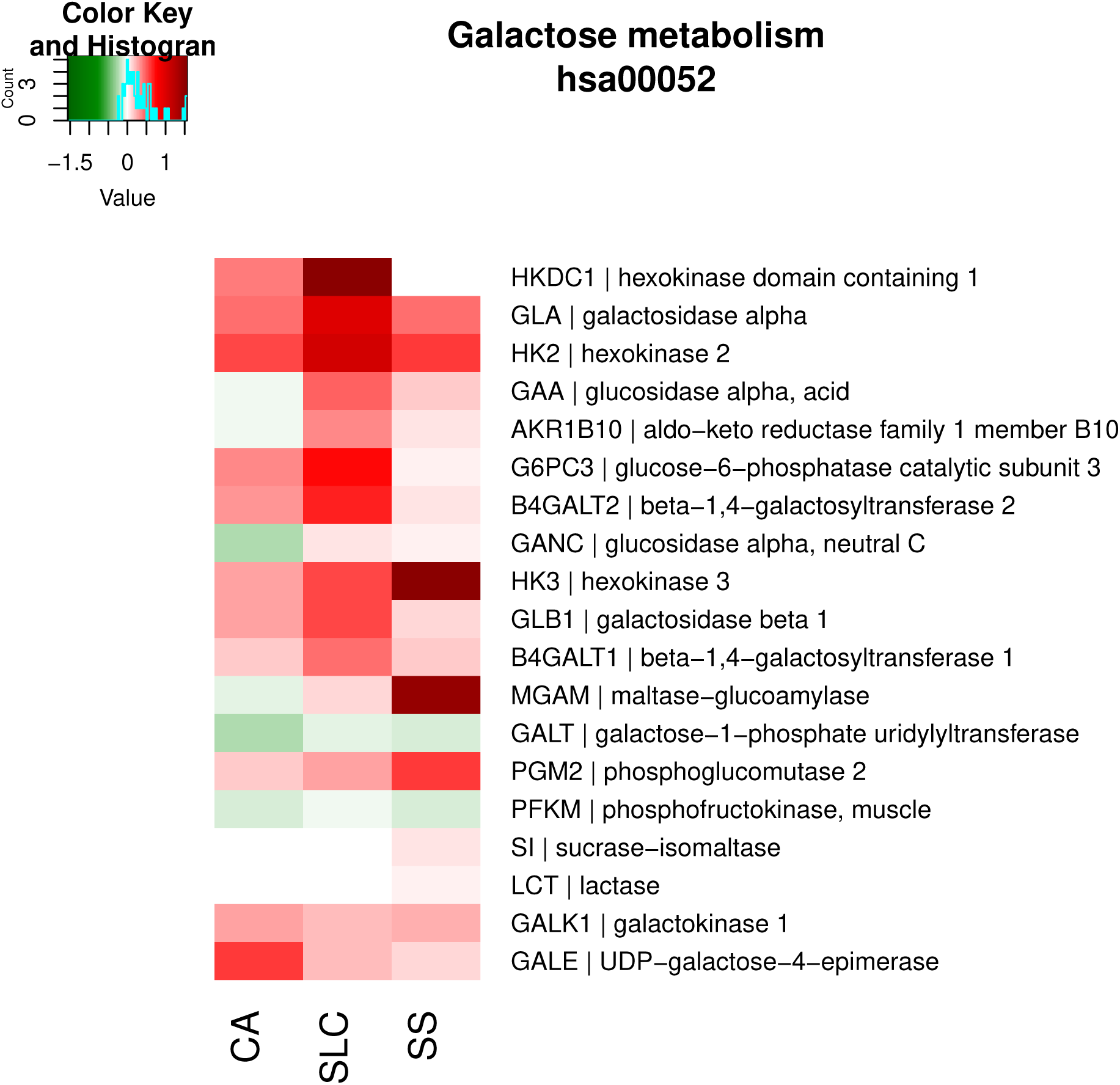

#### Endocrine system

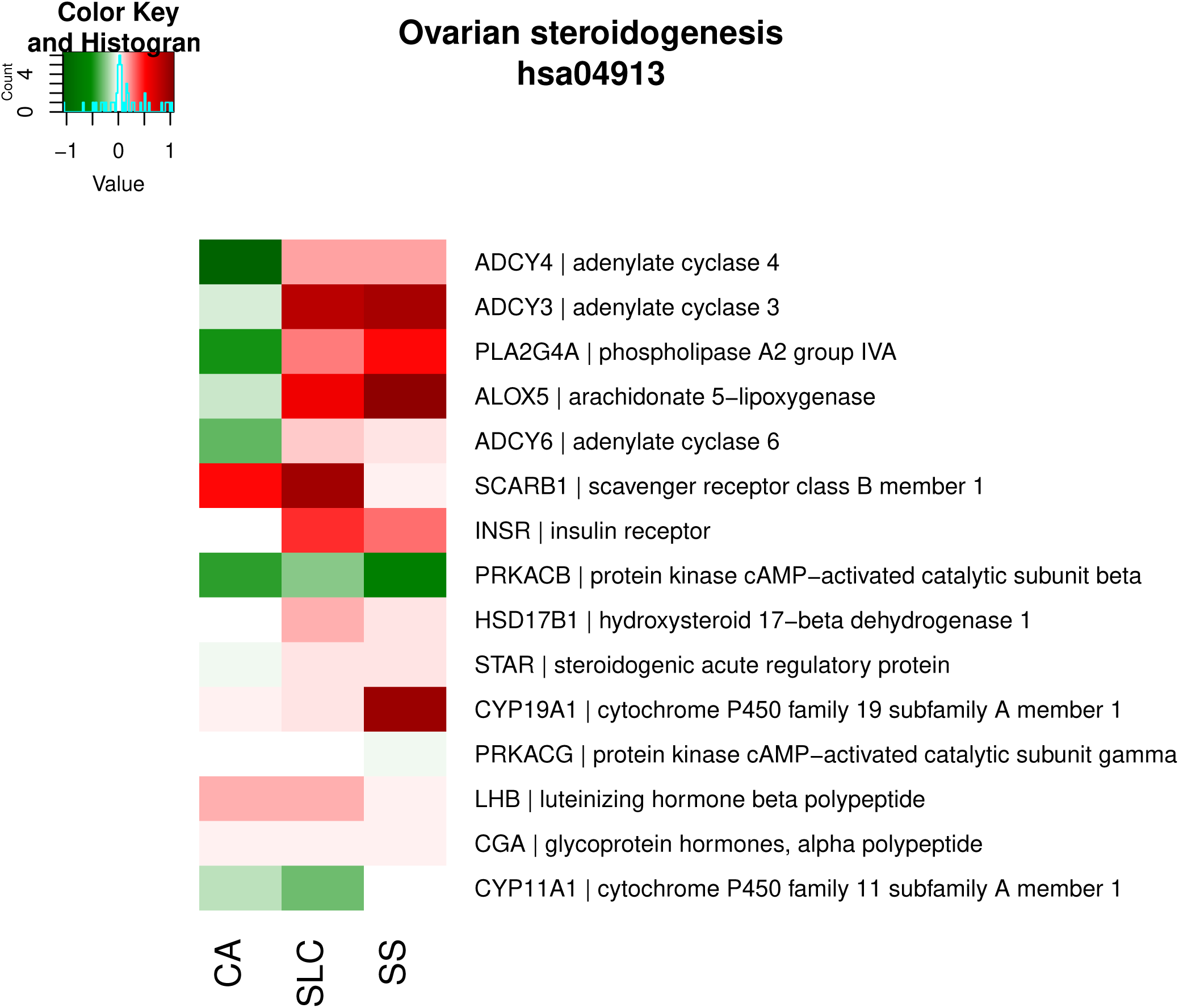

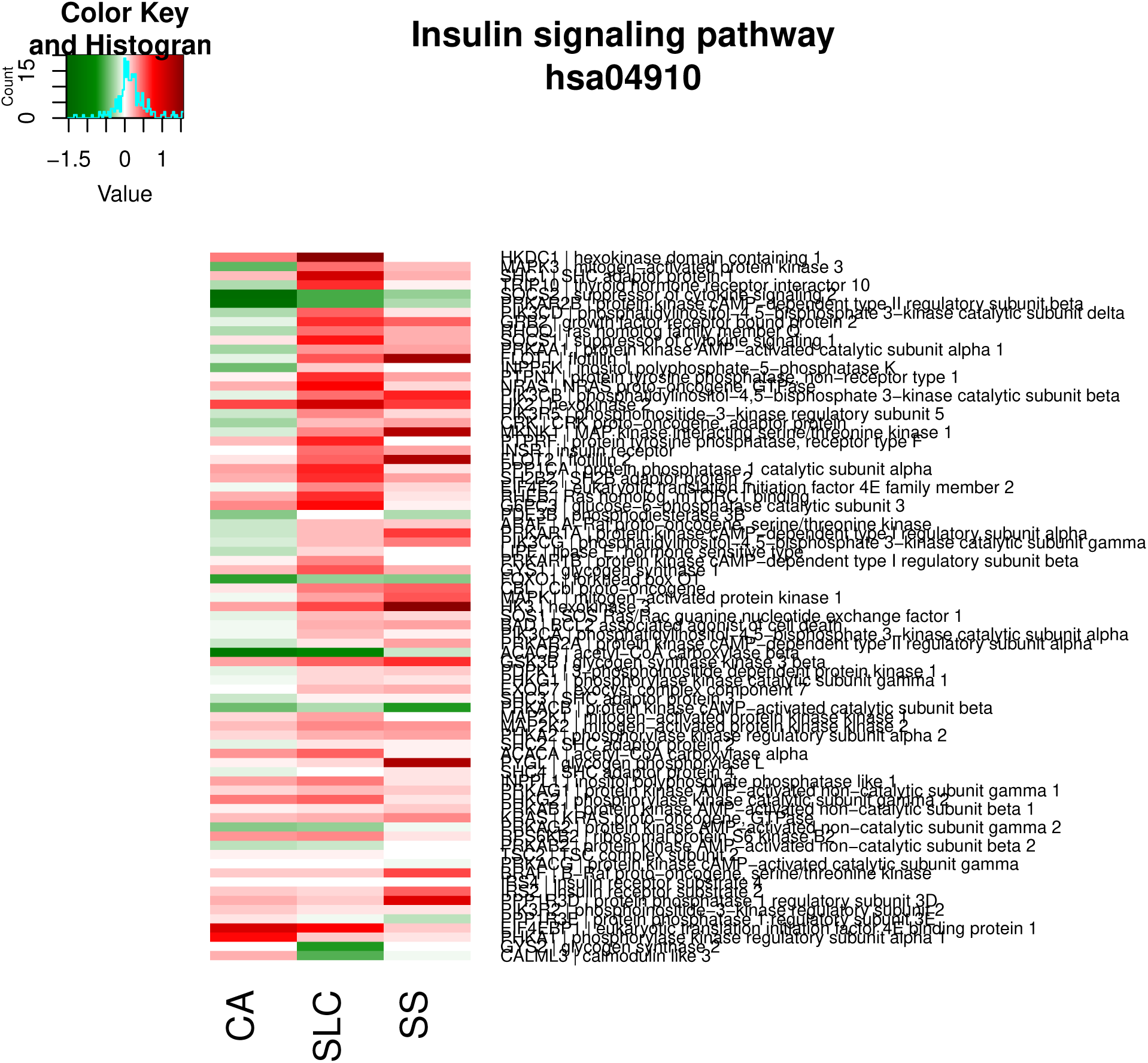

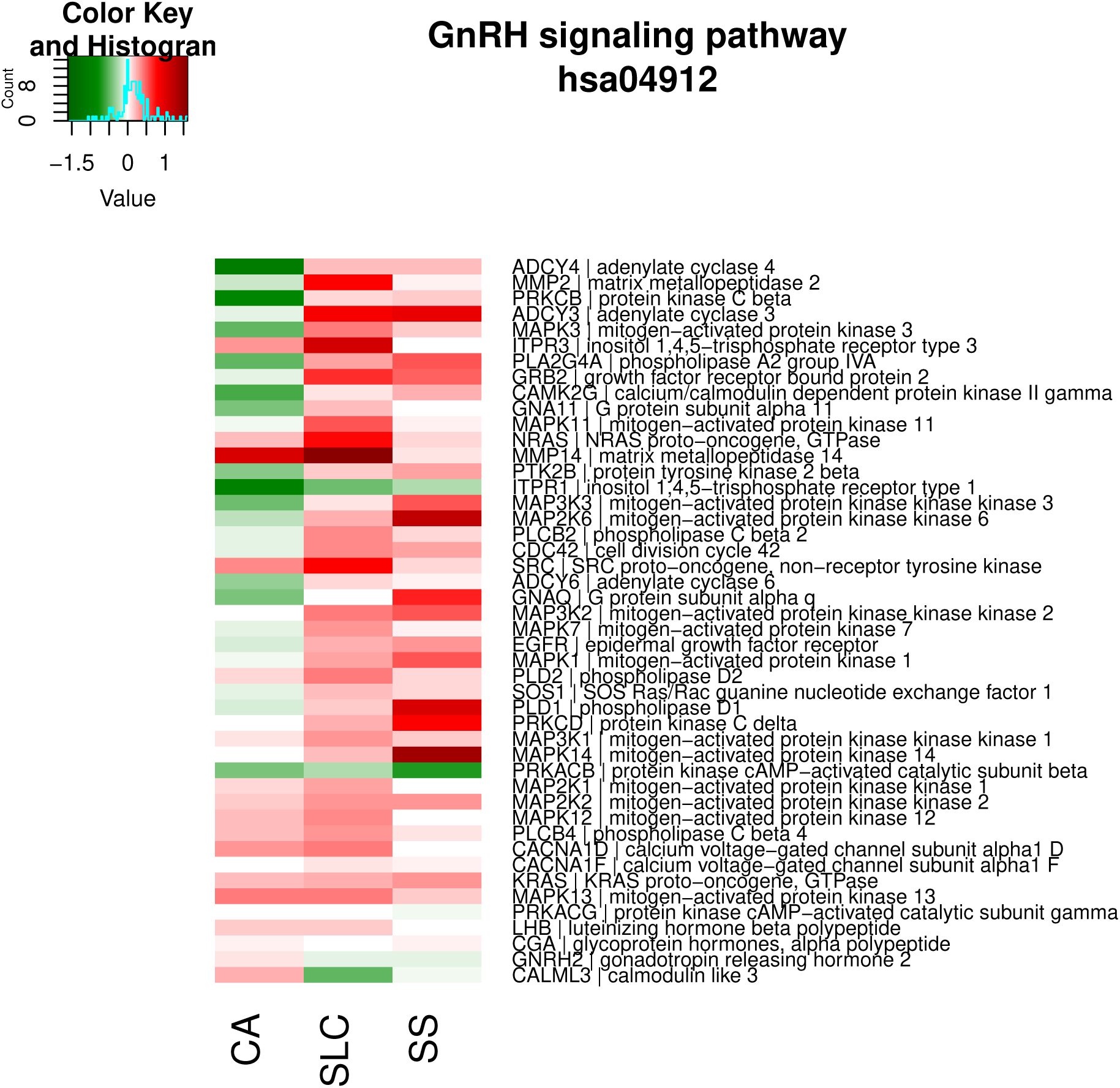

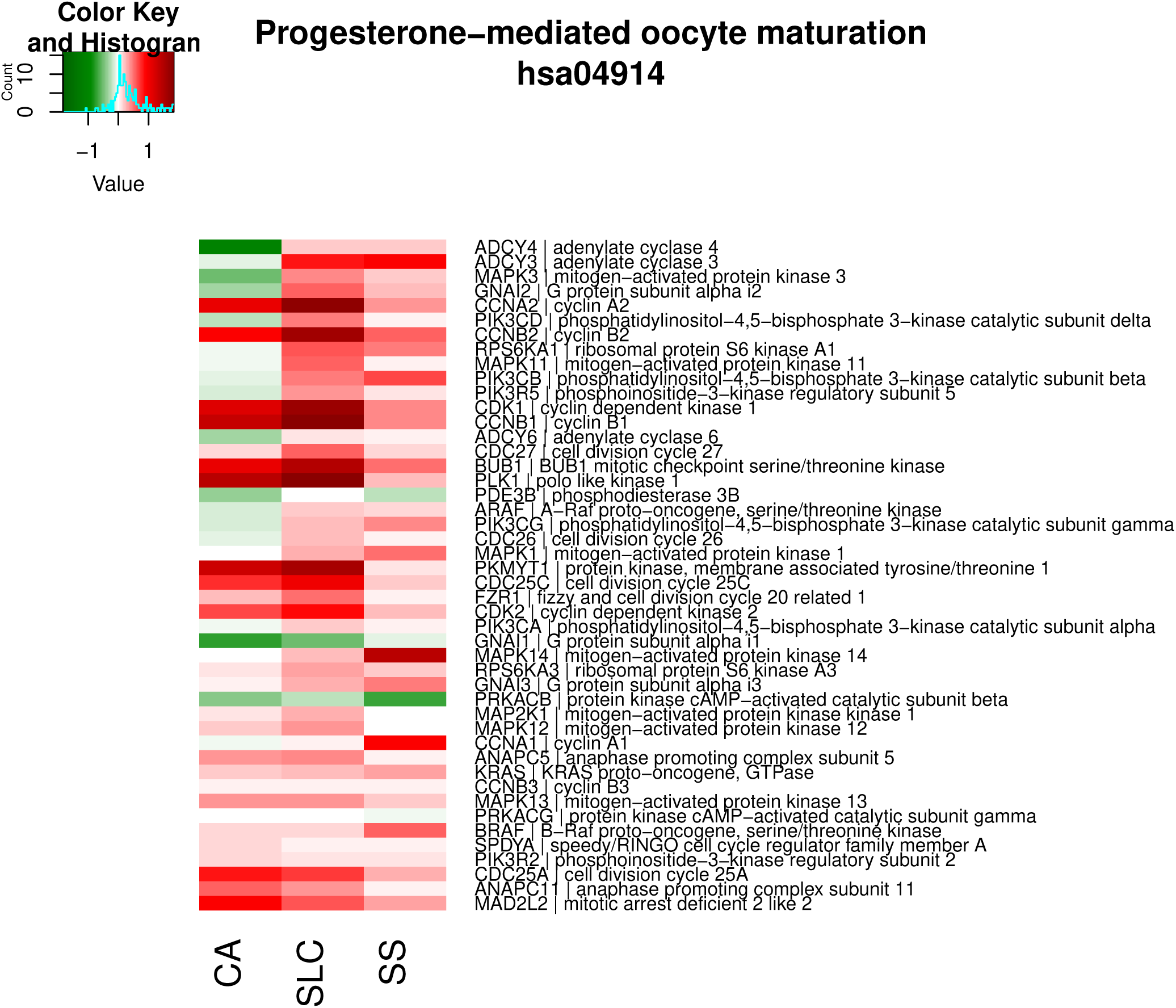

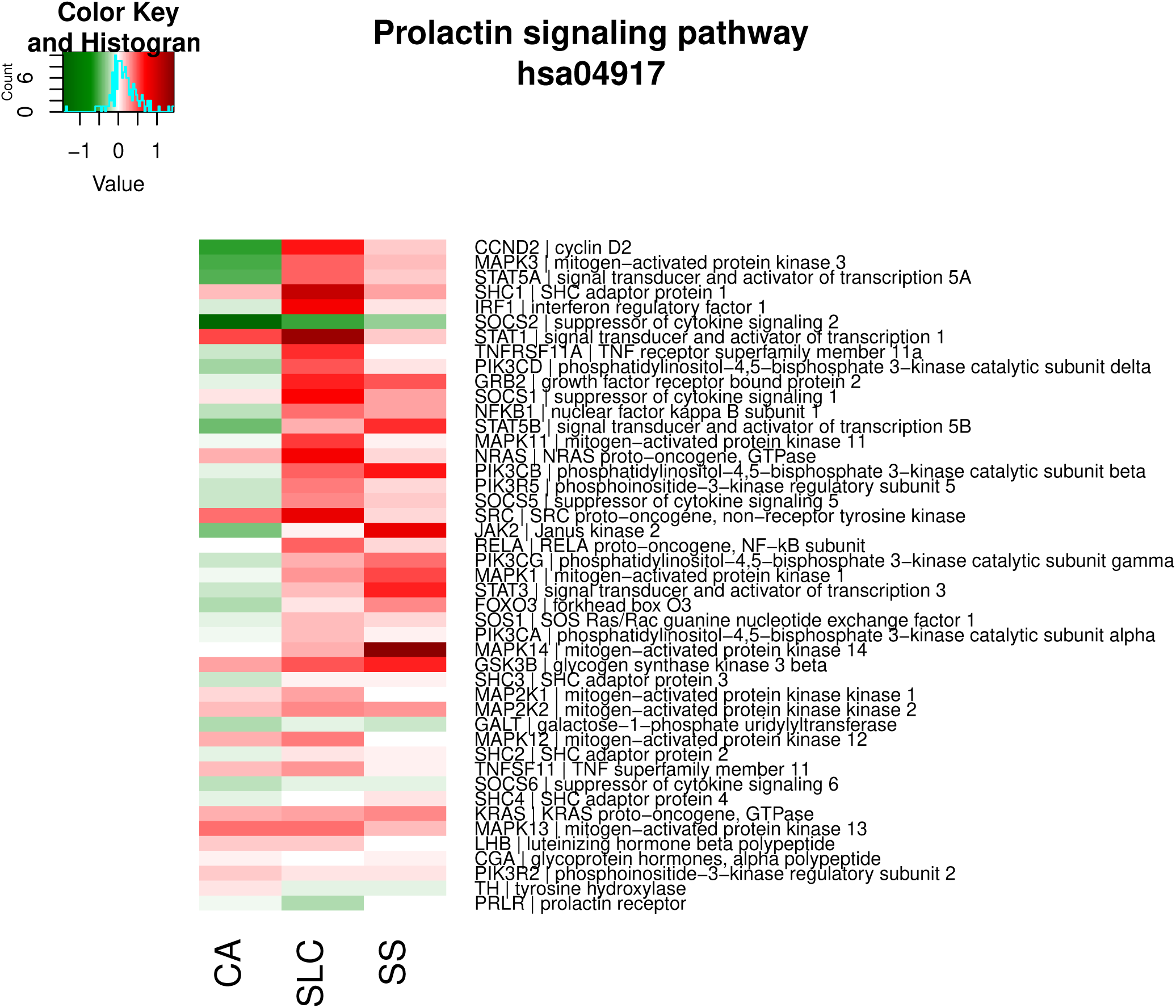

#### Immune system

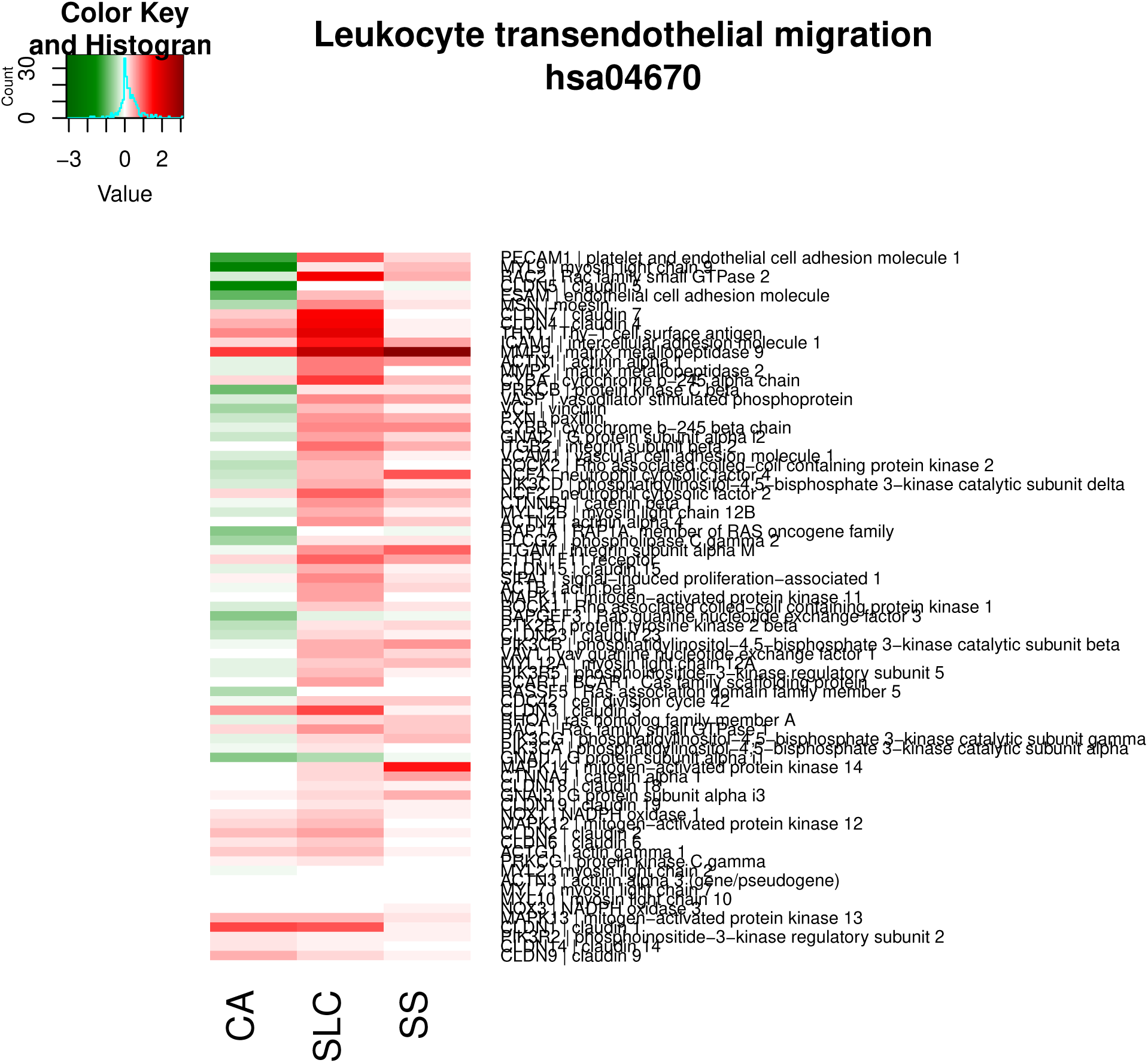

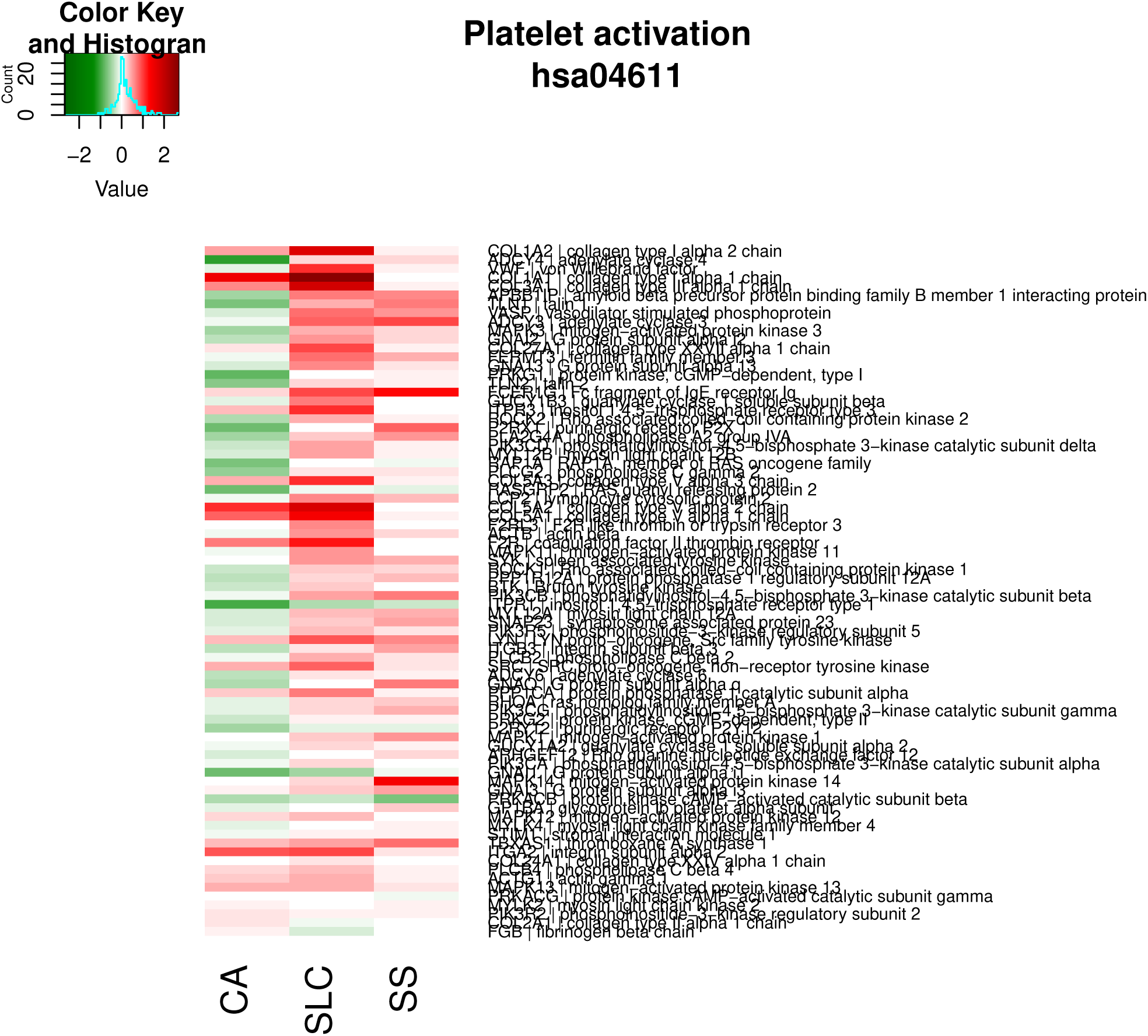

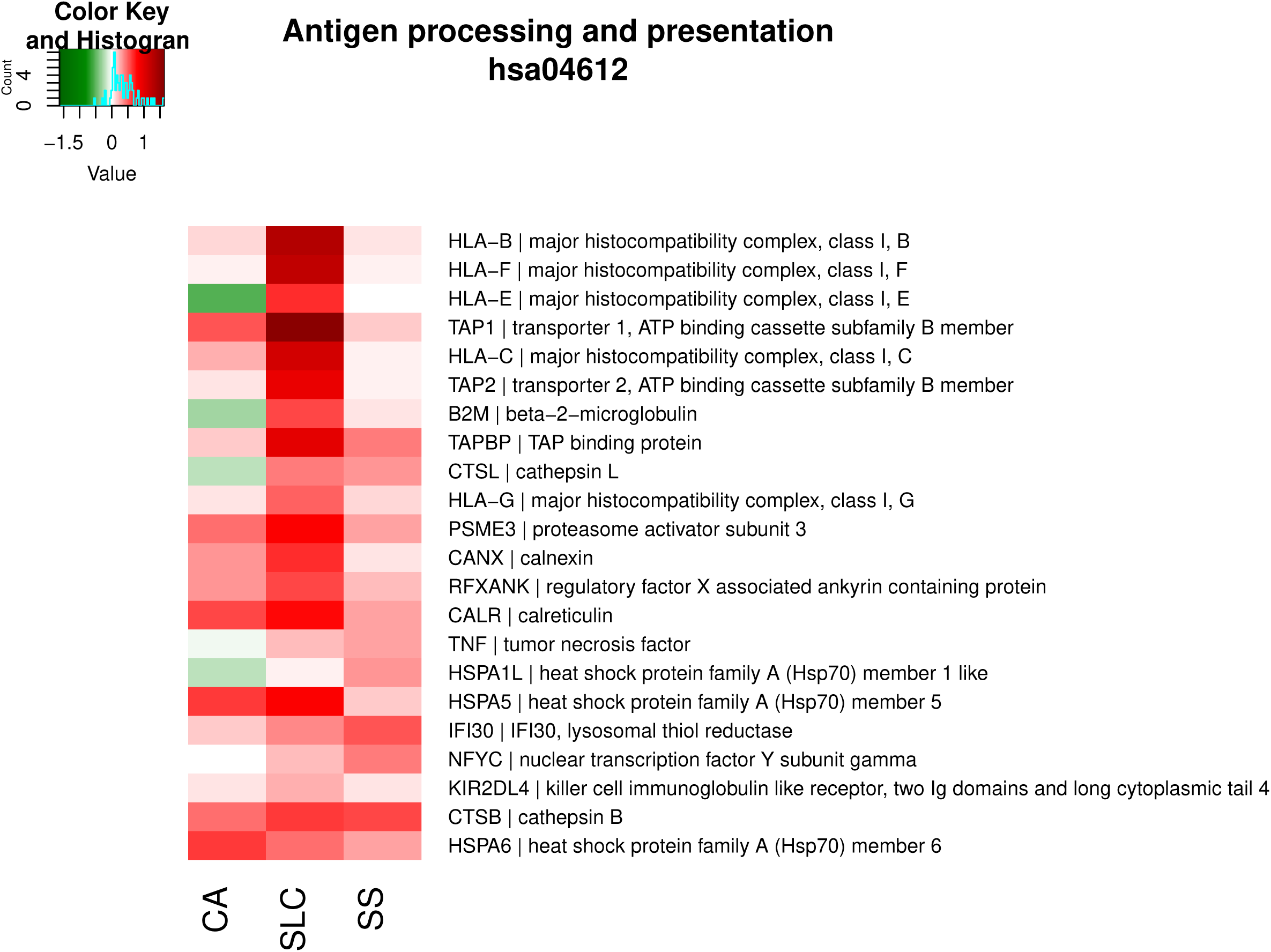

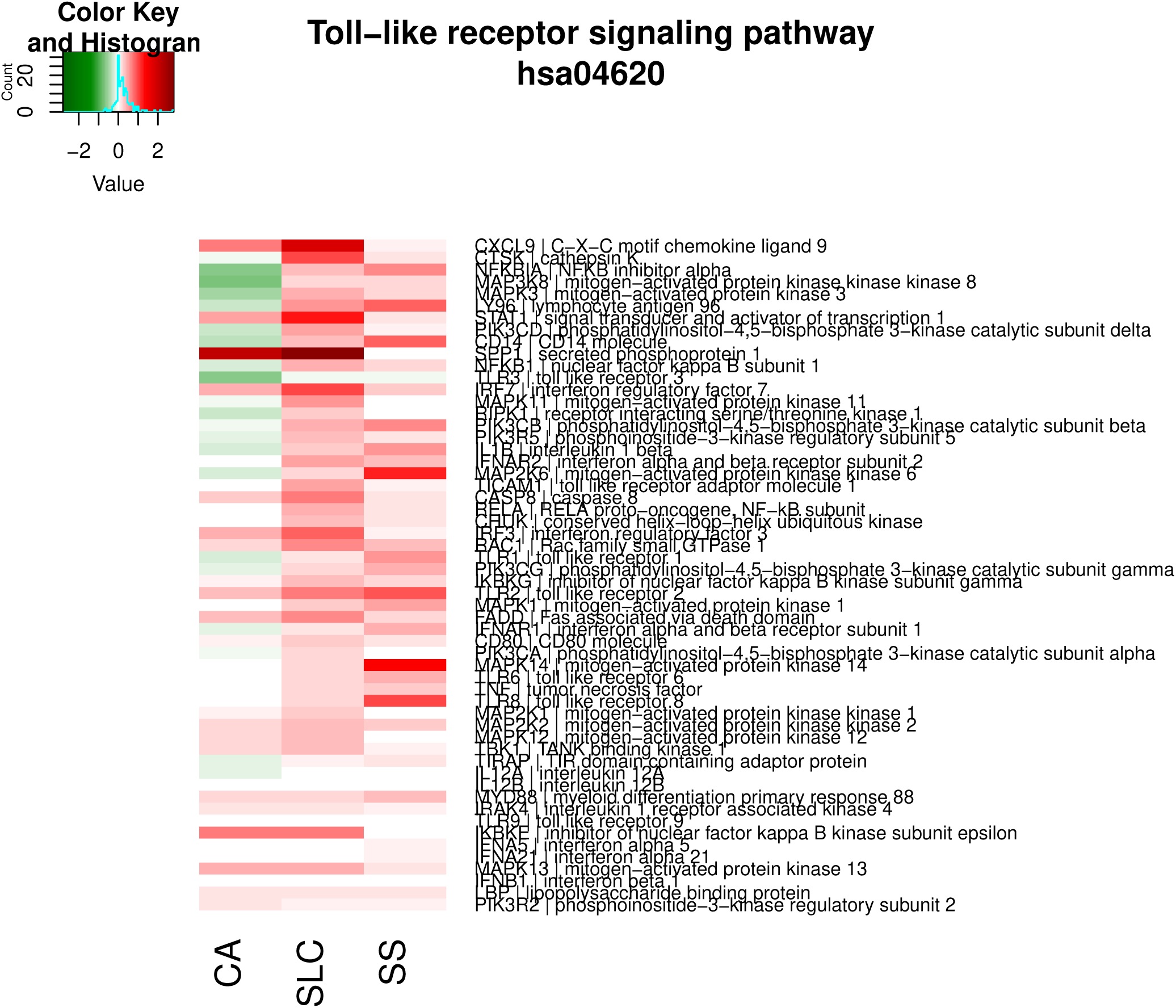

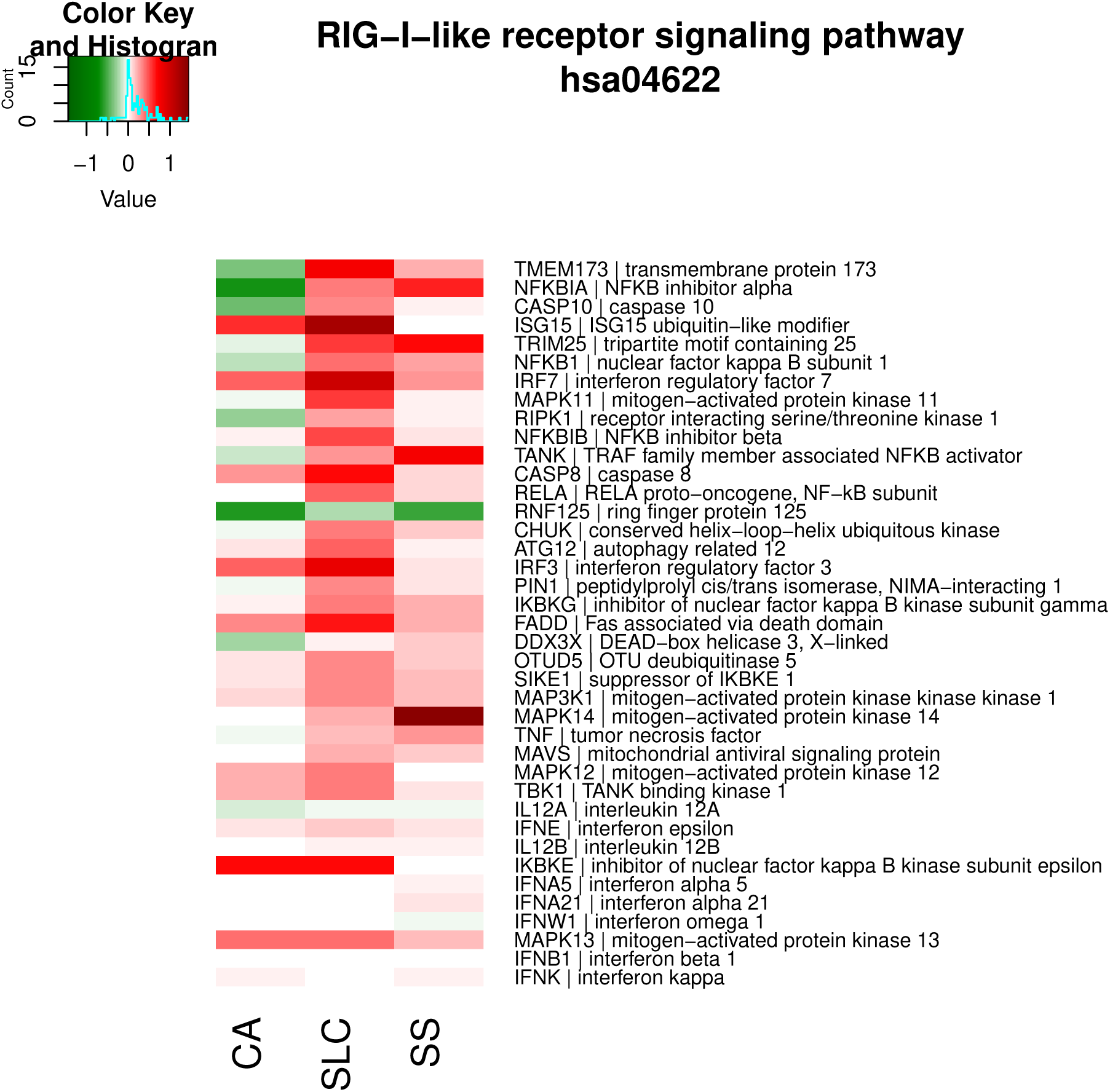

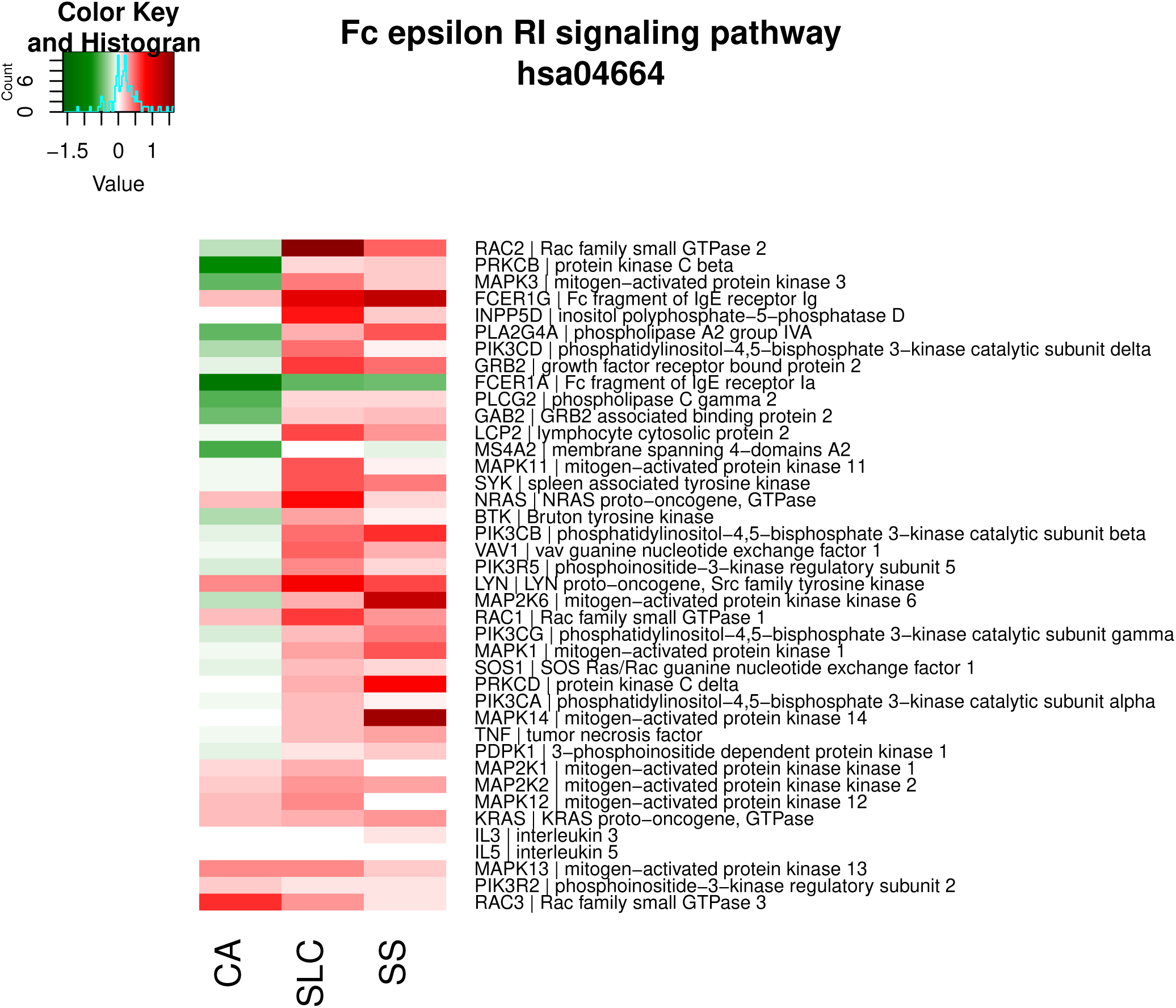

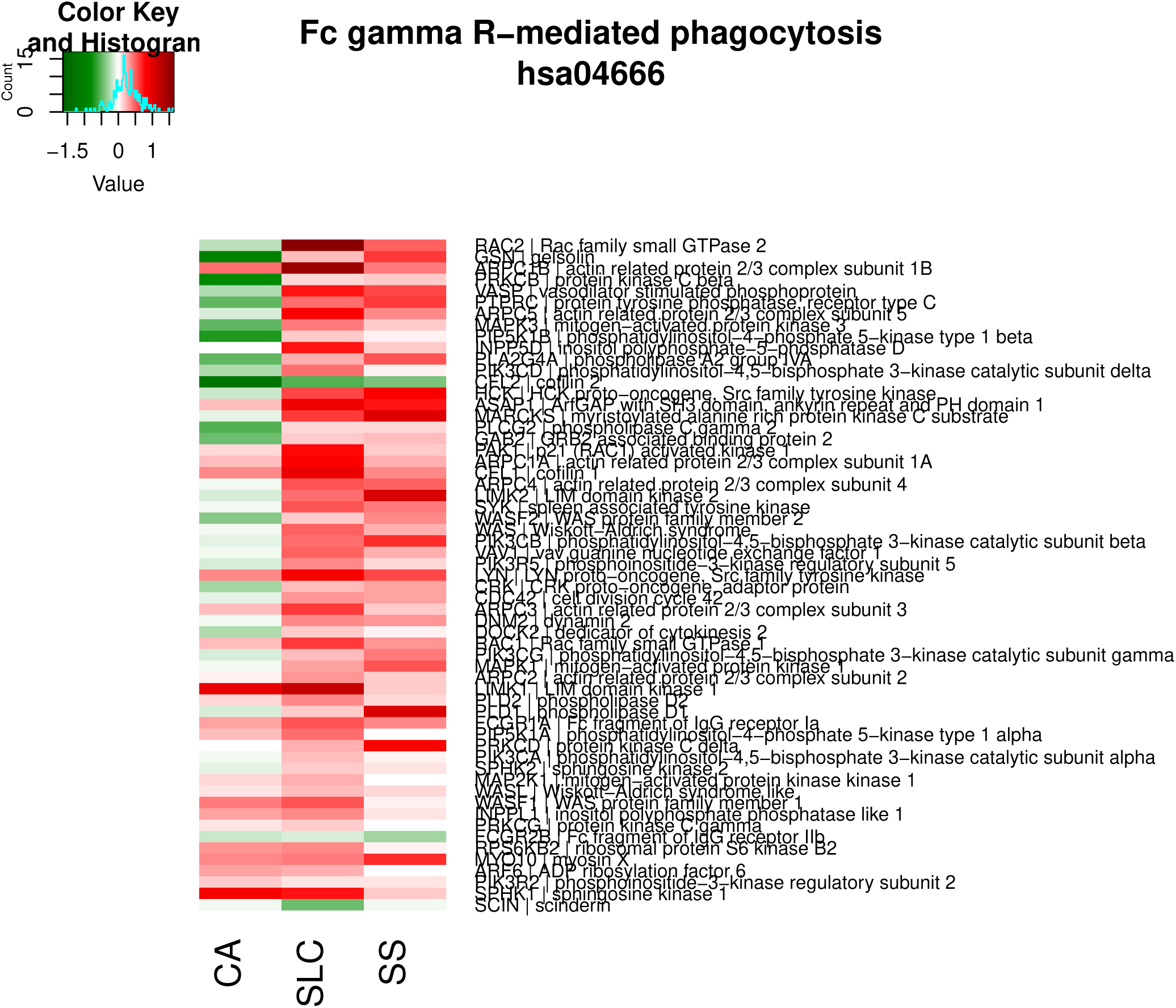

#### Infectious disease

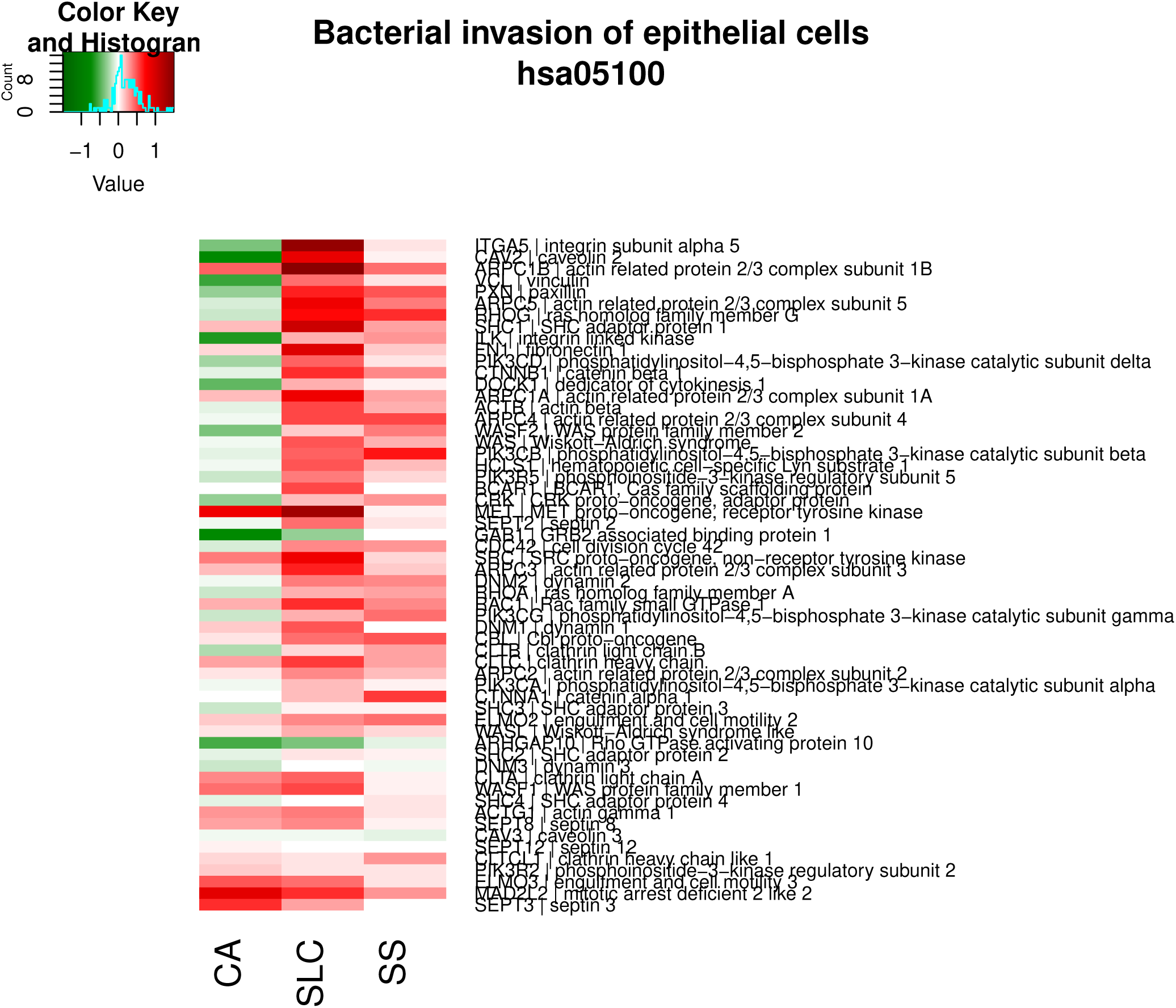

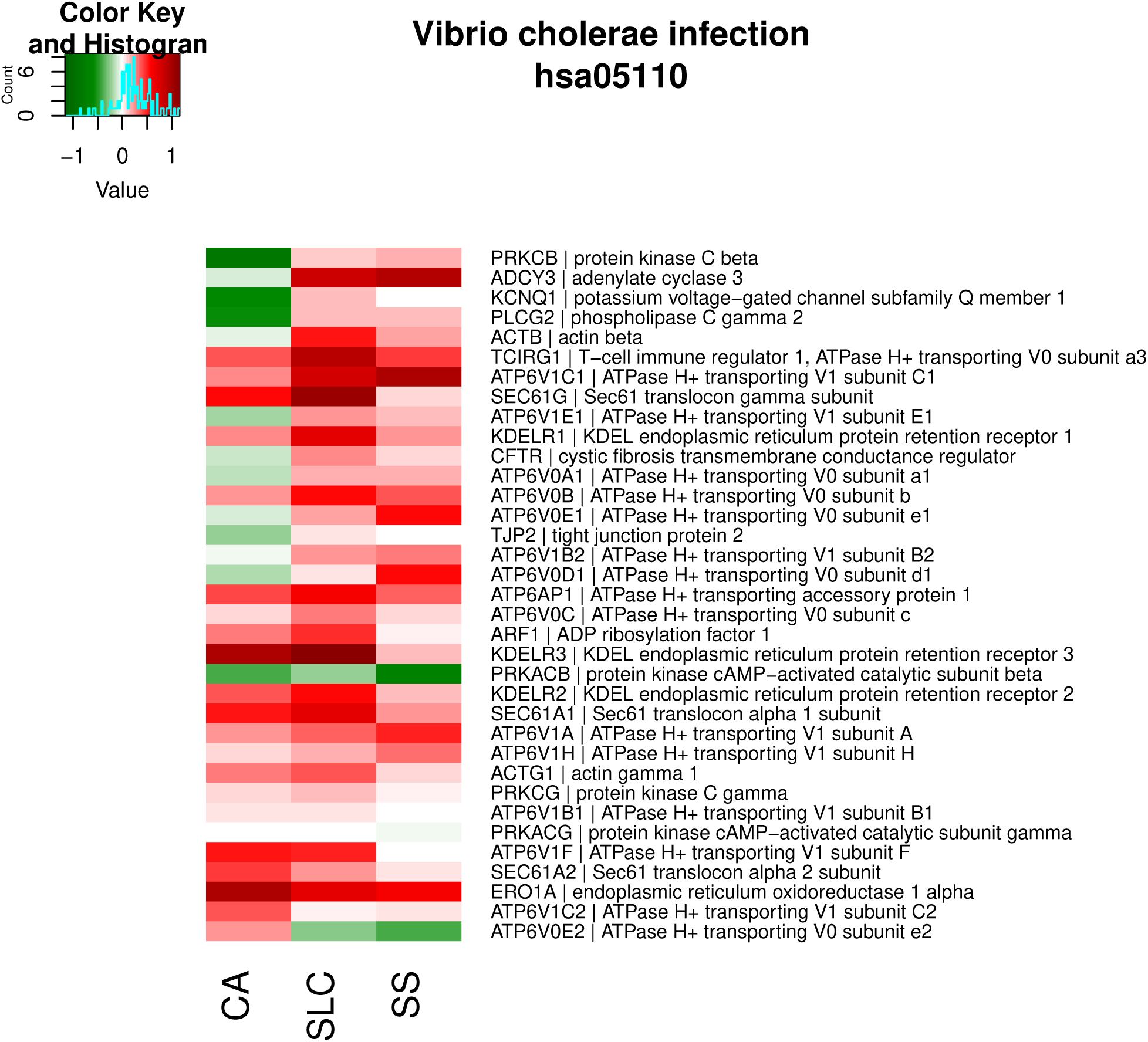

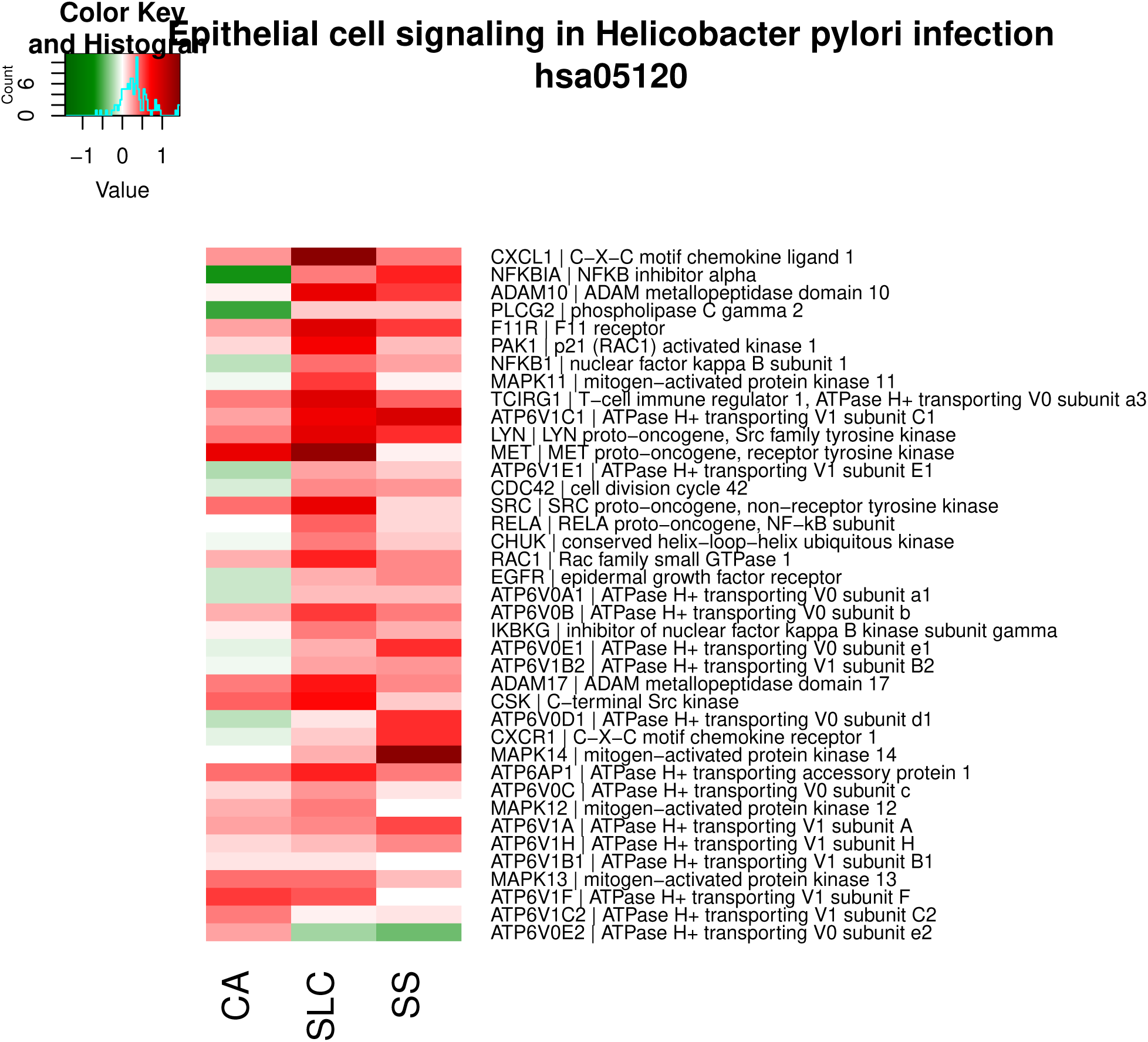

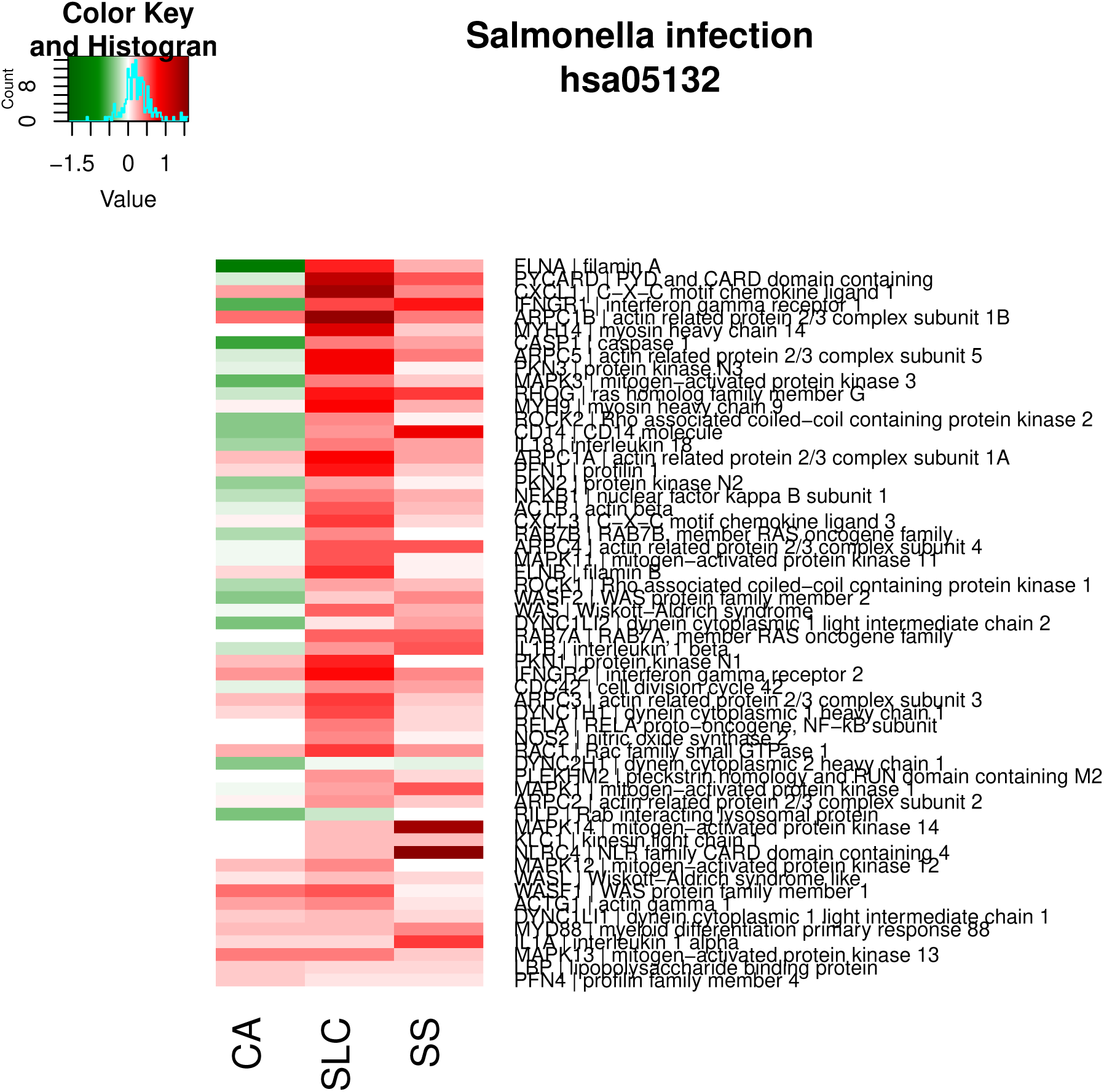

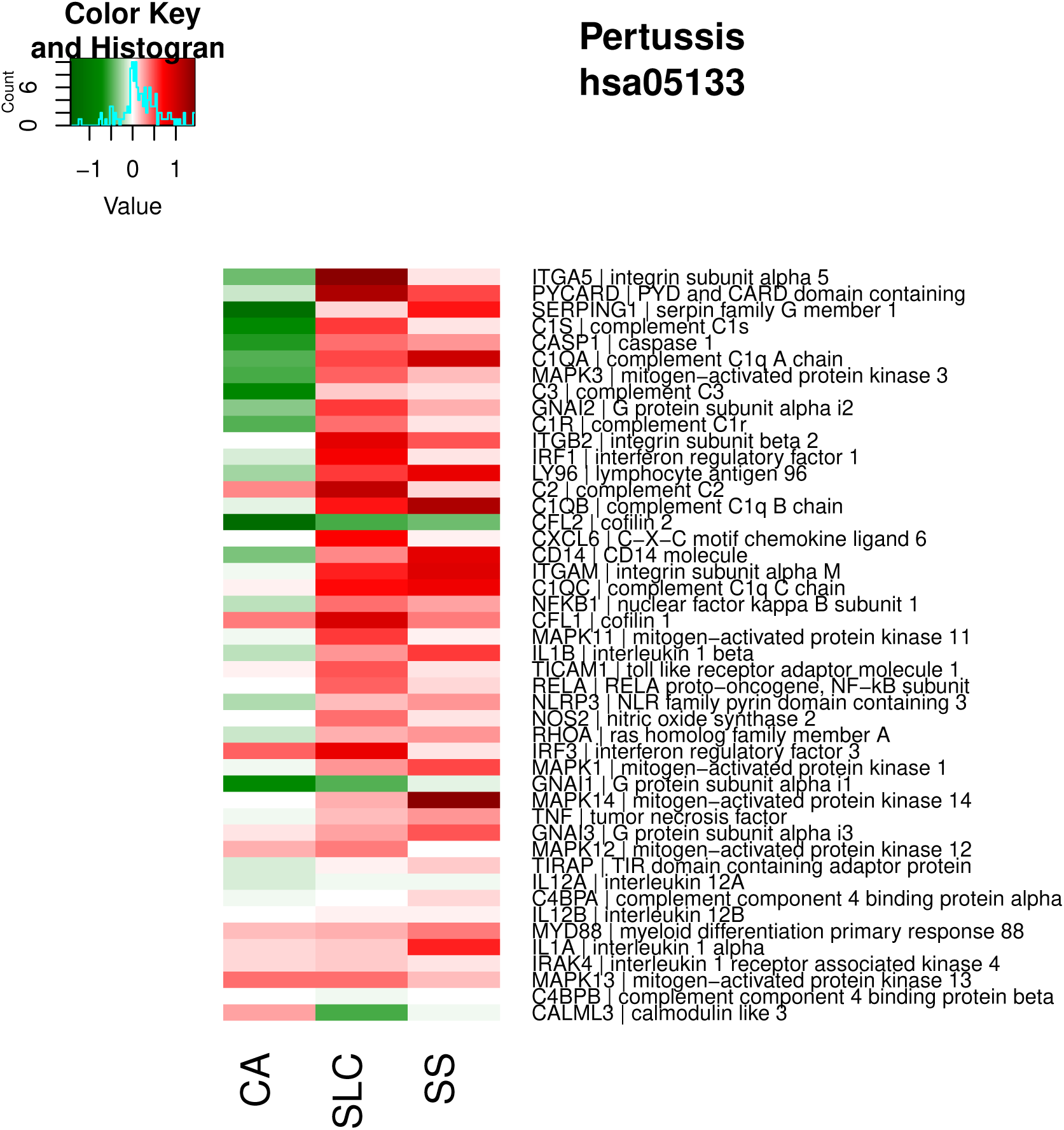

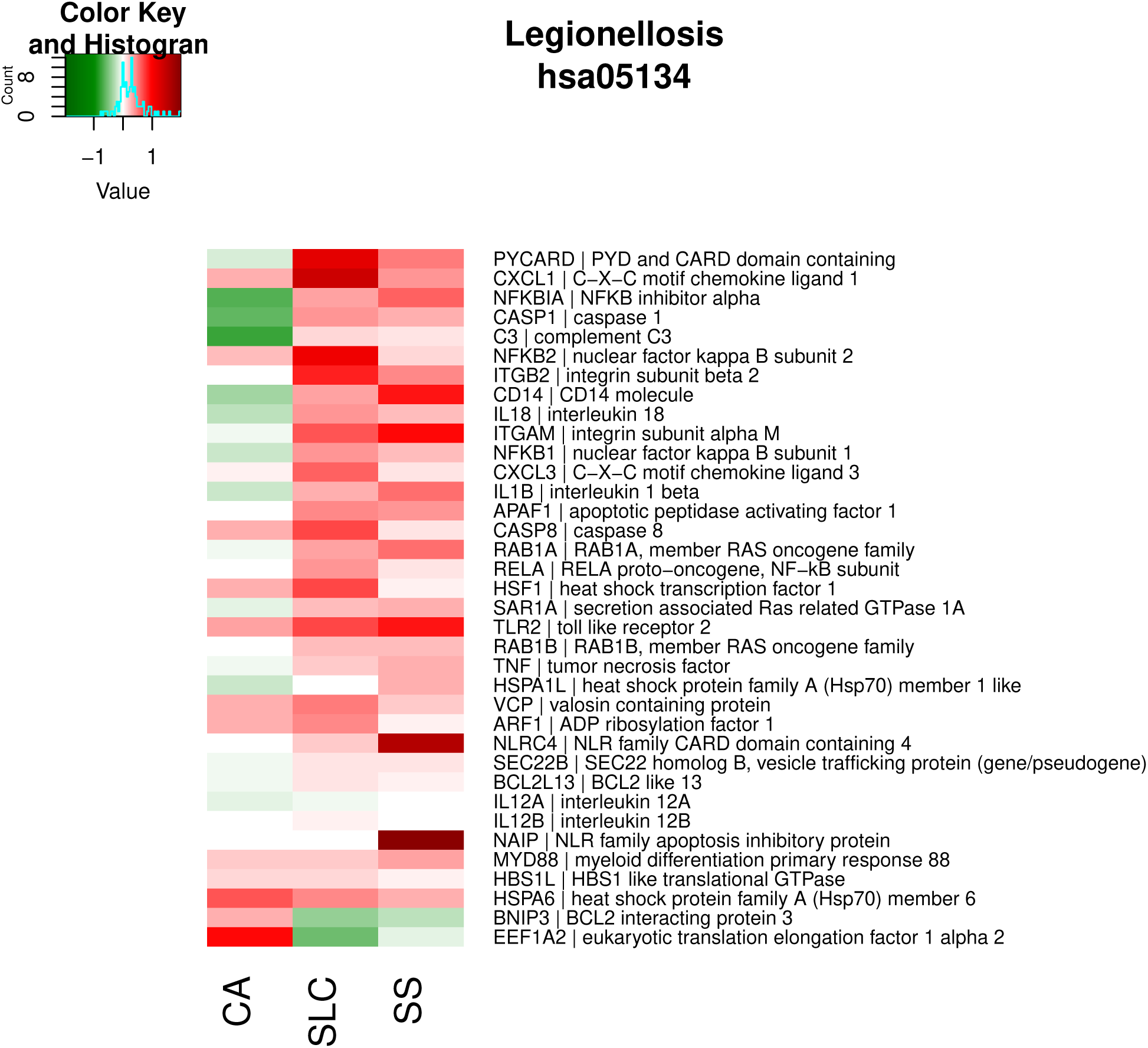

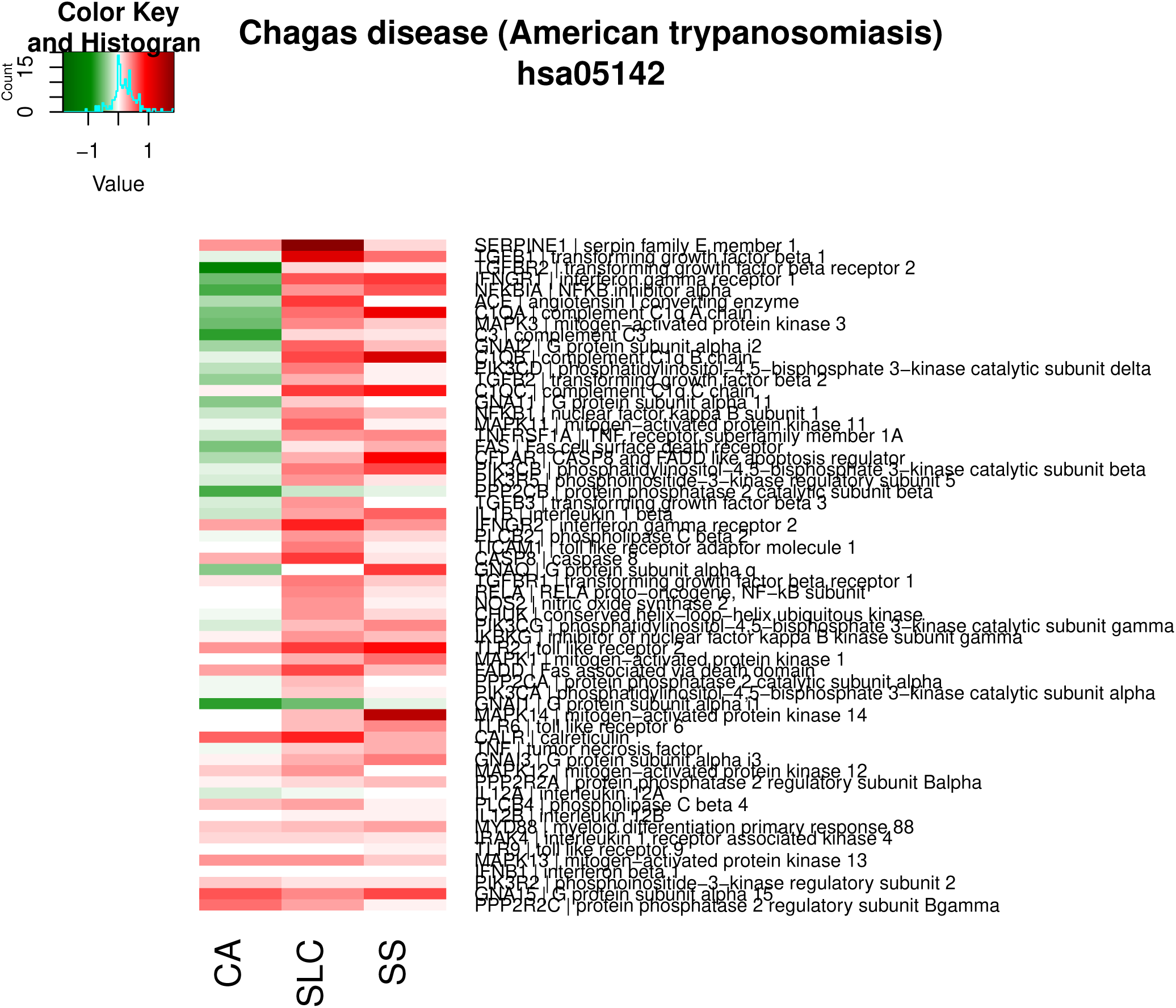

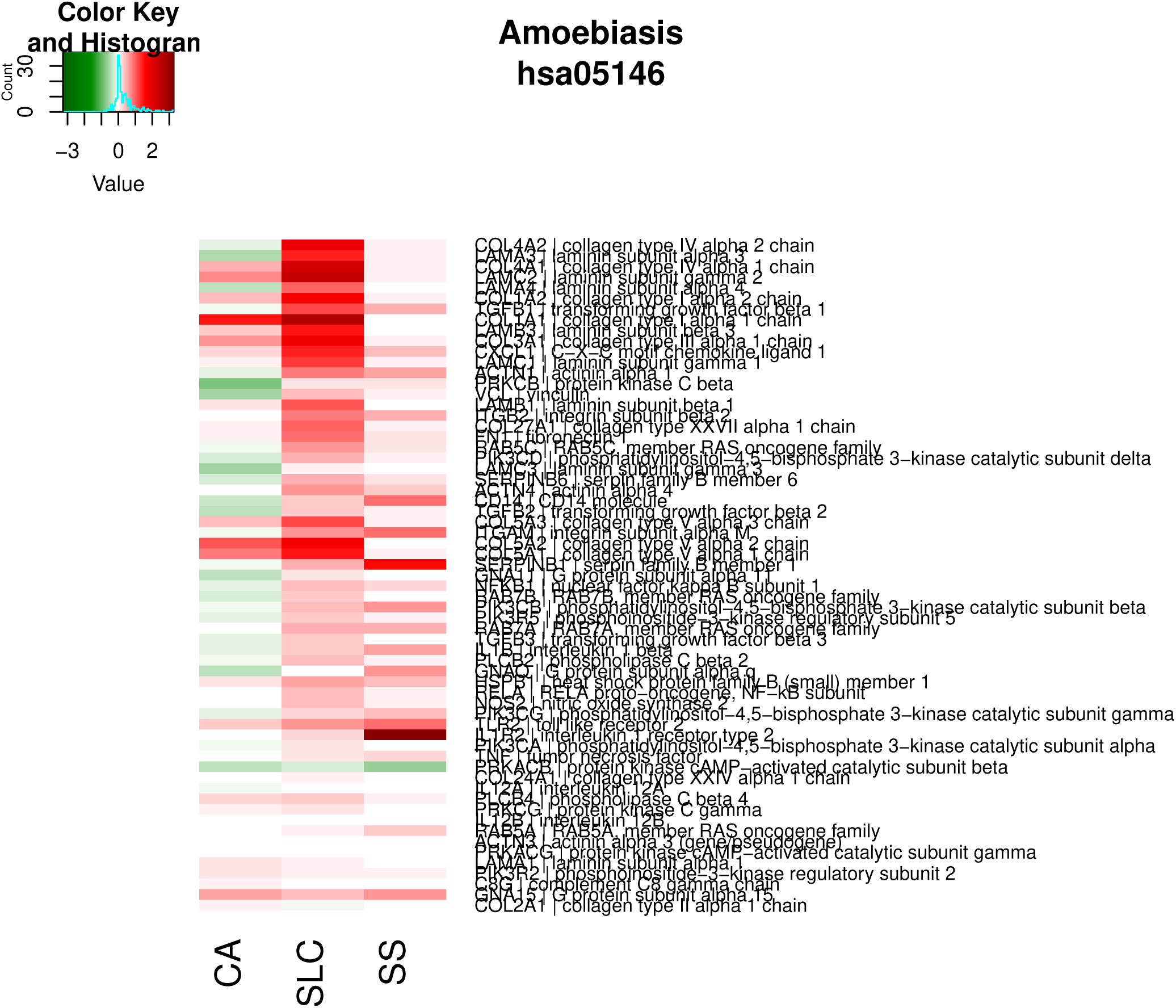

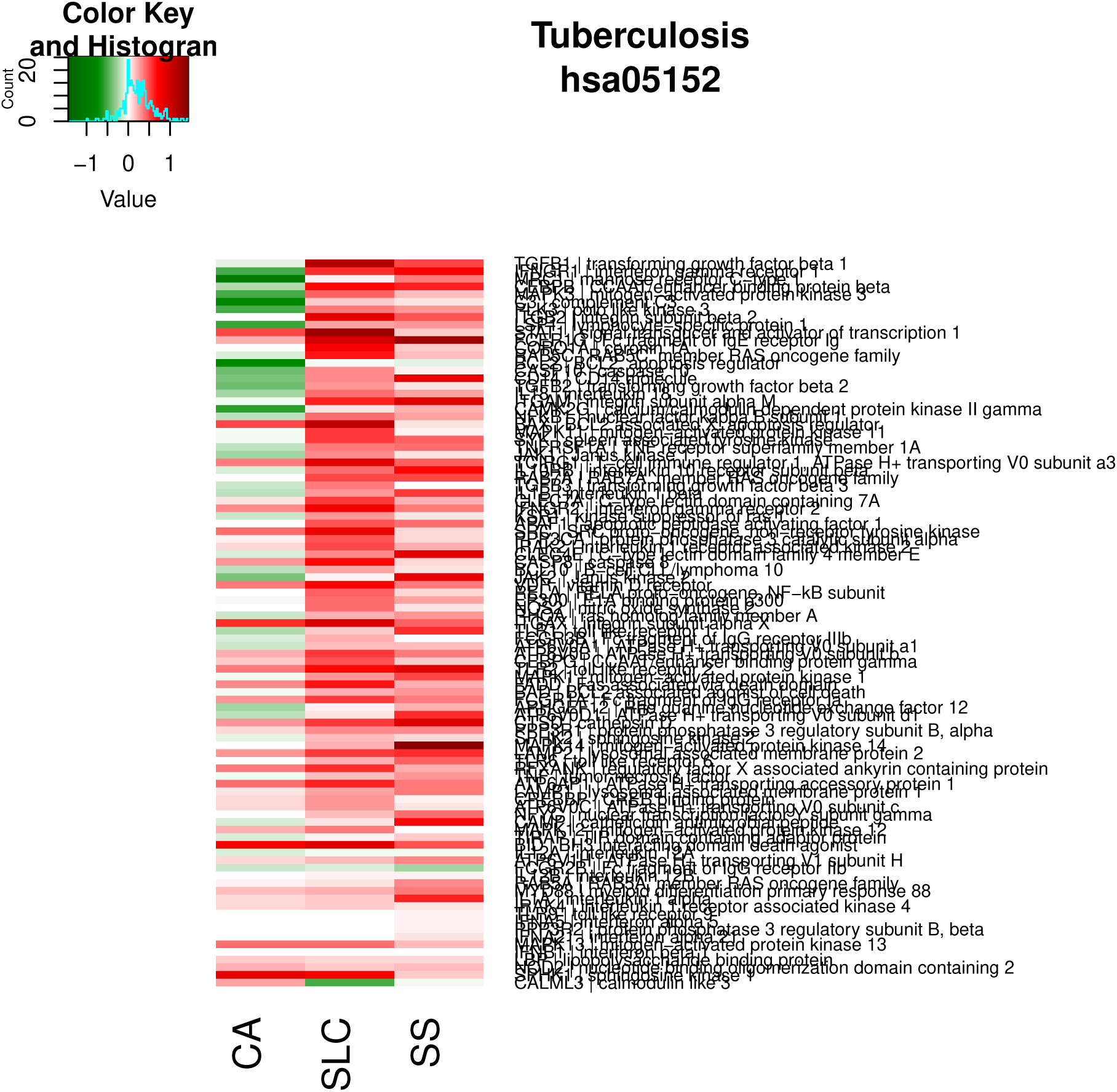

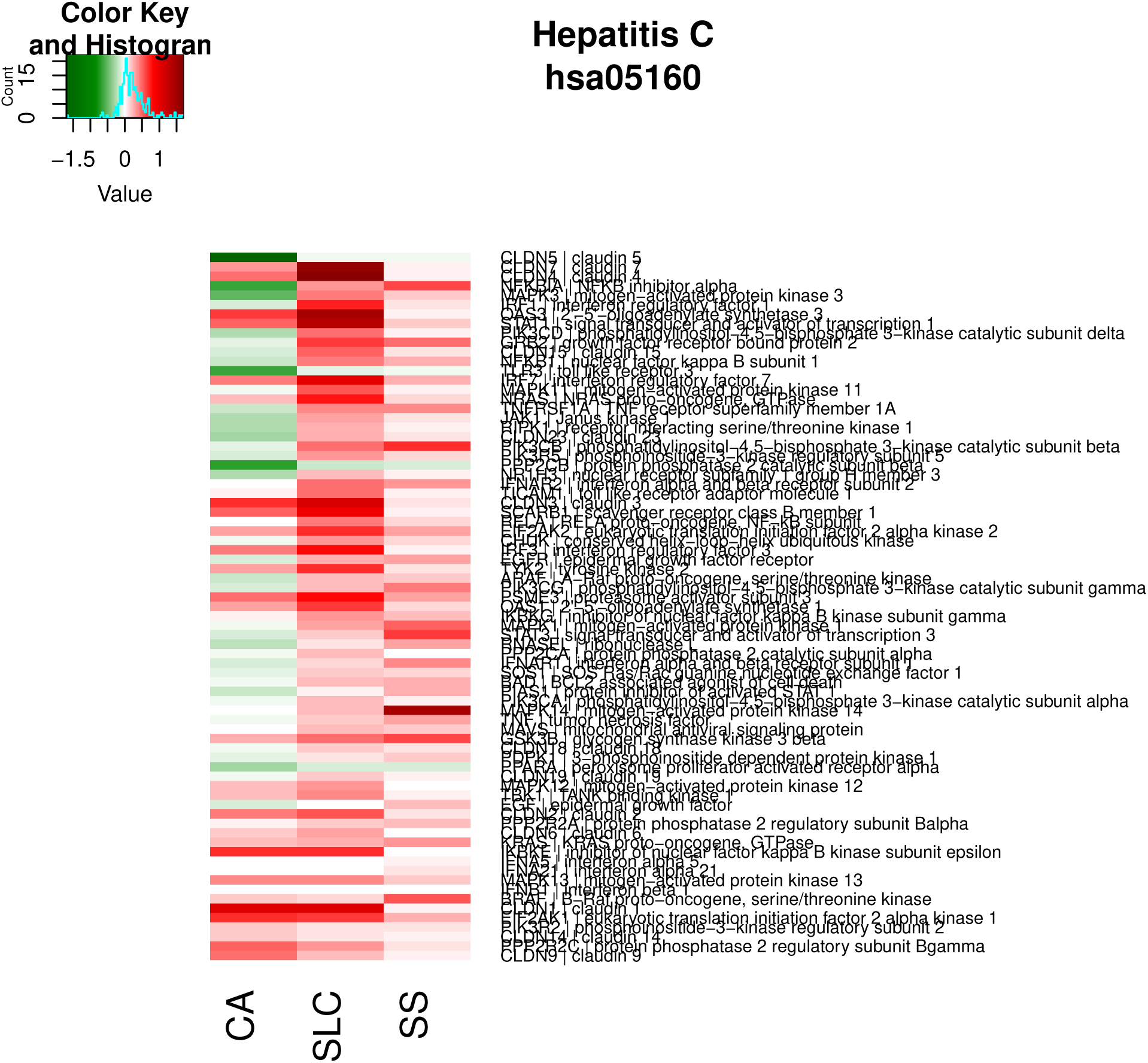

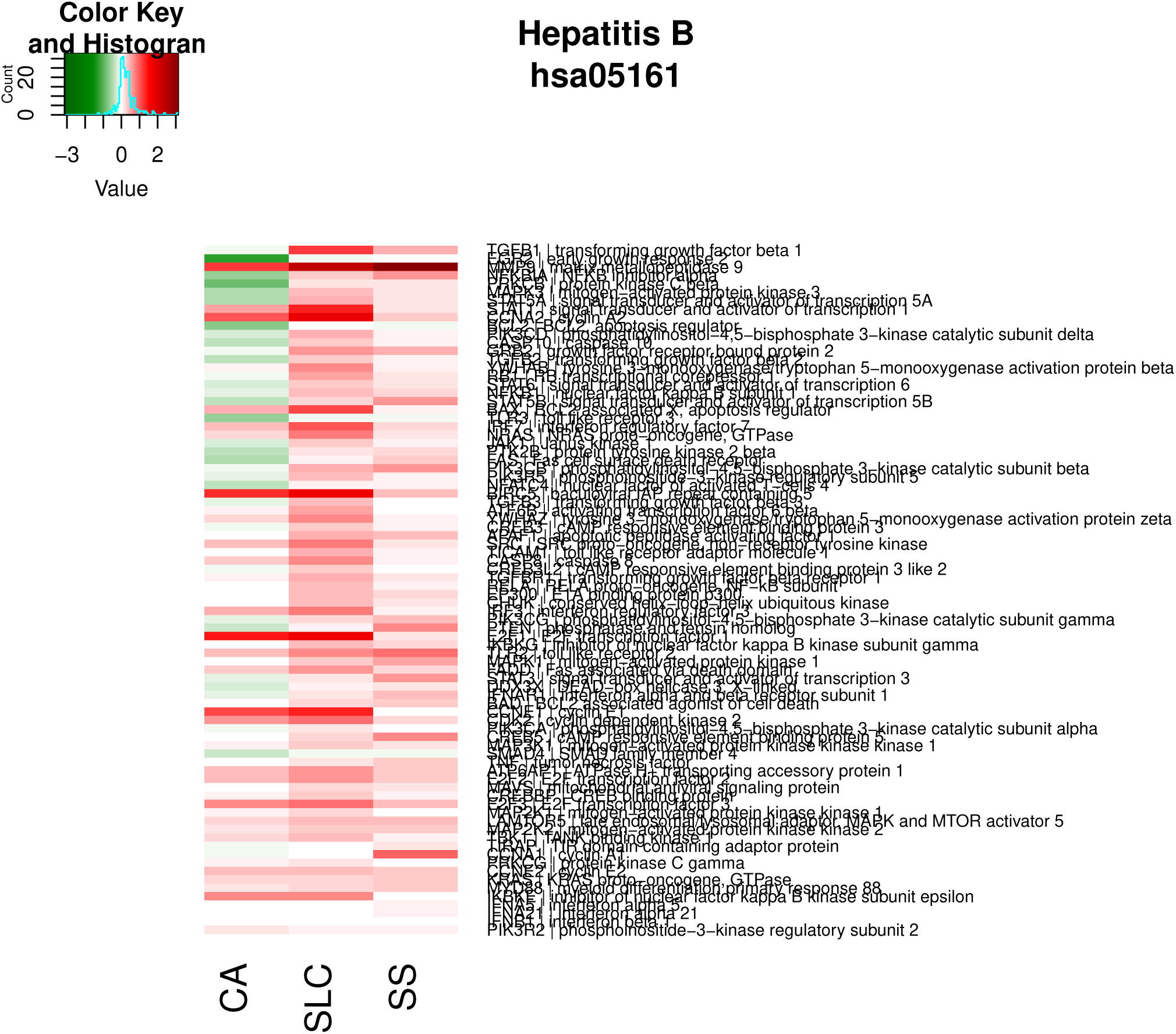

#### Other

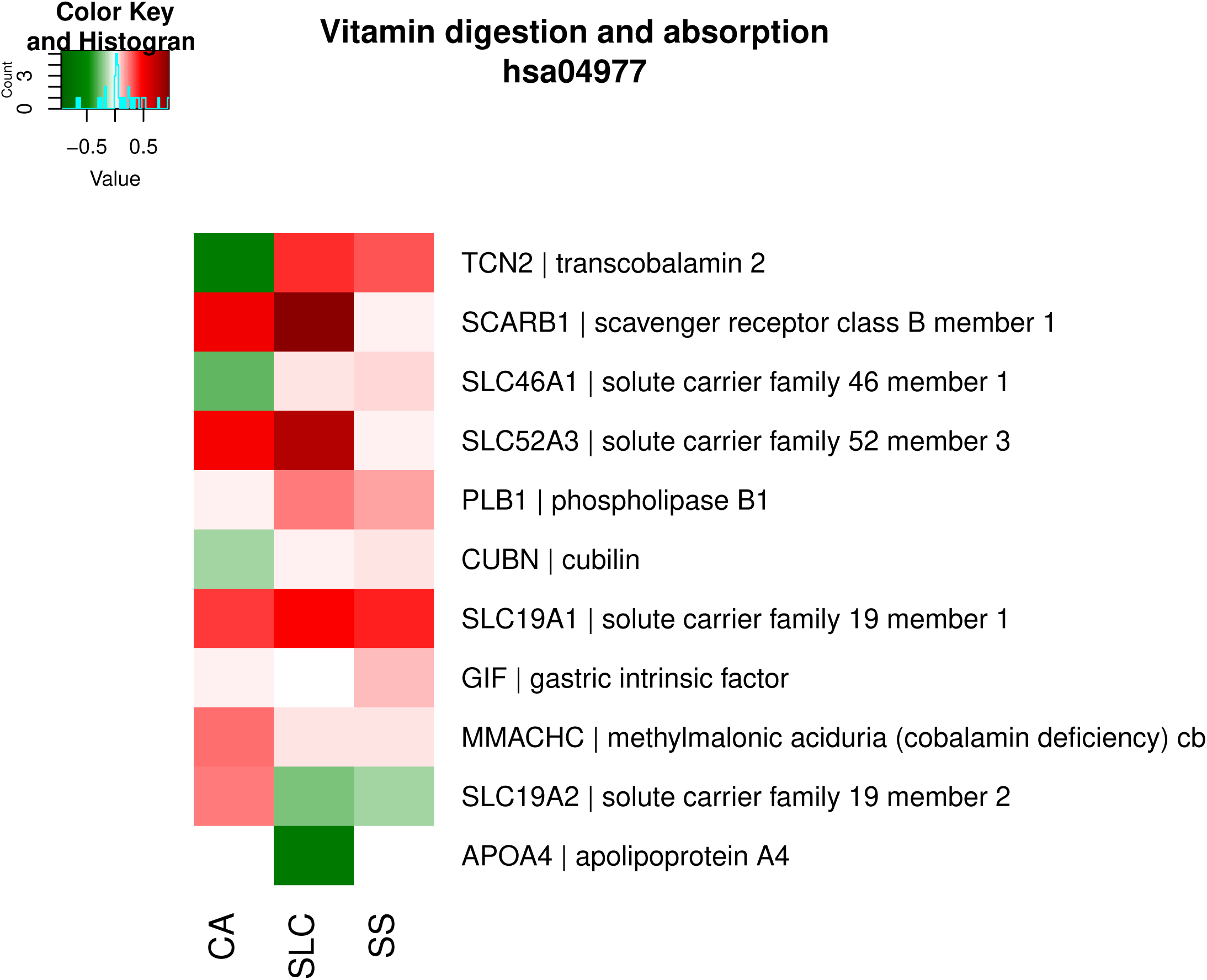

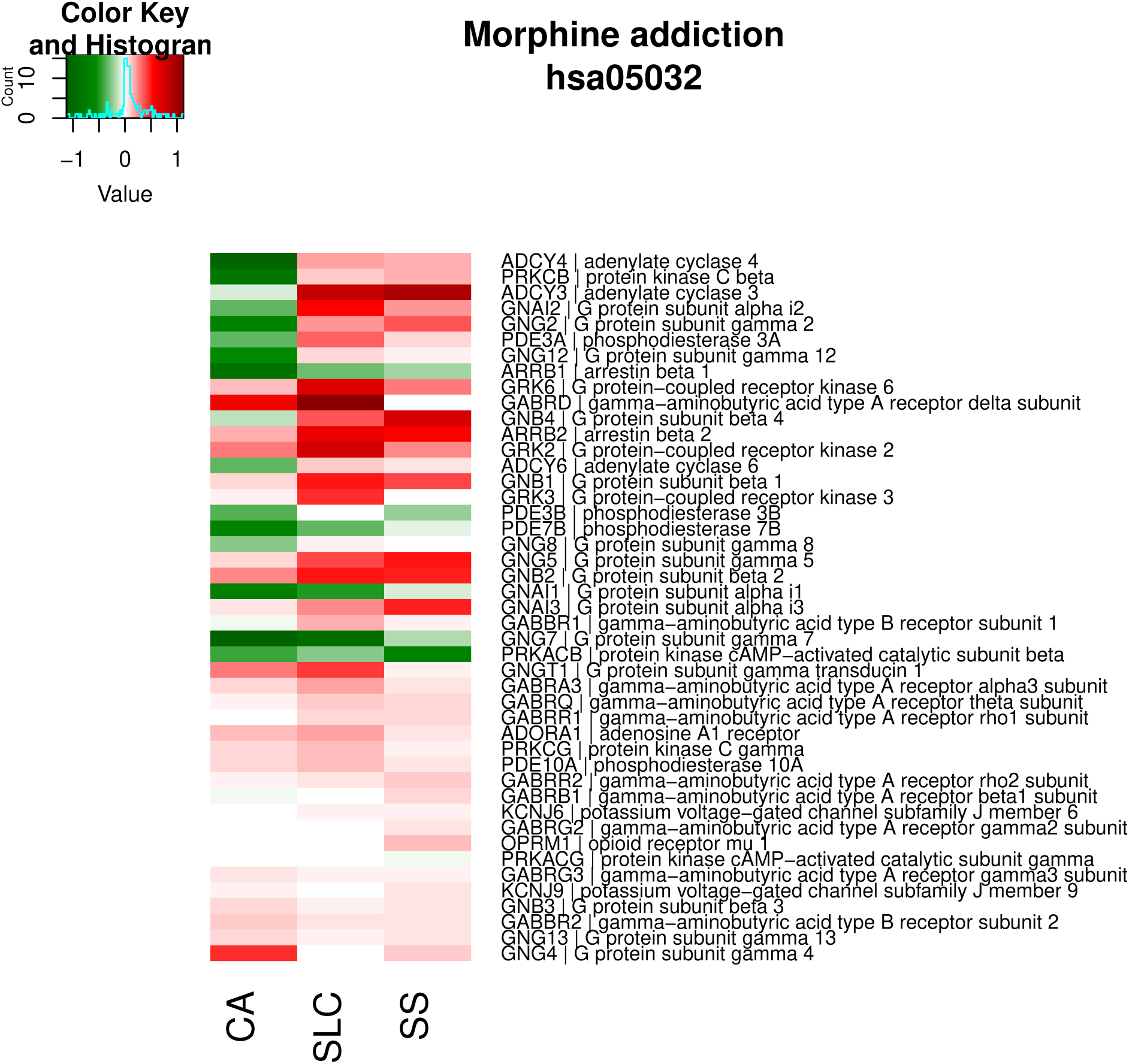

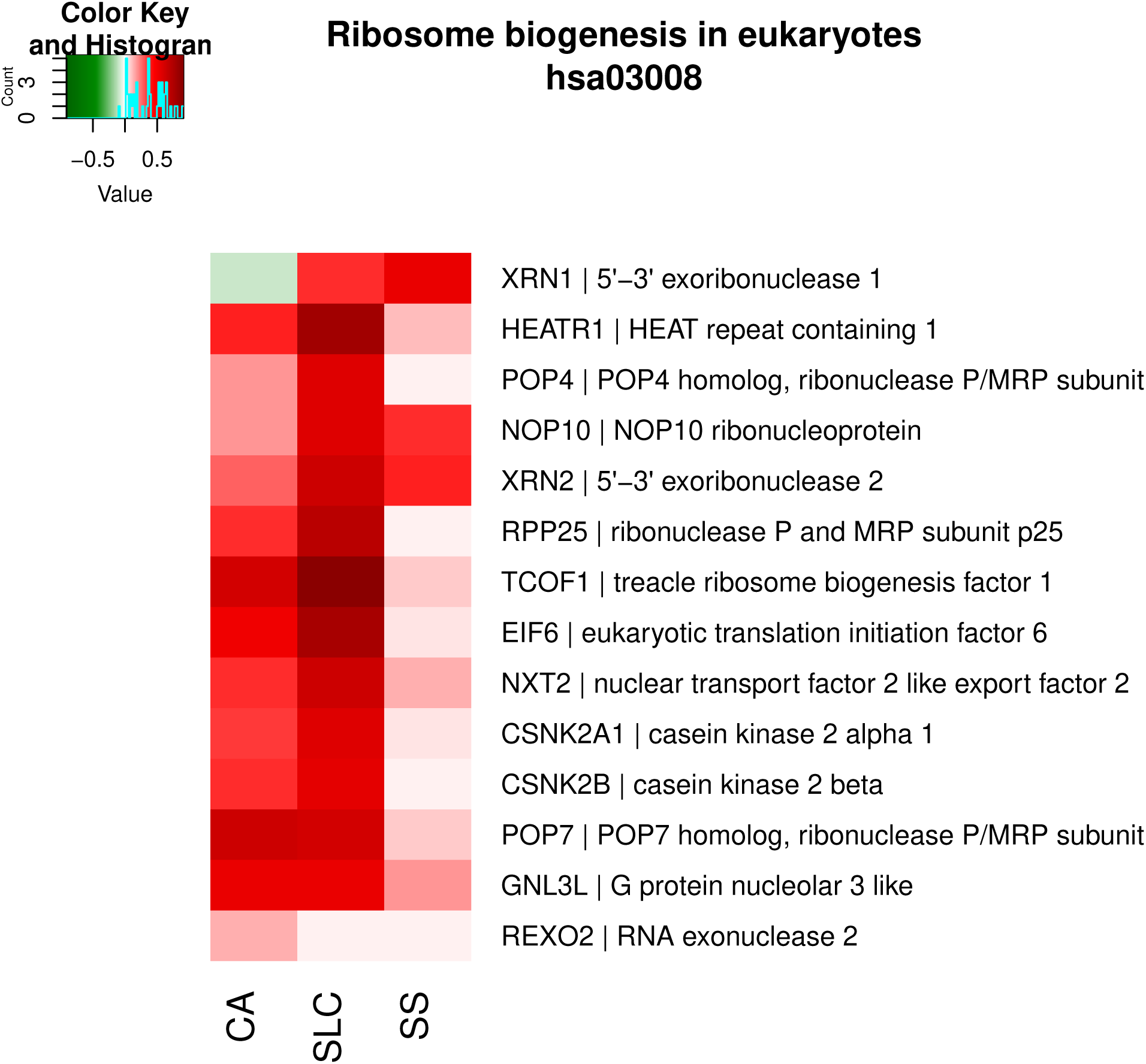

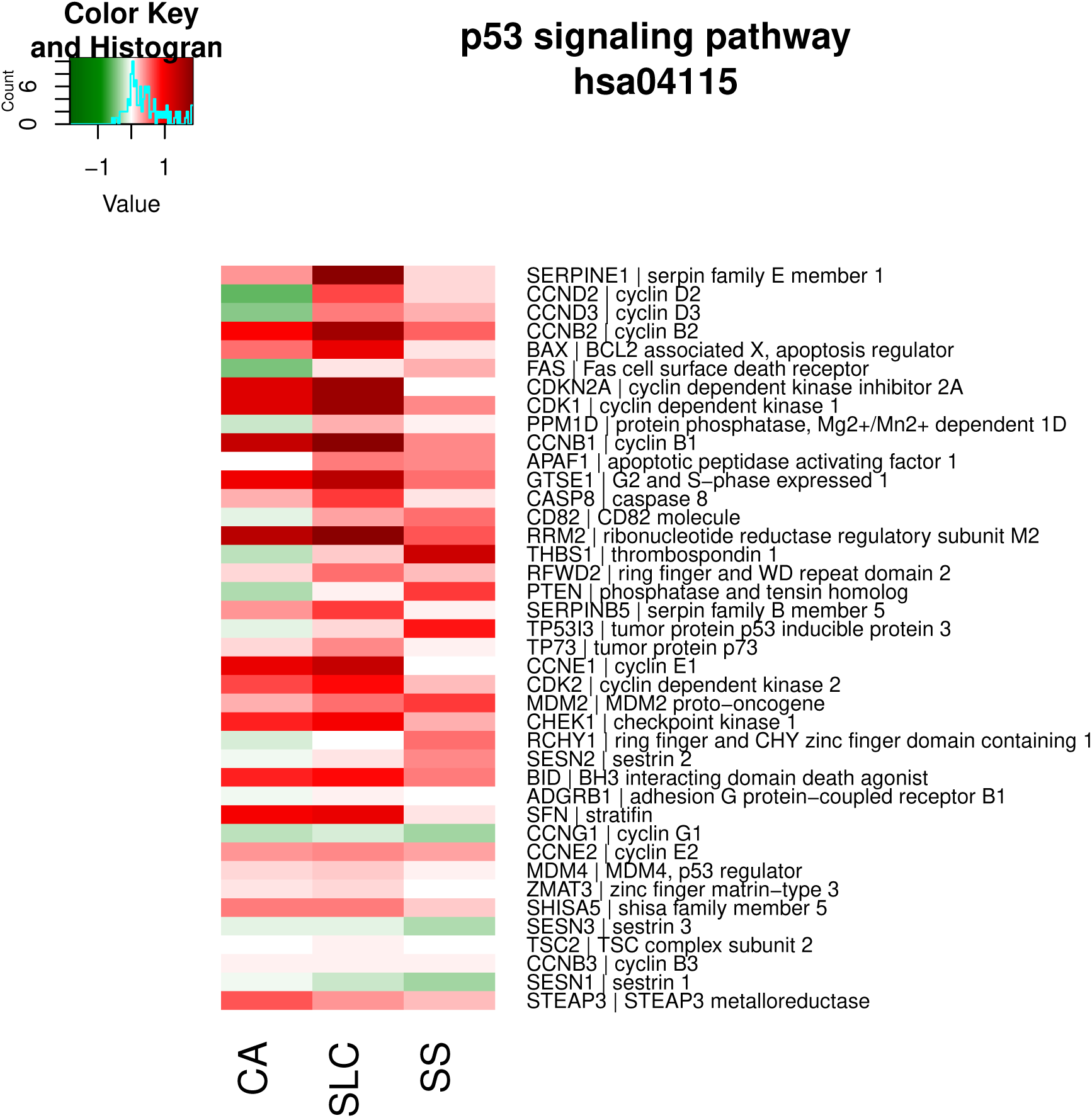

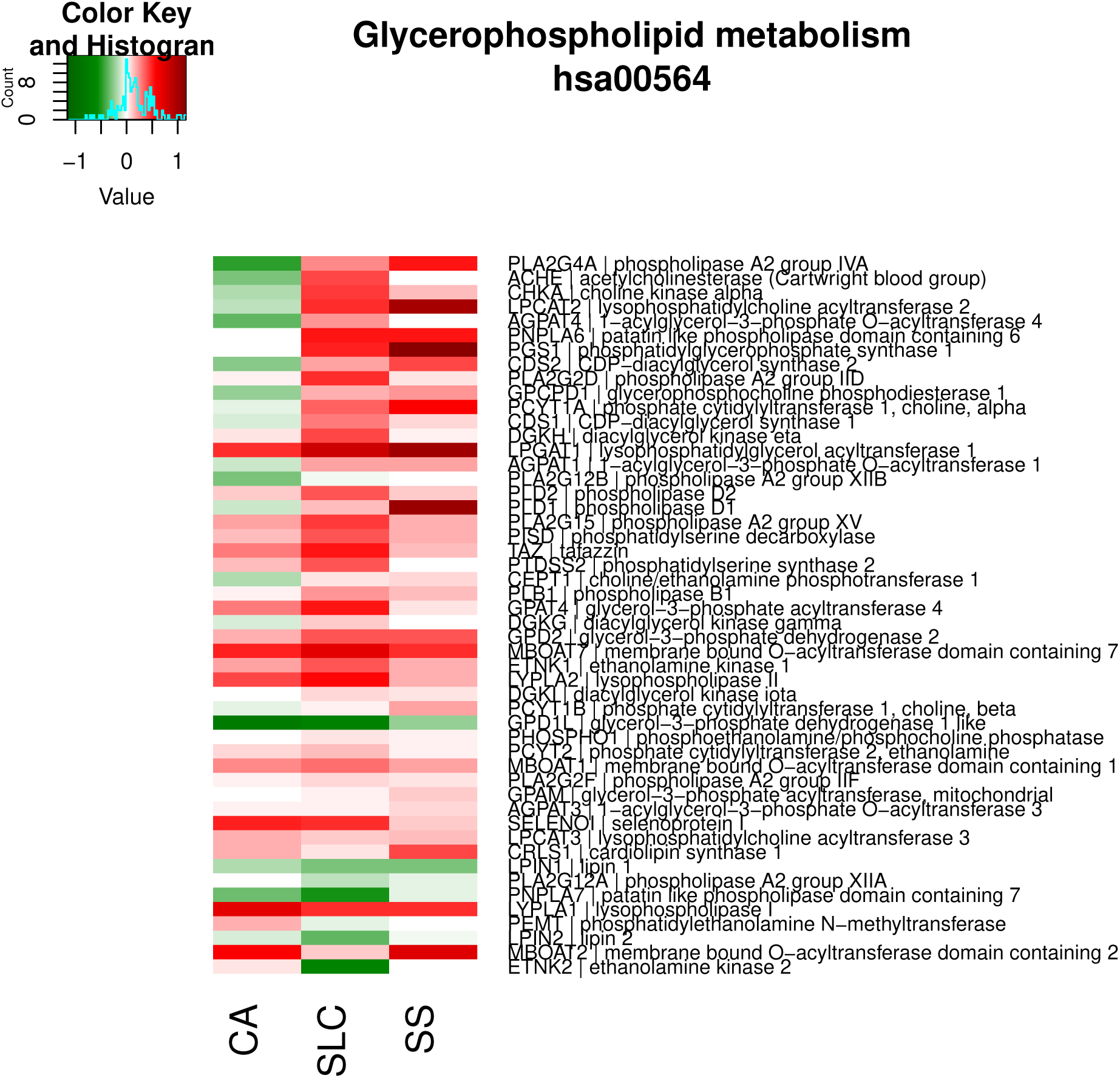

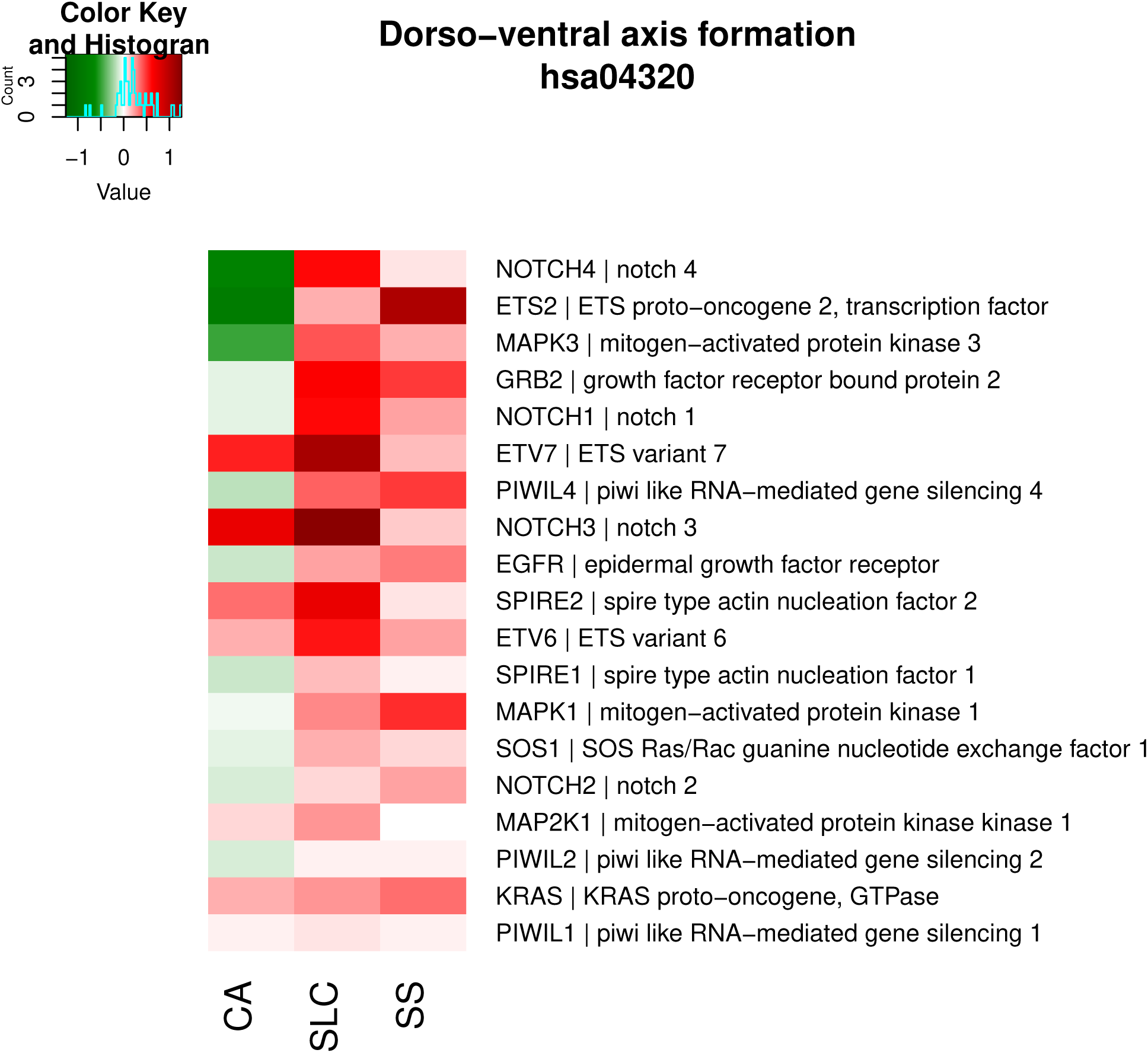

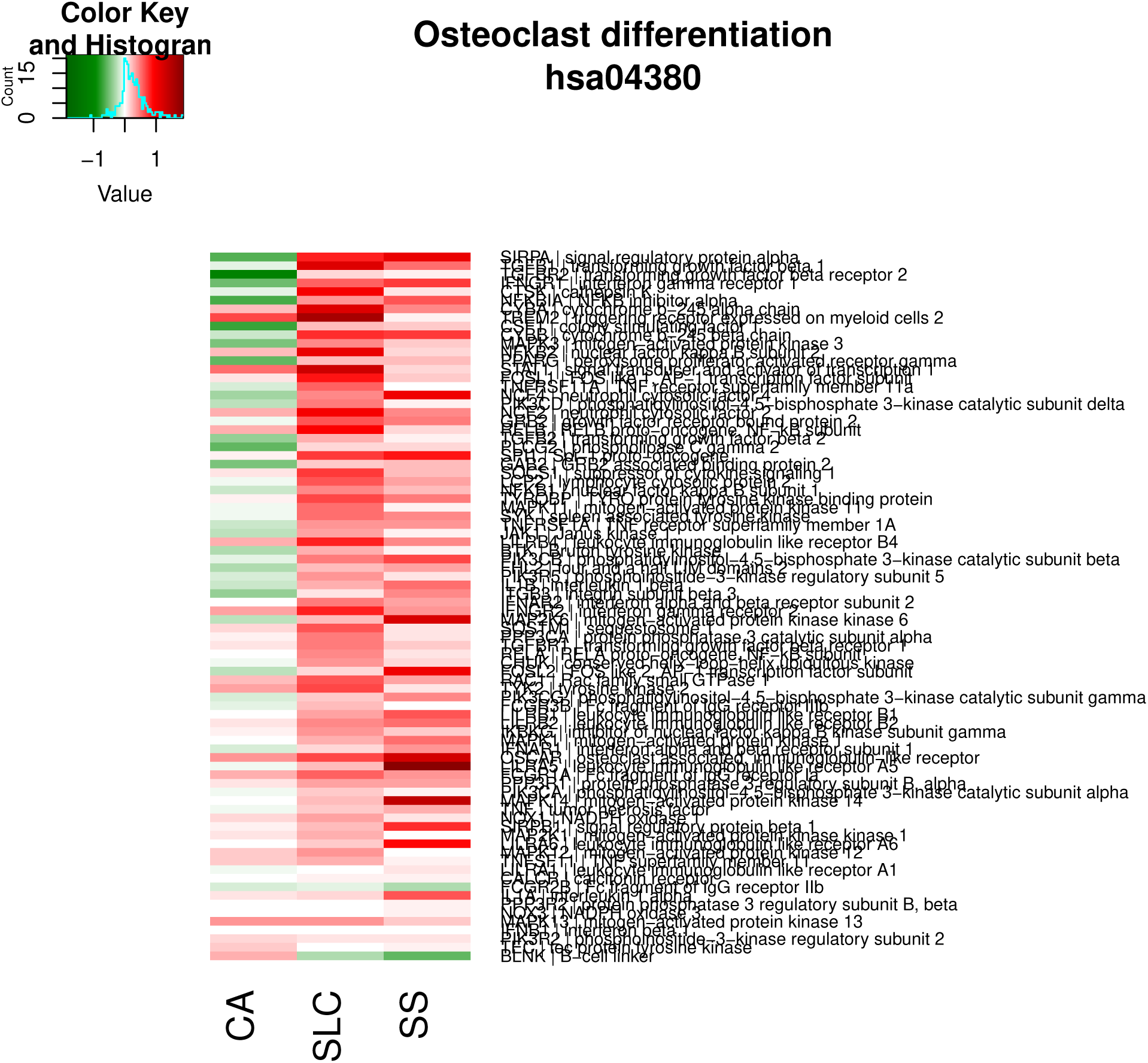

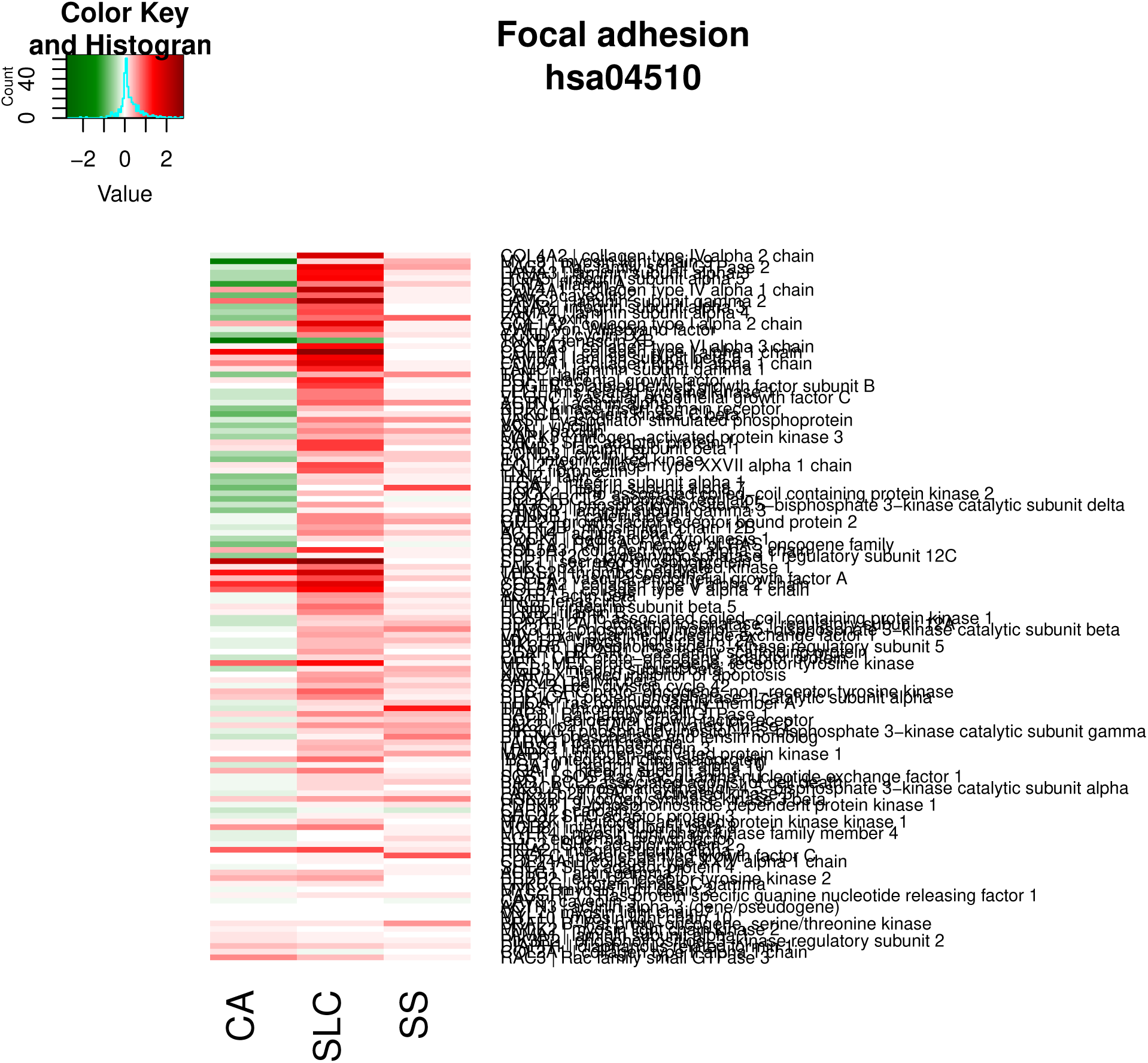

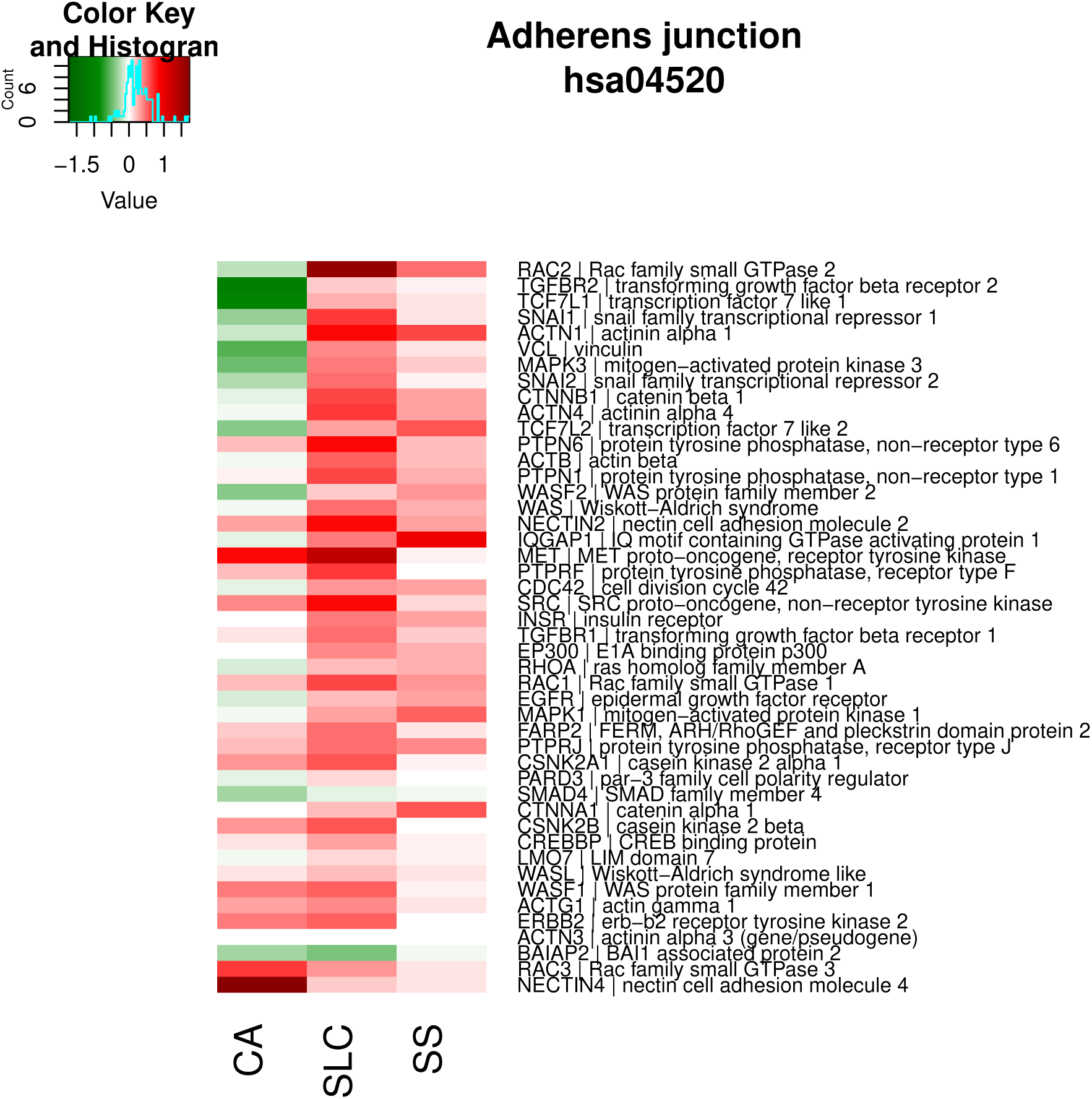

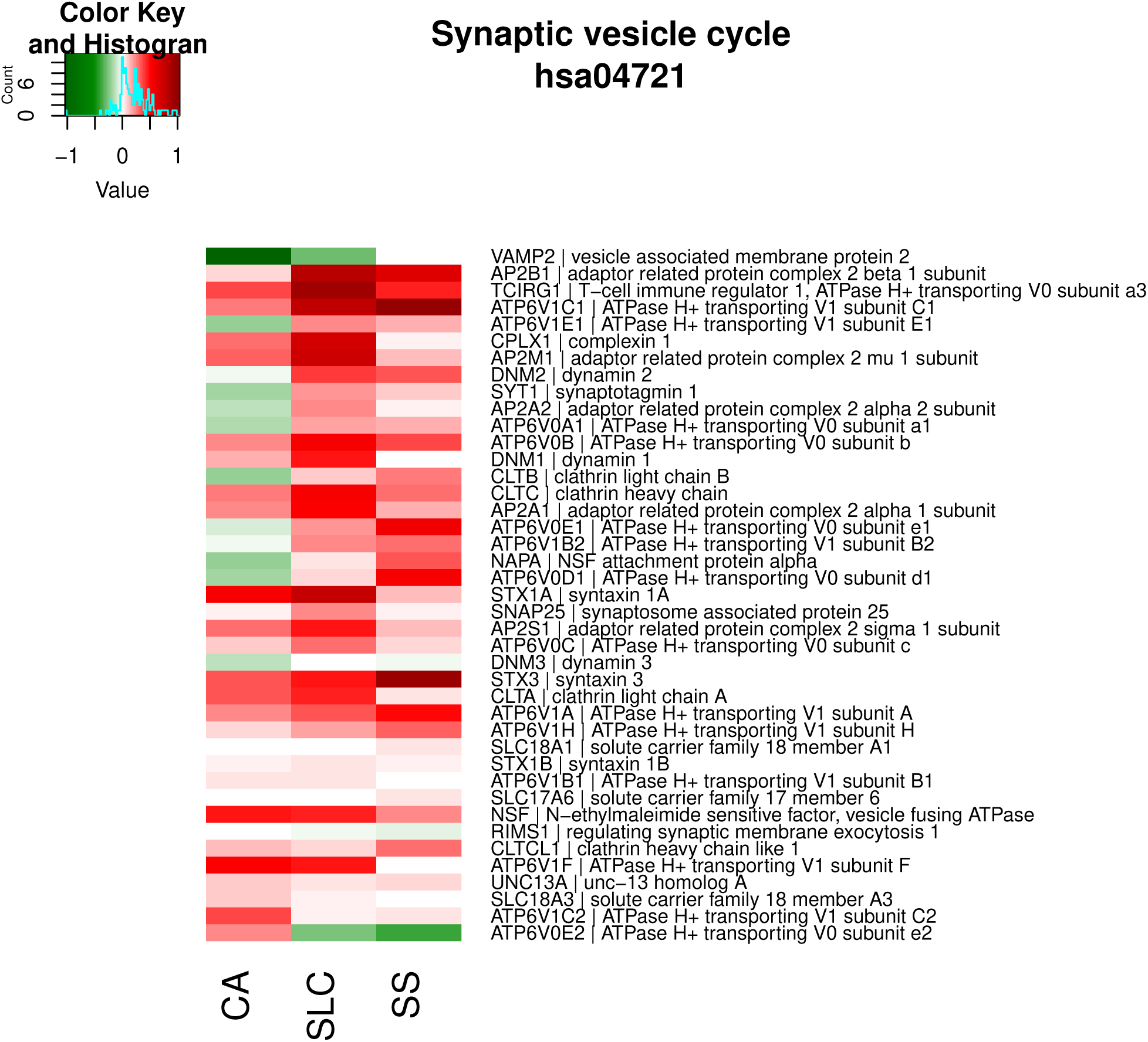

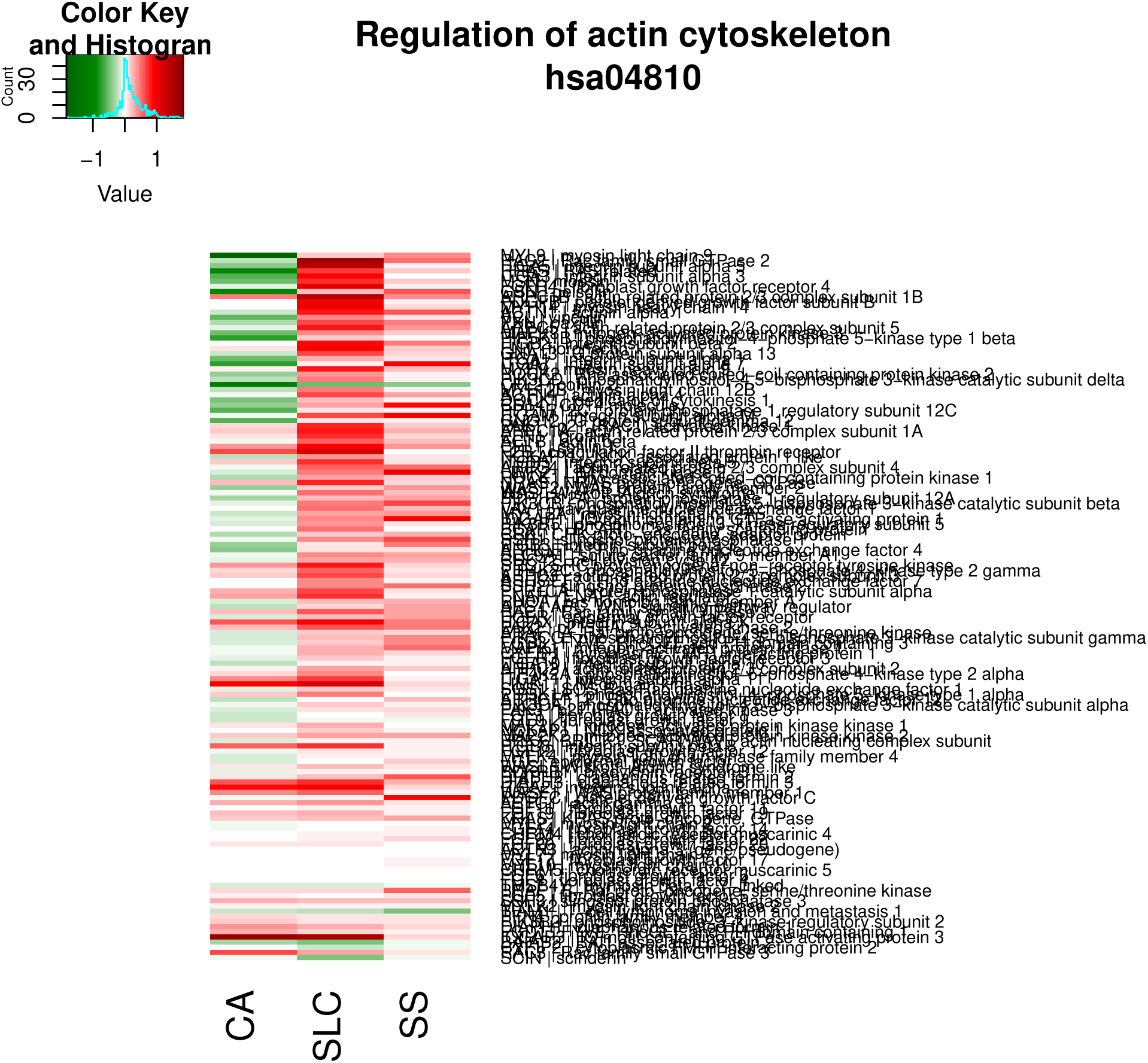

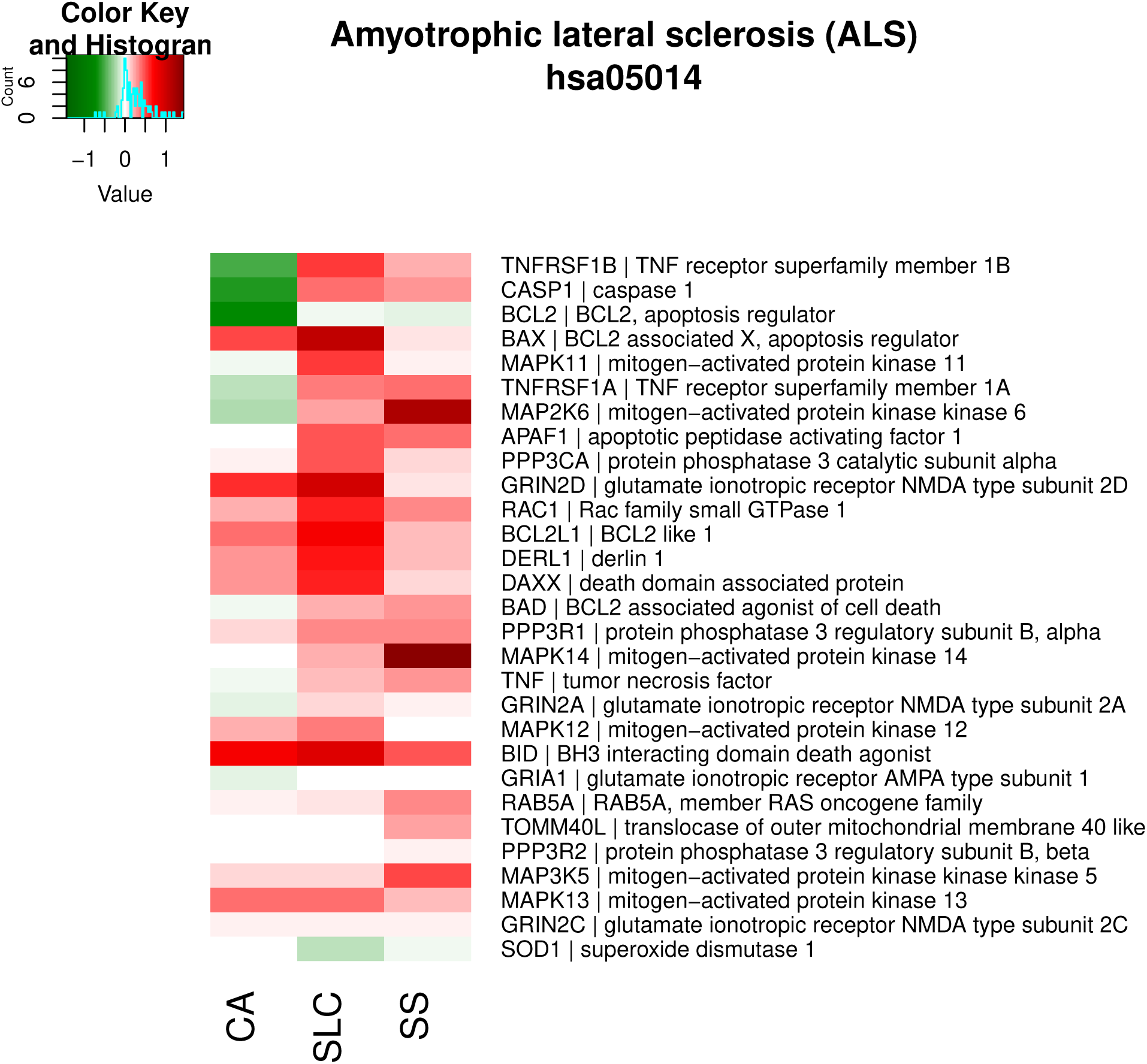

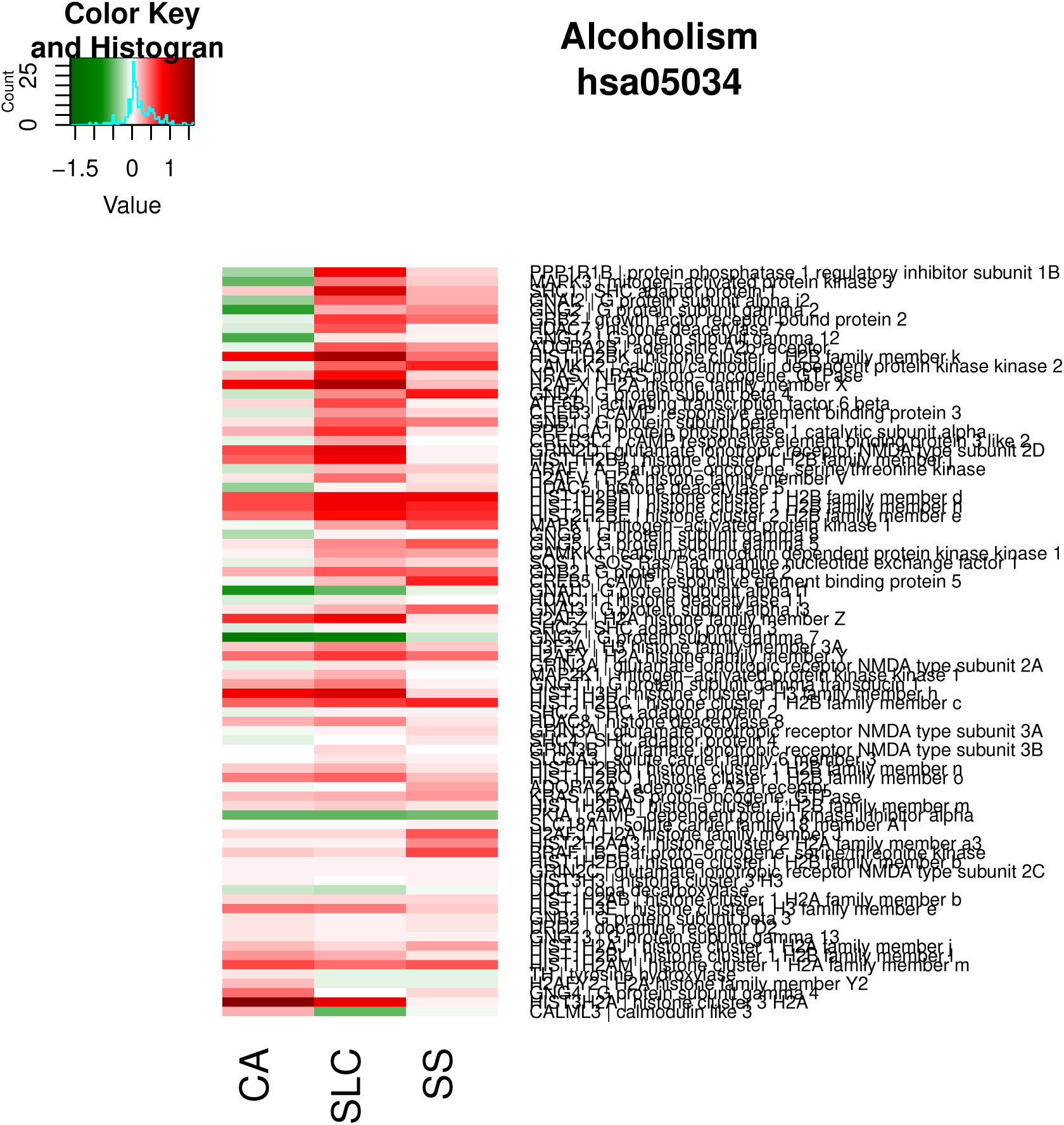

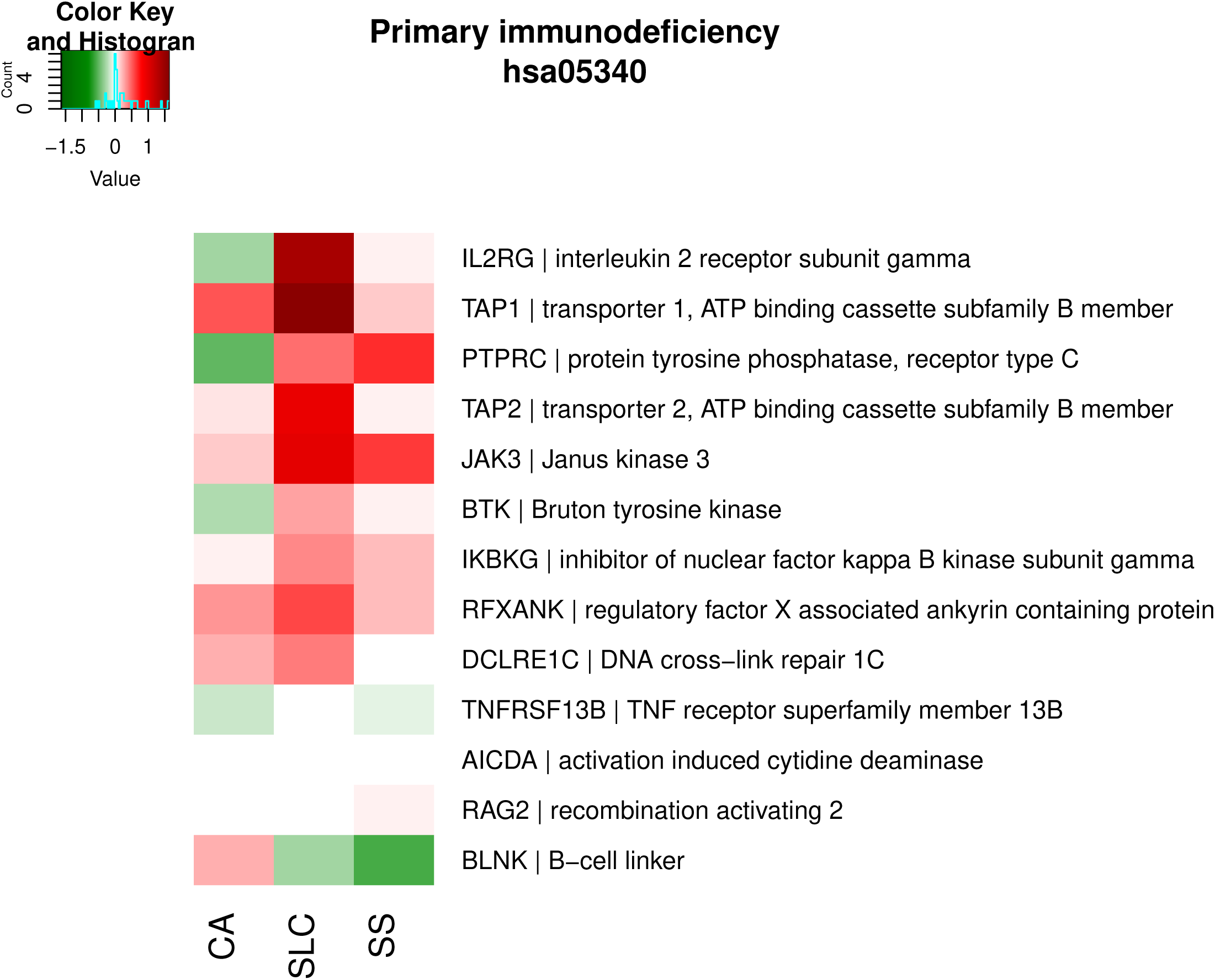

#### Signal transduction

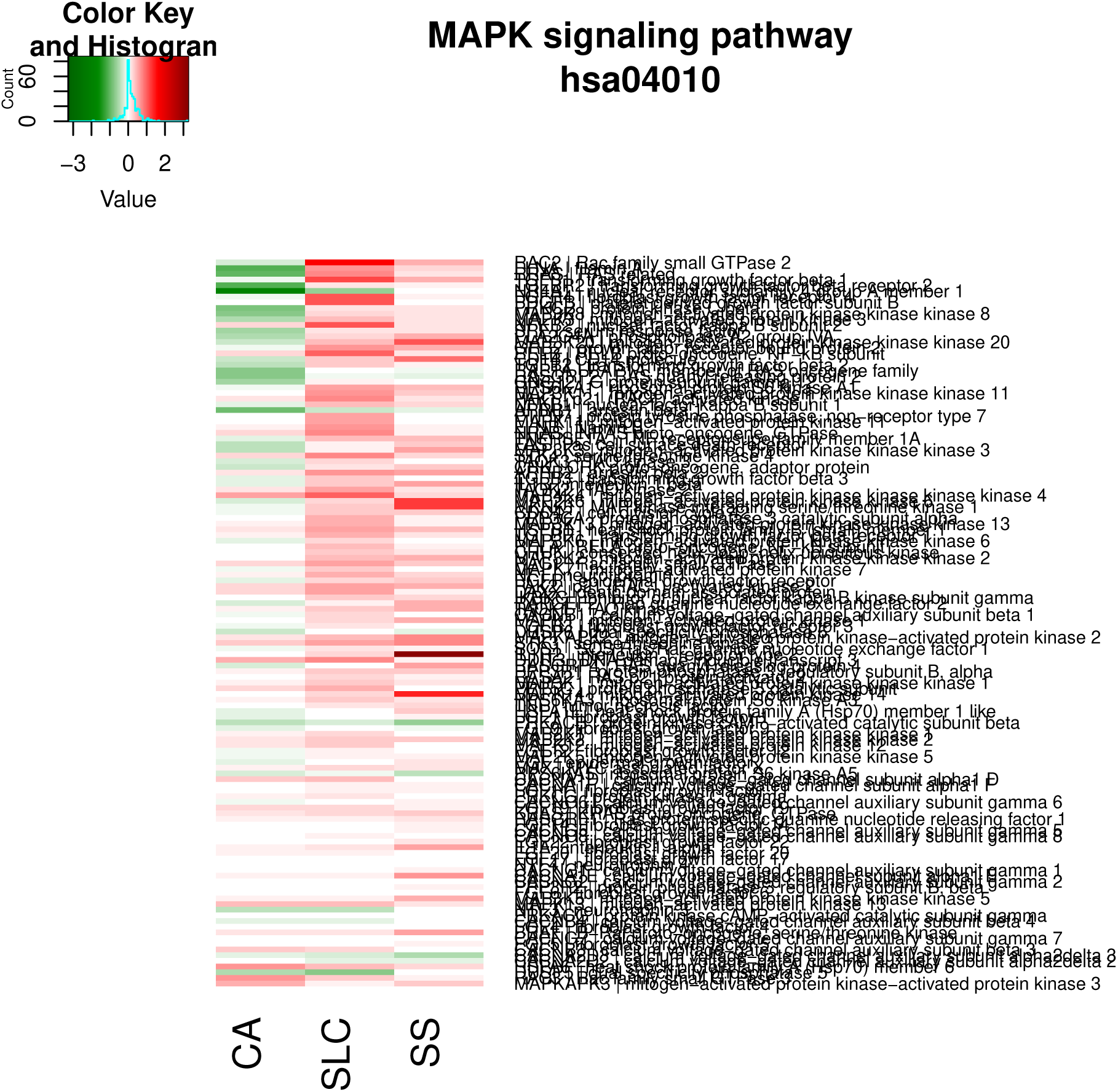

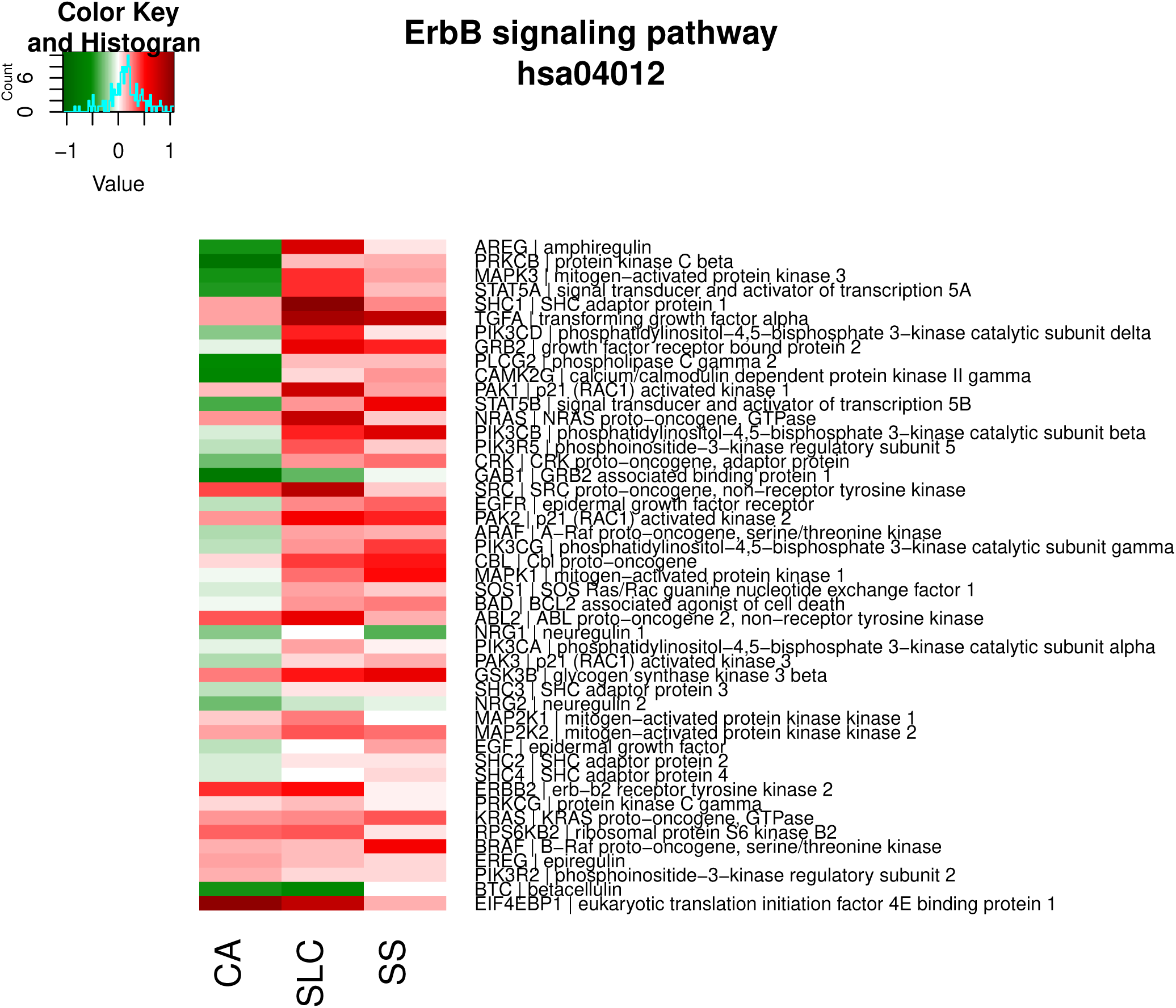

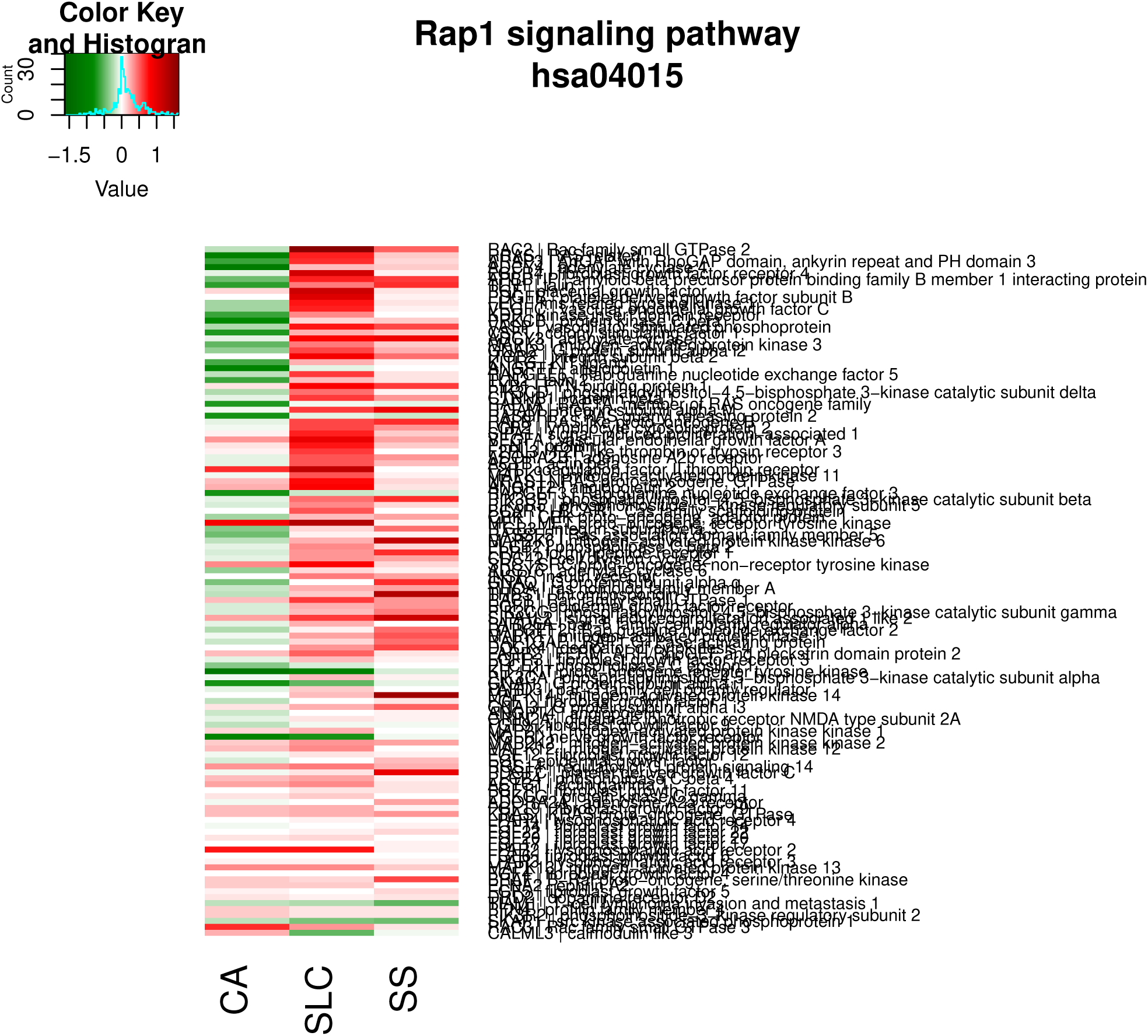

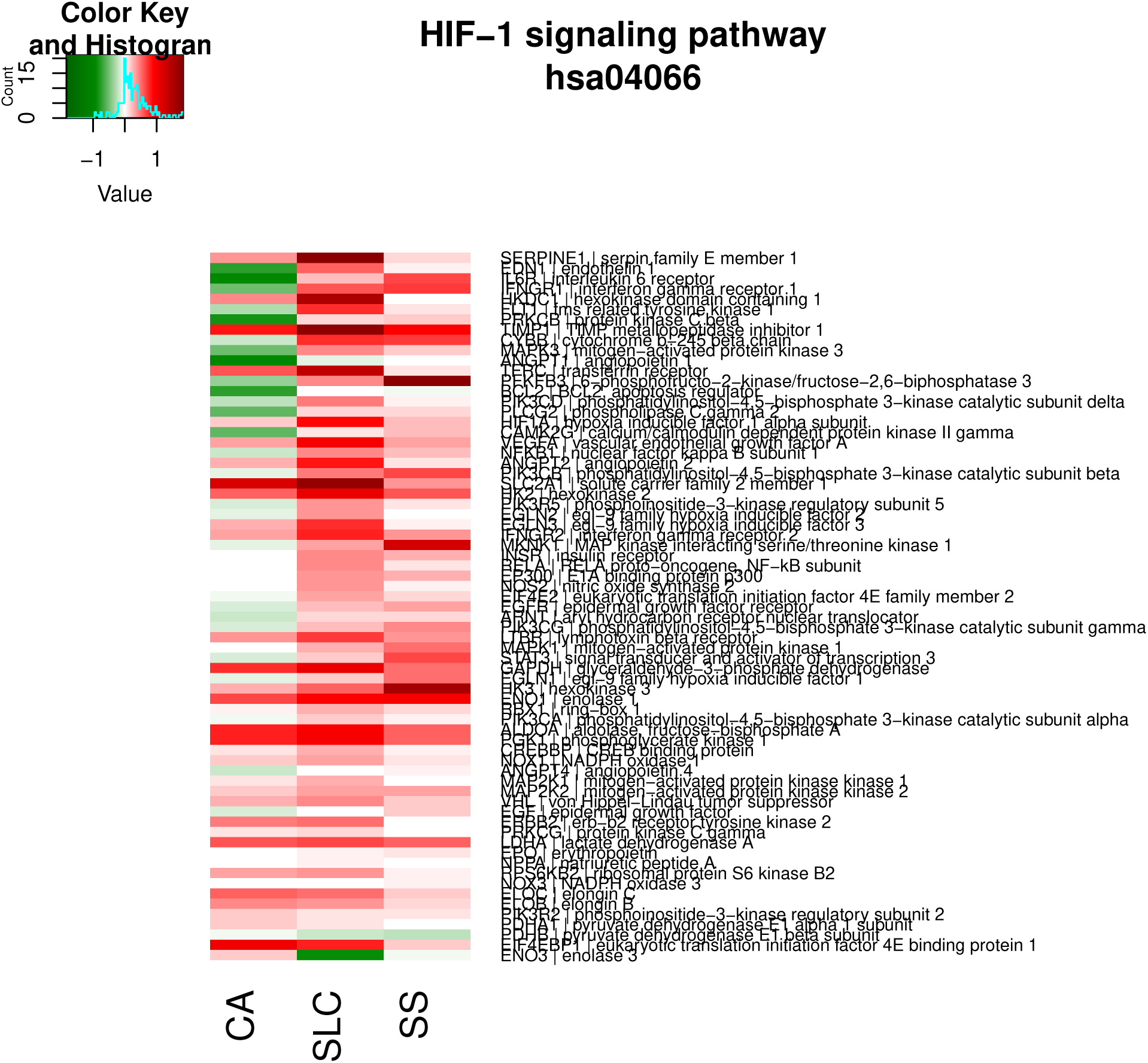

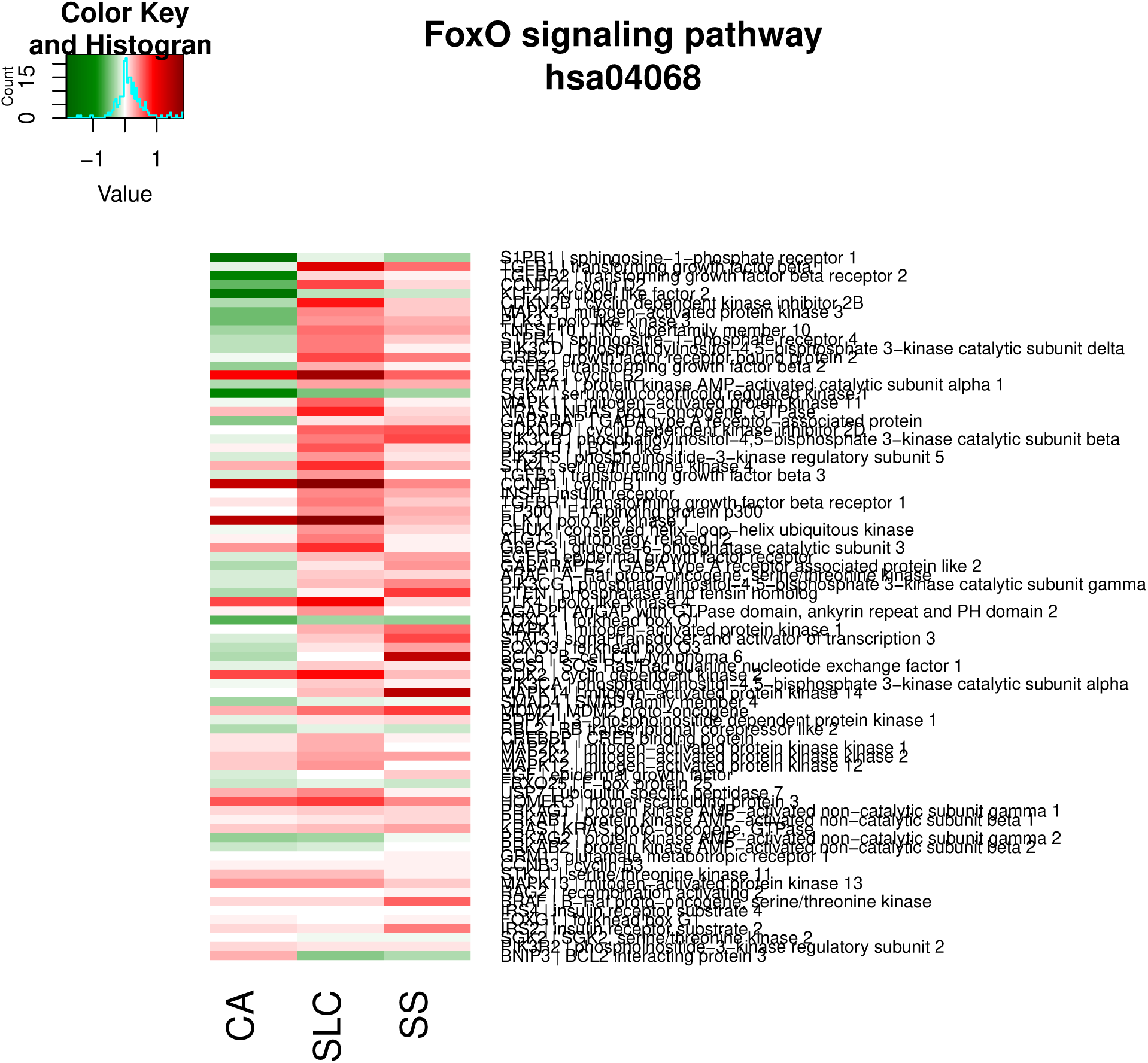

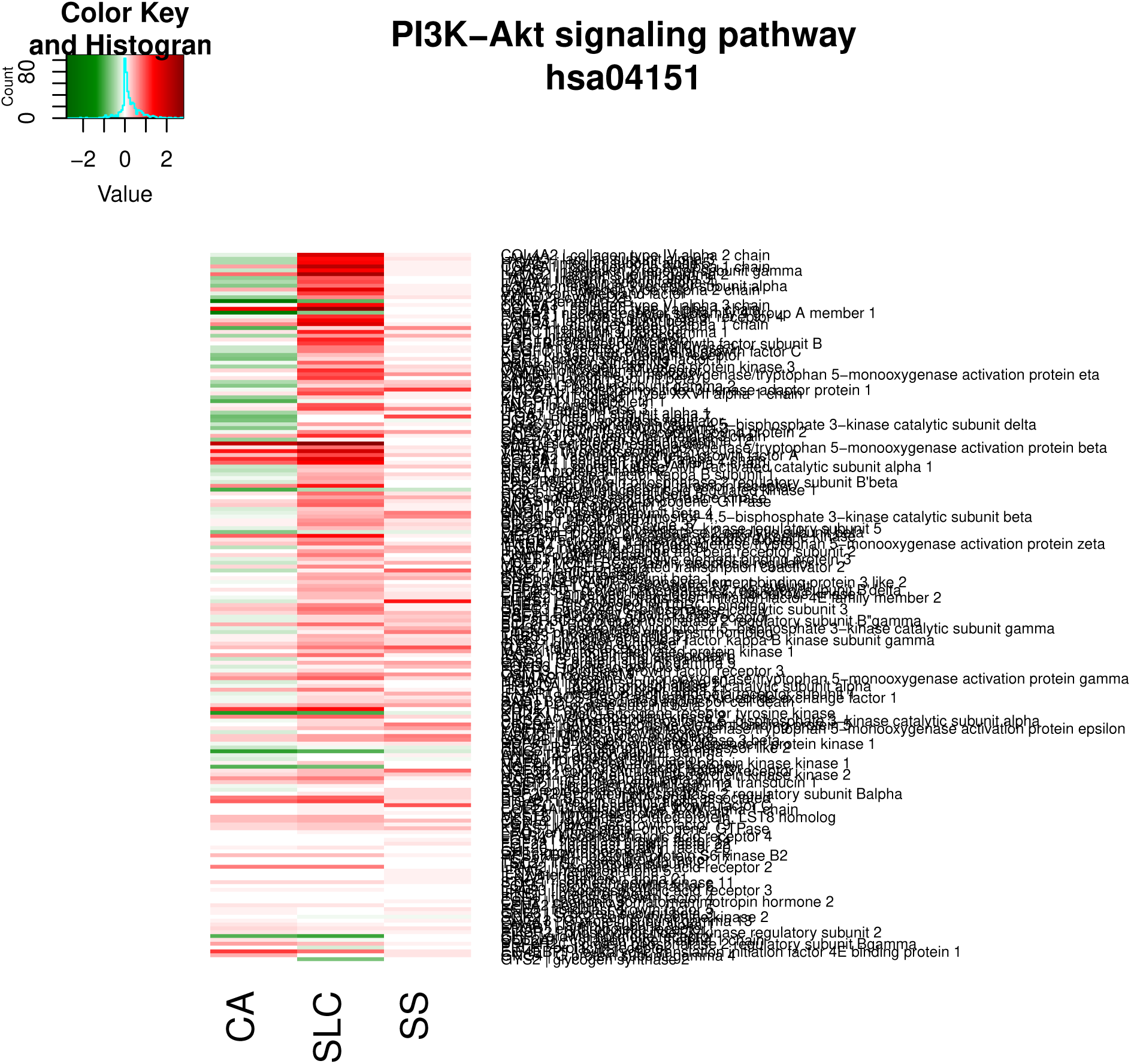

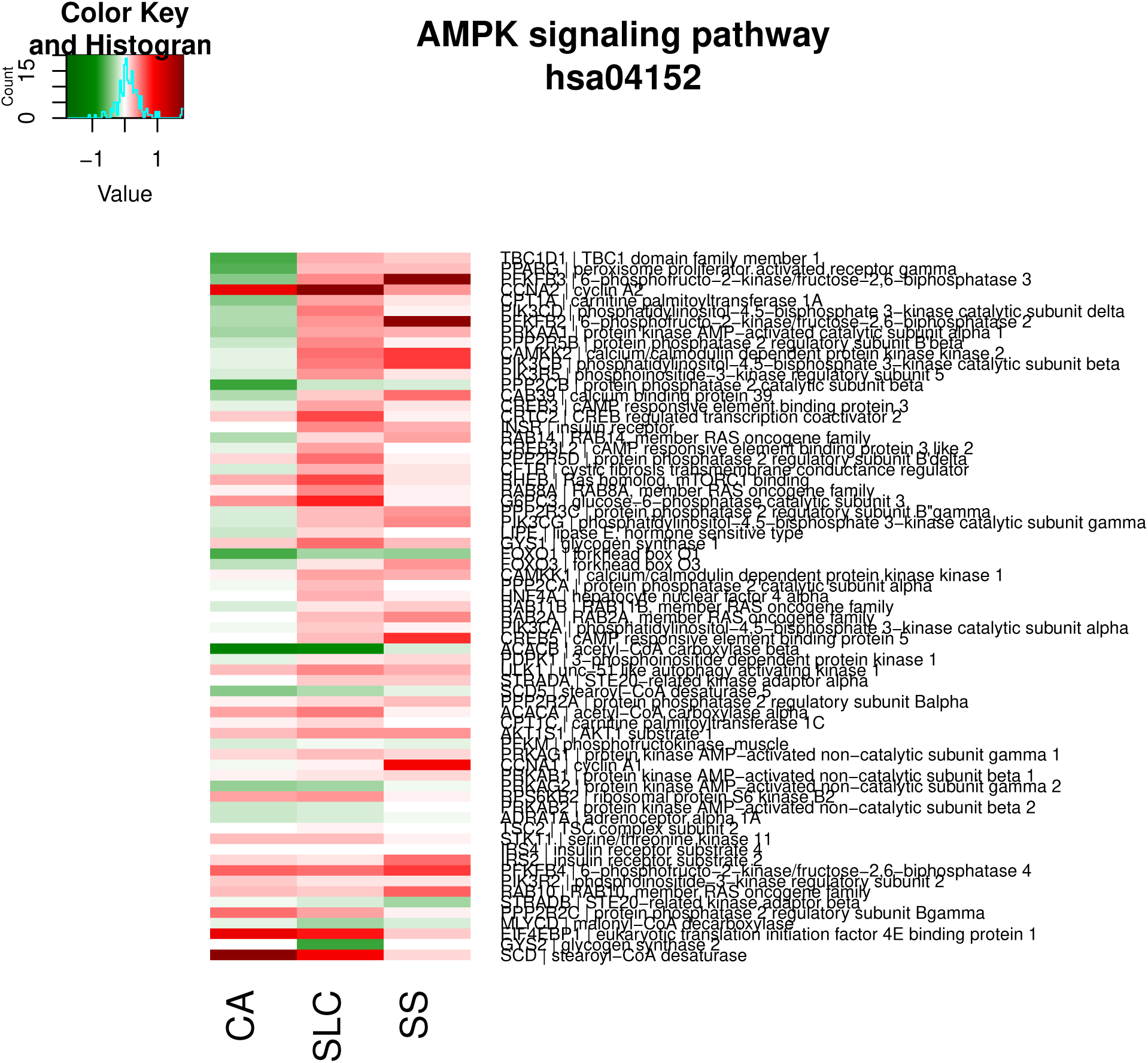

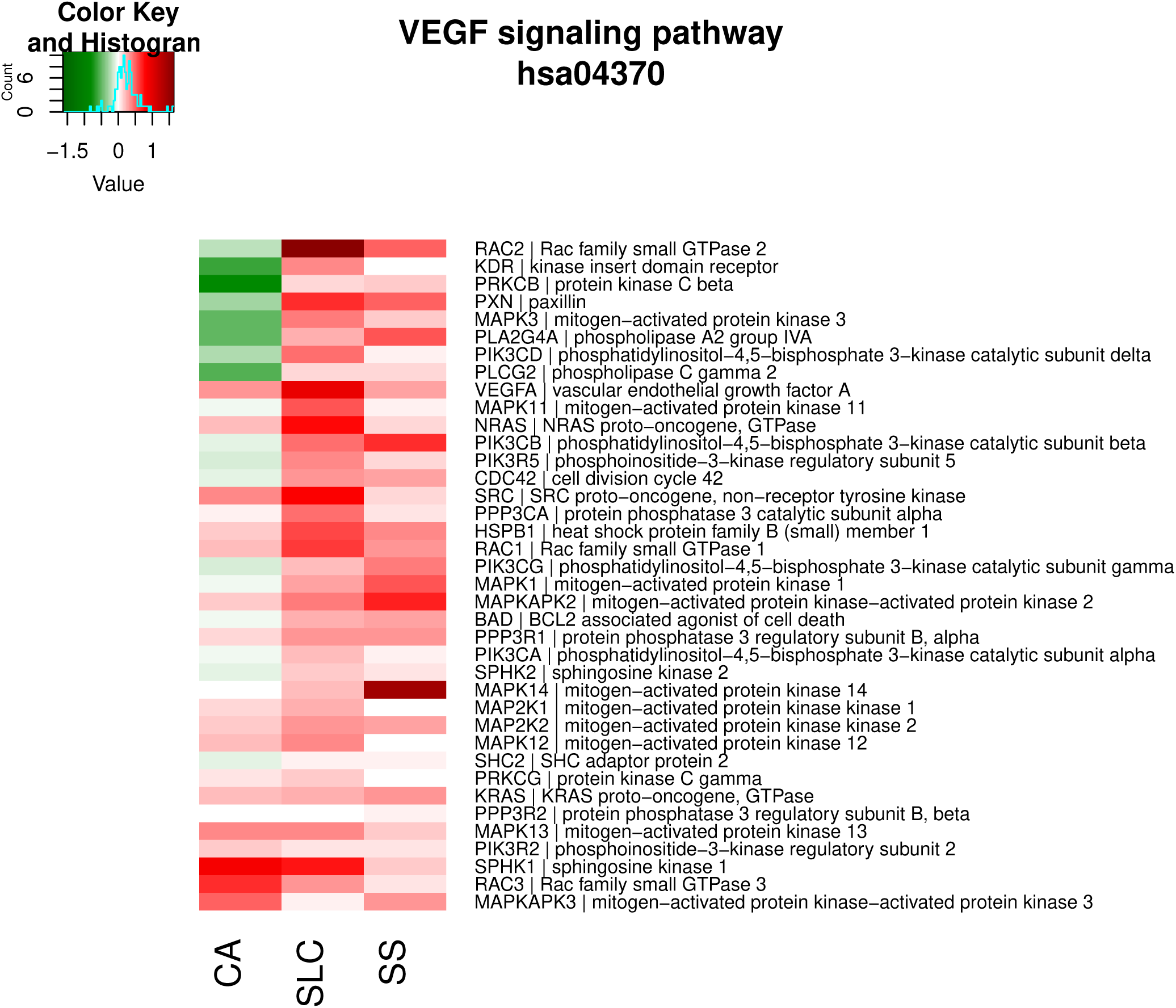

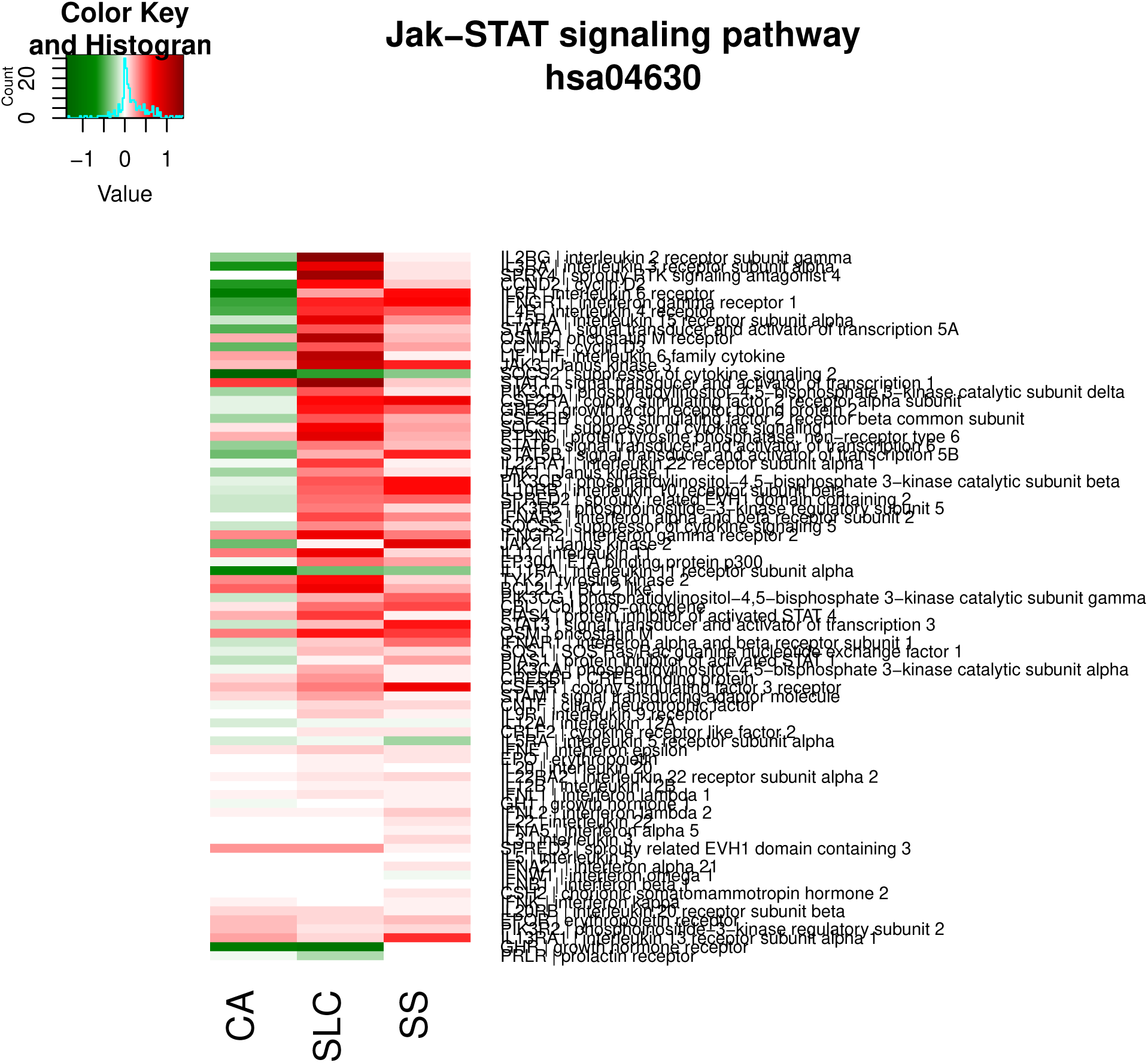

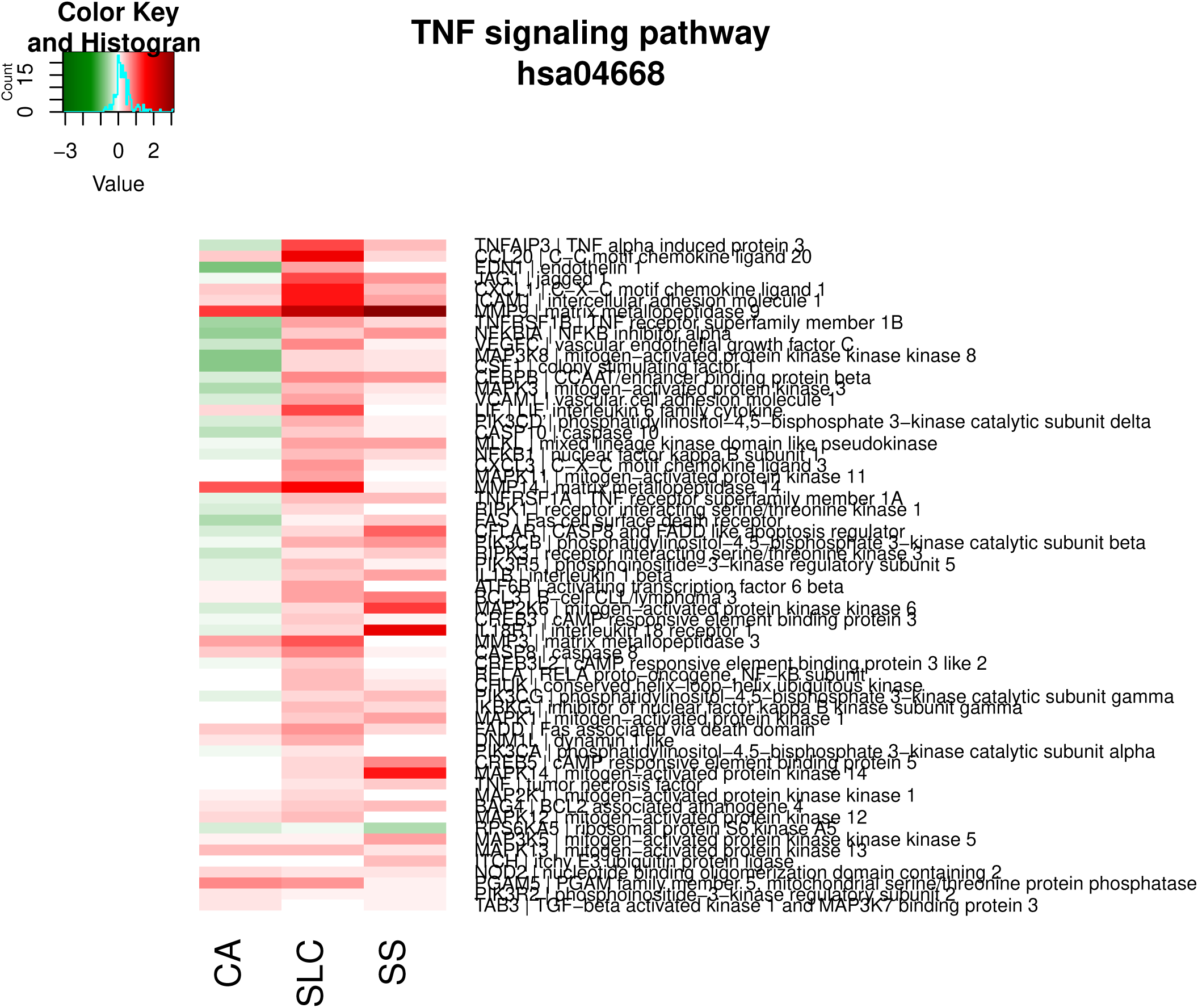

#### Transport and catabolism

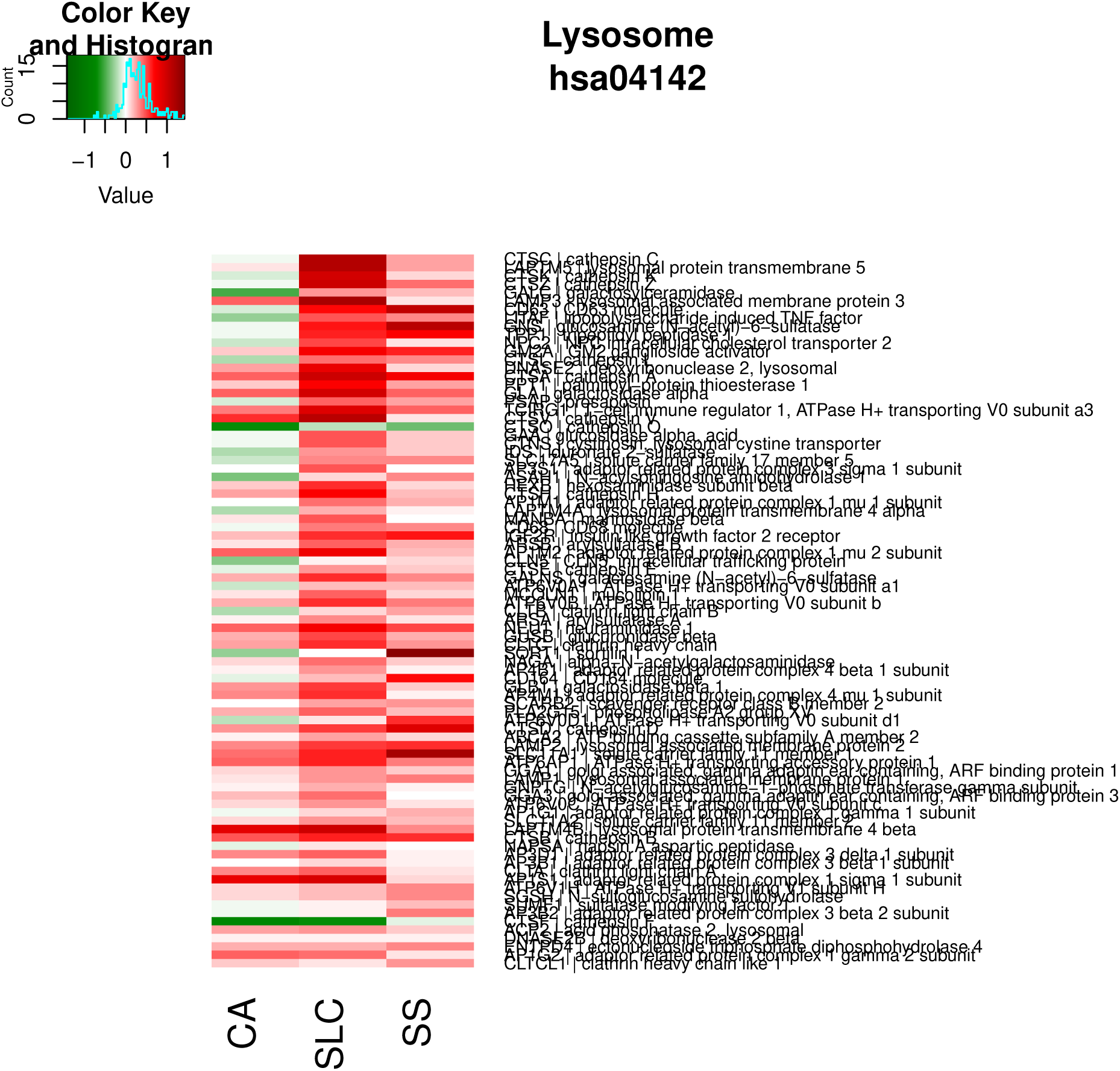

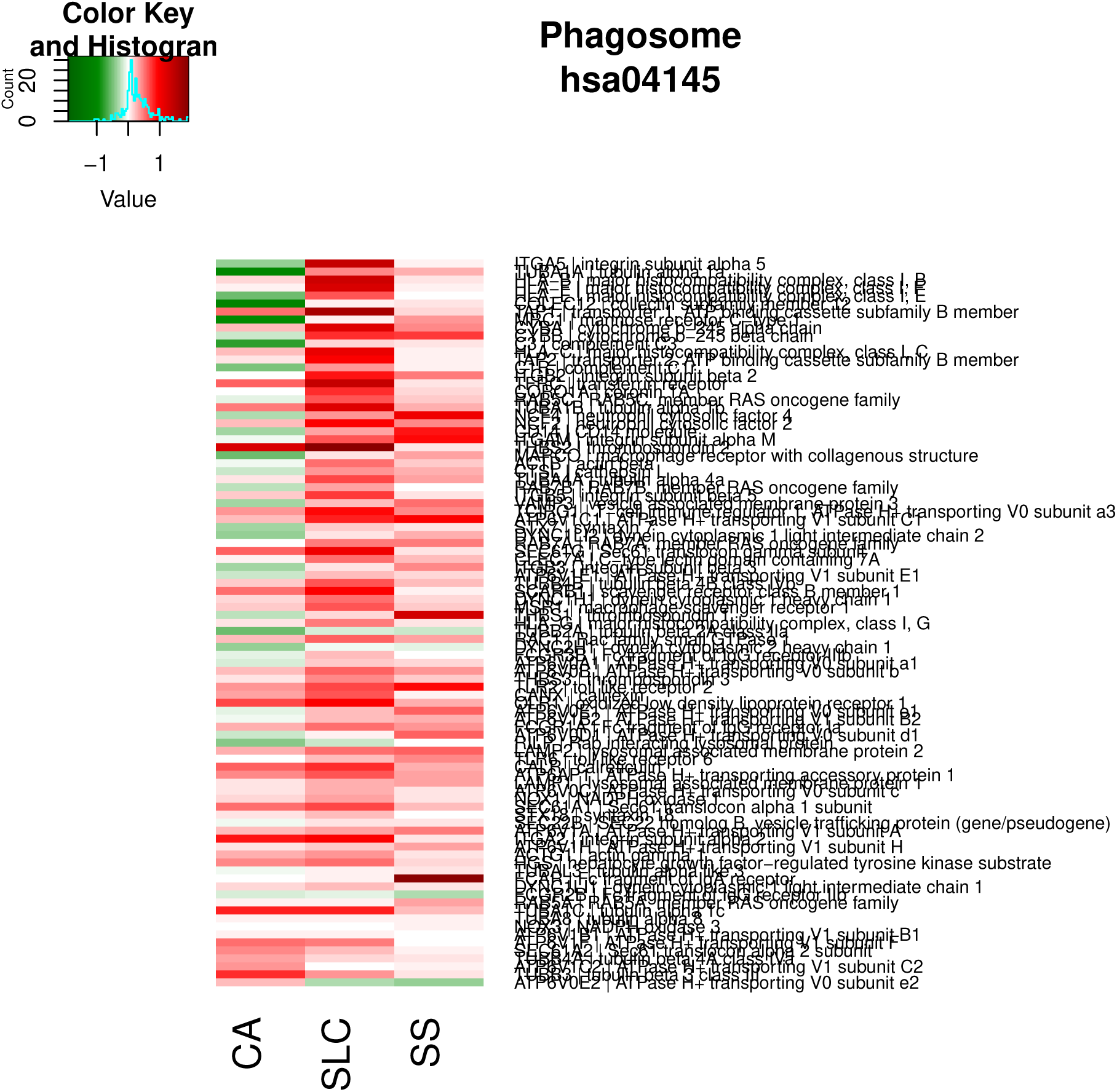

